# Novel Dissymmetric Ionizable Lipid-Assembled Lipid Nanoparticles for Delivery of Ferroptosis-Related siRNA in Diabetic Treatment

**DOI:** 10.64898/2026.08.26.747432

**Authors:** Han Zhang, Yuanyuan Liu, Fengyang He, Genping Xue, Yi Kang, Zepu Zhang, Jixin Ma, Junhai Xiao, Qingbin Meng

**Author notes:** Corresponding authors: Junhai Xiao - E-mail address; Qingbin Meng. These authors contributed to work equally (Han Zhang, Yuanyuan Liu, Fengyang He).

## Abstract

Small interfering RNA (siRNA) enables precise post-transcriptional gene silencing for refractory diseases, yet its clinical translation remains limited by the lack of safe and efficient delivery vectors. Inspired by the dissymmetric alkyl chain architecture of natural membrane phospholipids, we designed and synthesized 34 novel ionizable lipids with dissymmetric hydrophobic tails and formulated them into lipid nanoparticles (LNPs). Through systematic physicochemical and biological assessments, we established clear structure-activity relationships and identified two lead LNPs (O14-LNP, H18a-LNP) with superior endosomal escape capacity, enhanced *in vivo* gene silencing potency, and favorable biosafety relative to the clinical benchmark MC3-LNP. In both streptozotocin-induced and spontaneous db/db type 2 diabetes (T2D) mouse models, lead LNPs delivering ferroptosis-related siRNAs effectively ameliorated glucose and lipid metabolic disorders, restored islet function, and alleviated hepatic steatosis. This study not only lays a theoretical foundation for the rational design of novel ionizable lipids, but also validates the therapeutic potential of siRNA therapy targeting ferroptosis, providing a versatile delivery platform and targeted therapeutic strategy for the treatment of T2D.

## 1. Introduction

Nucleic acid drugs, with their precise gene sequence-based regulatory capabilities, have emerged as the third major class of drugs following small-molecule drugs and antibody therapeutics [1]. Among them, small interfering RNA (siRNA) can specifically degrade target mRNA through the RNA interference mechanism, enabling post-transcriptional silencing of disease-causing genes and providing a novel therapeutic strategy for intractable diseases such as cancer, genetic disorders, and metabolic diseases [2, 3]. However, naked siRNA has a large molecular weight and carries negative charges, making it difficult to spontaneously penetrate the lipid bilayer of cell membranes. Meanwhile, it is susceptible to rapid degradation by endogenous nucleases, resulting in an extremely short circulation half-life [1, 4, 5]. Therefore, the development of safe and efficient delivery vehicles has remained a critical bottleneck limiting the clinical translation of siRNA drugs. Lipid nanoparticles (LNPs) represent the most technologically mature and clinically successful non-viral nucleic acid delivery system to date, and have been validated in multiple approved siRNA drugs and mRNA vaccines [6–8]. In particular, ionizable lipids, as the core functional component of LNPs, are key determinants of *in vivo* delivery efficiency, endosomal escape capacity, and biosafety [9, 10].

The most widely used ionizable lipid in current clinical applications is represented by DLin-MC3-DMA (MC3), which serves as the key lipid component of patisiran, the first approved siRNA drug [11–13]. Despite its success, MC3 exhibits limitations such as insufficient endosomal escape efficiency and *in vivo* transfection activity, as well as poor delivery adaptability across different gene sequences, which cannot fully meet the growing demands of nucleic acid drug development [4, 6, 11–13]. Accordingly, developing novel ionizable lipids with superior delivery activity and better safety profiles remains a core research priority in the field of nucleic acid delivery. The molecular structure of ionizable lipids generally consists of three moieties: a hydrophilic headgroup, an intermediate linker, and hydrophobic tails. Among these, the protonation capacity of the headgroup, the chemical properties of the linker, and the chain length and unsaturation degree of the hydrophobic tails exert decisive effects on the delivery performance of LNPs [9, 14–16]. Notably, natural cell membrane phospholipids, such as sphingomyelin and phosphatidylcholine, commonly exhibit dissymmetric alkyl chain substitution. This dissymmetric architecture is essential for maintaining membrane bilayer fluidity, mediating membrane fusion, and promoting the formation of non-lamellar phases, such as the inverted hexagonal phase [17–20]. Inspired by this, we hypothesized that ionizable lipids with dissymmetric hydrophobic tails may be more favorable for driving the transition of lipid membranes from the lamellar phase to the inverted hexagonal phase in the acidic endosomal environment, thereby promoting endosomal membrane disruption and cytosolic release of siRNA and ultimately enhancing gene silencing efficiency. However, systematic studies on the structure-activity relationship (SAR) of ionizable lipids with dissymmetric hydrophobic tails remain scarce, and the regulatory mechanisms of their structural features on the physicochemical properties and *in vitro/in vivo* delivery activity of LNPs have not been fully elucidated.

Diabetes, particularly type 2 diabetes (T2D), is a chronic metabolic disease characterized by persistent hyperglycemia and lipid dysregulation, with its global prevalence continuously rising and posing a major public health challenge [21, 22]. Current clinical treatments primarily focus on symptomatic glycemic control, which fails to reverse disease progression from its pathological root cause. Long-term medication is also associated with adverse effects such as hypoglycemia, weight gain, and gastrointestinal reactions, underscoring the urgent need for novel therapeutic targets and intervention strategies [23, 24]. Recent studies have revealed that ferroptosis, an iron-dependent form of programmed cell death defined by the accumulation of lipid peroxidation, plays a critical role in the development and progression of pancreatic β-cell dysfunction, hepatic steatosis, and insulin resistance in T2D [25–28]. Therefore, targeted modulation of the ferroptosis pathway holds promise for delaying the progression of diabetes at its source and represents a clinically valuable therapeutic approach. Compared with small-molecule ferroptosis inhibitors, siRNA can precisely regulate key nodes of ferroptosis at the genetic level, with the advantages of high target specificity and long-lasting efficacy, providing a novel strategy for the targeted treatment of diabetes. In this study, three ferroptosis-related targets with well-defined functions and distinct regulatory mechanisms were selected for validation. ALOX12 encodes arachidonic acid 12-lipoxygenase, which directly catalyzes the oxidation of polyunsaturated fatty acids and acts as a key enzyme initiating lipid peroxidation [29, 30]. ACSL4 encodes acyl-CoA synthetase long-chain family member 4, which catalyzes the esterification of long-chain polyunsaturated fatty acids, providing substrates for lipid peroxidation [31, 32]. Kelch-like ECH-associated protein 1 (Keap1) is a negative regulatory protein of nuclear factor erythroid 2-related factor 2 (Nrf2), and silencing Keap1 activates the Nrf2 antioxidant pathway to indirectly inhibit ferroptosis [33, 34]. In addition, glutathione peroxidase 4 (GPX4) is a core intracellular defense molecule against ferroptosis that specifically reduces and clears lipid peroxides, so its expression level and activity directly determine cellular sensitivity to ferroptosis [35]. The above three pathways participate in ferroptosis regulation from the dimensions of pro-oxidation, substrate supply, and antioxidant defense, respectively. The selection of these targets not only allows for a comprehensive validation of the generality of ferroptosis intervention strategies but also enables simultaneous evaluation of the delivery adaptability of the LNP platform for different siRNA sequences.

Building on the above background, this study first designed and synthesized 34 novel ionizable lipids with dissymmetric hydrophobic tail structures. Using symmetric controls and MC3 as references, these lipids were formulated into LNPs to systematically investigate the effects of hydrophobic tail number, chain length, unsaturation degree, headgroup type, and linker chemical properties on the physicochemical properties, *in vitro* cellular uptake, gene silencing efficiency, and *in vivo* biodistribution of LNPs, to clarify their SAR. After safety screening, lead lipids were identified, and their cellular uptake and endosomal escape mechanisms were elucidated. Finally, siRNAs targeting different ferroptosis pathways were delivered in both streptozotocin (STZ)-induced and db/db T2D mouse models. Therapeutic efficacy was comprehensively evaluated across glucose/lipid metabolism, islet function, hepatic steatosis, and ferroptosis pathway modulation, confirming the delivery versatility of the vector and the therapeutic potential of ferroptosis-targeted nucleic acid therapy for T2D (Fig. 1). Besides developing novel ionizable lipids outperforming MC3, this study not only reveals the regulatory rules of dissymmetric hydrophobic tails on LNP delivery efficiency, providing experimental evidence for rational lipid design, but also confirms that ferroptosis-targeted siRNA therapy effectively ameliorates glucose/lipid metabolism disorders and organ damage in T2D, offering a novel target strategy and delivery platform for nucleic acid drug development against T2D.

**Fig. 1.**
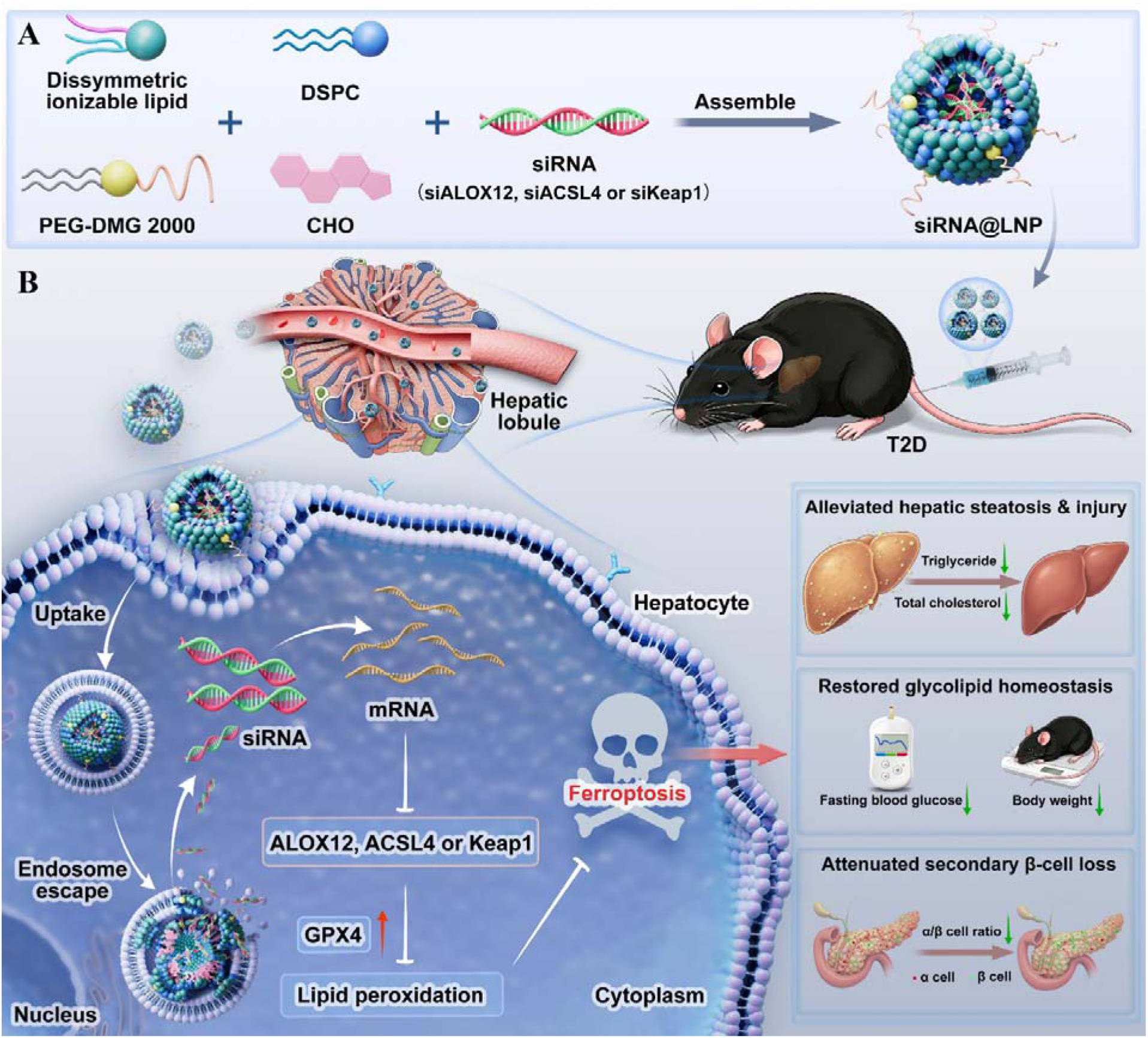
Schematic diagram depicting the formulation of siRNA@LNPs and their anti-ferroptosis mechanism against type 2 diabetes. (A) The preparation processes of siRNA-encapsulated LNPs composed of ionizable lipid, DSPC, cholesterol, DMG-PEG2000 and target siRNA. (B) Following endocytosis and endosomal escape, siRNA released from LNPs silences ACSL4, Keap1 or ALOX12 mRNA, restrains lipid peroxidation, upregulates GPX4, blocks ferroptosis, and ameliorates β-cell death, dyslipidemia, liver damage and hyperglycemia in type 2 diabetes.

## 2. Experimental Section

### 2.1 Materials

All chemicals and solvents were purchased from commercial suppliers and used directly, with solvents dried following standard laboratory procedures. Both ^1^H nuclear magnetic resonance (NMR) and ^13^C NMR spectra were acquired on a 600 MHz Bruker spectrometer (Switzerland). All samples were dissolved in CDCl_3_, with tetramethylsilane serving as the internal standard, and chemical shifts were reported in ppm. NMR data were analyzed and processed using MestReNova software (Version 6.1.0, Mestrelab Research, Spain). The molecular weights of all final synthetic products were measured on a Bruker ultrafleXtreme matrix-assisted laser desorption/ionization (MALDI) TOF/TOF or electrospray ionization (ESI) mass spectrometer (Switzerland).

Human bronchial epithelial cells (BEAS-2B), mouse macrophages (RAW 264.7), human umbilical vein endothelial cells (HUVEC), human hepatocellular carcinoma cells (HepG2) and HepG2 cells stably expressing firefly luciferase (HepG2-Luc) were all obtained from the Peking Union Cell Bank, Beijing, China. RPMI-1640 medium, DMEM medium, fetal bovine serum (FBS), penicillin-streptomycin solution, PBS buffer, trypsin, Lipofectamine 2000 (Lipo 2000) and Ribogreen RNA assay kit were purchased from Thermo Fisher Scientific (USA). Hoechst 33342 live cell staining solution, Lyso-Tracker Red, C11-BODIPY 581/591 probe and 2% phosphotungstic acid staining solution, as well as sodium citrate buffer (pH = 4.0), were supplied by Beyotime Biotechnology, Shanghai, China. CPZ, amiloride, MβCD and paraformaldehyde were acquired from Sigma-Aldrich (Merck, Shanghai, China). Firefly luciferase reporter assay kits were obtained from Yeasen Biotechnology Co., Ltd. Cell Counting Kit-8 (CCK-8) was purchased from Dojindo Laboratories, Japan. In addition, Cy5-negative control siRNA (siNC), siLuc, siKeap1 and siALOX were provided by GenePharma Co., Ltd., Suzhou, China.

### 2.2 Synthesis of Compounds

The synthetic routes, detailed procedures, and structural characterization data (^1^H NMR, ^13^C NMR and mass spectrometry) of all 34 novel ionizable lipids are provided in the Supporting Information.

### 2.3 Formulation of LNPs

The newly synthesized ionizable lipids or positive control lipid MC3 were blended with cholesterol (CHO), 1,2-dioctadecanoyl-sn-glycero-3-phosphocholine (DSPC) and 1,2-dimyristoyl-rac-glycero-3-methoxypolyethylene glycol-2000 (DMG-PEG2000) at a molar ratio of 50:38.5:10:1.5, and fully dissolved in ethanol. Under continuous stirring, the resultant lipid solution was added into three volumes of citrate buffer (0.05 M, pH 4.0), followed by stirring for 2-3 min to form blank LNPs. Subsequently, siRNA dissolved in citrate buffer containing 25% (v/v) ethanol was introduced into the LNP mixture at 1:15 (wt/wt) siRNA/total lipids, and the system was incubated at 37 for 30 min to achieve efficient siRNA encapsulation. The obtained LNP suspension was dialyzed against PBS for 2 h using a dialysis bag with a molecular weight cutoff of 3500 Da to remove residual citrate and unencapsulated siRNA. The final siRNA@LNPs were stored at 4 for subsequent use.

### 2.4 LNPs characterization

The particle size, polydispersity index (PDI), and zeta potential of LNPs before and after siRNA encapsulation were measured using a Brookhaven NanoBrook ZetaPALS Potential Analyzer (Brookhaven Instruments Corporation, Holtsville, NY, USA) with a fixed scattering angle of 90°. LNP suspensions were diluted to a final concentration of 0.02 mg/mL with deionized water for relevant characterization, and each sample was tested in triplicate.

For transmission electron microscopy (TEM) observation, a drop of LNP suspension (1 mg/mL) was placed onto a 300-mesh carbon-coated copper grid and allowed to dry completely overnight. The grid was subsequently stained with 2% phosphotungstic acid solution for negative staining. Following 4 min of staining, excess liquid was carefully blotted with filter paper, and the sample was air-dried at ambient temperature. The morphological features of LNPs were visualized by TEM (Hitachi, HT7650, Tokyo, Japan). Negatively stained LNPs displayed a bright and transparent morphology against a dark background.

The encapsulated siRNA concentration in LNPs was quantified with the Quant-iT RiboGreen RNA Assay Kit according to the manufacturer’s instructions. In brief, LNPs and RNA standards were diluted with Tris-EDTA (TE) buffer and dispensed into a 96-well plate. Meanwhile, LNPs were separately incubated in TE buffer containing 2% Triton X-100 for 30 min to fully release encapsulated nucleic acids. Subsequently, freshly prepared RiboGreen working solution was added to each well. Fluorescence intensity was measured *via* a microplate reader (SpectraMax® M5, Molecular Devices, CA, USA) with excitation at 490 nm and emission at 520 nm. The siRNA concentration was calculated from the standard curve, and the encapsulation efficiency (E.E.) was computed using the following formula: E.E. (%) = [(total siRNA - free siRNA)/total siRNA] × 100%.

### 2.5 Dissociation constant (p*K*_a_) determination of LNPs

To evaluate the apparent p*K*_a_ of LNPs, LNP suspensions were diluted to 30 μM using a buffer system composed of 150 mM sodium chloride, 20 mM sodium dihydrogen phosphate, 20 mM ammonium acetate, and 25 mM ammonium citrate. The buffer pH was finely adjusted by acid-base titration to cover a range of 2.0 - 12.0 with a step interval of 0.5 pH units. 2-(*P*-toluidinyl)naphthalene-6-sulfonic acid (TNS) was supplemented into each LNP sample to yield a final concentration of 6 μM. Fluorescence signals were measured on a microplate reader at an excitation wavelength of 322 nm and an emission wavelength of 431 nm. The pH value at half-maximum fluorescence intensity was determined as the apparent p*K*_a_ of LNPs. All measurements were conducted in triplicate.

### 2.6 Cell culture

BEAS-2B, RAW 264.7, HepG2 and HepG2-Luc cells were cultured in DMEM plus 10% FBS and 1% penicillin-streptomycin solution. HUVEC cells were cultured in RPMI-1640 medium plus 10% FBS and 1% penicillin-streptomycin solution. All cells were cultured at 37 °C under a humidified atmosphere containing 5% CO_2_.

### 2.7 Cellular uptake

Flow cytometry was used to evaluate the cellular uptake efficiency of LNPs. HepG2 cells were seeded in 12-well plates at a density of 1□×□10^5^ cells per well and cultured for 24 h. The cells were then treated with Cy5-labeled siRNA (Cy5-siRNA)@LNPs. After 4 h of incubation, the cells were washed three times with PBS buffer, digested with 0.25% trypsin, and resuspended in 350 μL of 2% paraformaldehyde solution. Cellular fluorescence signals were detected using an Accuri C6 plus flow cytometer (BD Biosciences, San Jose, CA, USA). At least 10,000 valid cells were collected for each sample, and all experimental data were analyzed using FlowJo V10 software.

### 2.8 *In vitro* siRNA delivery

HepG2-Luc cells were seeded into 96-well plates at a density of 1□×□10^4^ cells per well and incubated for 24 h. The cells were then treated with various siLuc@LNPs at a final siLuc concentration of 50 nM for 4 h. After incubation, the supernatant was discarded, and 100 μL of fresh antibiotic-free medium was added to each well. The cells were further cultured for 20 h in an incubator. Subsequently, cell lysis buffer was added to fully lyse the cells, and intracellular firefly luciferase activity was measured using a luciferase assay system.

### 2.9 Animal studies

All animal experimental procedures were performed in accordance with the guidelines for the care and use of laboratory animals issued by the Academy of Military Medical Sciences (Approval Number: IACUC-2025−003B). C57BL/6 mice and ICR mice (6-8 weeks old, weighing approximately 20 g) were purchased from SPF Biotechnology Co., Ltd (Beijing). db/db mice and wild-type m/m mice were obtained from Gempharmatech Co., Ltd. (Jiangsu, China). All mice were housed under a controlled environment at a constant temperature of 25 °C with a 12 h light/12 h dark cycle, and free access to food and water was provided throughout the experiment.

### 2.10 *In vivo* biodistribution

C57BL/6 mice were intravenously injected *via* the tail vein with PBS or Cy5-siRNA@LNPs at a dose of 1 mg/kg. At 4 h post-injection, *in vivo* fluorescence imaging was performed. The mice were then euthanized at the same time point, and major organs were harvested for *ex vivo* imaging analysis. The average radiant efficiency [p/s/cm^²^/sr] / [µW/cm^²^] was quantified and analyzed using Living Image software.

### 2.11 Cell viability

Cell viability was assessed using the CCK-8 assay. Briefly, BEAS-2B cells were seeded into 96-well plates at a density of 1□×□10^4^ cells per well and incubated at 37 °C for 24 h. The cells were then treated with LNPs containing ionizable lipids at final concentrations of 10, 20, 40, 80, 160, and 320 μM, followed by another 24 h of incubation. Afterwards, the culture medium was replaced with fresh medium supplemented with 10% CCK-8 solution, and the cells were incubated at 37 °C for 1 h. The absorbance value at 450 nm was measured using a microplate reader. Cell viability was calculated according to the following formula: (experimental group OD − blank group OD mean) / (control group OD mean − blank group OD mean)□×□100%. Each group had three replicate wells. The above procedures were also applied to RAW 264.7, HepG2, and HUVEC cells for cytotoxicity evaluation.

### 2.12 Hemolysis test

Fresh rabbit blood (1 mL) was centrifuged at 1000 rpm for 5 min under low-temperature conditions, and the supernatant was discarded. The precipitate was washed three times with PBS, followed by resuspension in 20 mL of PBS to prepare a 5% (v/v) red blood cell (RBC) suspension. Subsequently, 1 mL of the prepared RBC suspension was incubated with PBS, 1% Triton X-100, or LNPs at final concentrations of 5, 10, 20, 40, 80, and 160 μM. All samples were incubated at 37 °C for 2 h and 16 h, respectively. After incubation, the samples were centrifuged at 1000 rpm for 5 min in a low-temperature environment, and macroscopic morphological changes were photographed and observed. The absorbance of the supernatant in each group was measured at 405 nm using a microplate reader to quantify hemoglobin release from erythrocytes. The hemolysis rate was calculated according to the following formula: Hemolysis rate (%) = [(OD of samples - OD of PBS) / (OD of Triton X-100 - OD of PBS)] × 100%.

### 2.13 *In vivo* biosafety evaluation

Thirty male ICR mice were randomly divided into six groups: PBS group, siNC@O18-LNP group (3 mg/kg), siNC@H18a-LNP group (3 mg/kg), siNC@O14-LNP group (3 mg/kg), siNC@K6-LNP group (3 mg/kg), and siNC@MC3-LNP group (3 mg/kg). All treatments were administered *via* tail vein injection. The mice were euthanized after 7 days of administration. Blood samples and major organs, including the heart, liver, spleen, lungs, and kidneys, were collected for routine blood analysis, serum biochemical detection, and histological examination. Histological sections were scanned and observed using the VS200 digital scanning system (Olympus Inc., Japan).

### 2.14 Subcellular localization and mechanism of internalization

HepG2 cells were seeded in 35 mm culture dishes at a density of 1□×□10^5^ cells per well and incubated for 24 h. To explore the cellular internalization mechanism of LNPs, adherent cells were pretreated with chlorpromazine (CPZ, 10 μg/mL), amiloride (am, 50 μM), methyl-β-cyclodextrin (MβCD, 5 mM), or fresh DMEM for 30 min, respectively. Subsequently, the medium was replaced with antibiotic-free DMEM, and the cells were incubated with Cy5-siRNA@LNPs at a final Cy-siNC concentration of 50 nM. After 4 h of incubation, the cells were rinsed with PBS. Then, cells were stained with LysoTracker Red (50 nM) and Hoechst 33342 (10 μg/mL) at 37 °C for 20 min separately. Following staining, the staining solution was discarded, and the cells were washed three times with PBS, prior to the addition of 1 mL PBS for imaging observation. Cellular fluorescence imaging was performed using a laser confocal microscope (Dragonfly 200, Andor, UK). Colocalization analysis was conducted using ImageJ software.

### 2.15 Half-maximal effective dose (ED_50_) determination

Sixty-four C57BL/6 mice were randomly assigned into 16 groups (four mice per group) and intravenously administered PBS, siALOX@H18a-LNP, siALOX@O14-LNP, or siALOX@MC3-LNP *via* tail vein injection at doses ranging from 0.03 to 0.5 mg/kg. All mice were euthanized 48 h post-administration, and liver tissues were harvested. The mRNA expression levels of ALOX12 in liver tissues were quantified *via* quantitative polymerase chain reaction (qPCR) analysis.

### 2.16 Lipid peroxidation (LPO) determination

Intracellular LPO levels were determined using the C11-BODIPY (581/591) probe according to the manufacturer’s instructions. Briefly, RAW 264.7 cells were seeded in 35 mm culture dishes at a density of 1□×□10^5^ cells per dish and cultured for 24 h. The cells were first stimulated with 30 mM glucose solution for 4 h, followed by treatment with different LNPs and further incubation at 37 °C for 4 h. Subsequently, the cells were rinsed with PBS. After that, cells were stained with 1 μM Hoechst 33342 for 20 min and 5 μM C11-BODIPY (581/591) for 1 h at 37 °C. Fluorescence imaging was performed *via* laser confocal microscopy. All quantitative analyses were processed using ImageJ software.

### 2.17 Treatment of STZ-induced diabetic mice

Five male C57BL/6 mice weighing 18 - 22 g were fed with standard chow and assigned as the blank control group (C57BL/6 PBS group). Another 20 male C57BL/6 mice with the same weight range were supplied with a high-fat diet for 50 consecutive days. After 12 h of fasting, mice received an intraperitoneal injection of STZ at a dose of 100 mg/kg, and their fasting blood glucose was monitored daily for 7 consecutive days. Mice with fasting blood glucose levels higher than 11.1 mM were considered successfully modeled and included in subsequent experiments.

The modeled mice were randomly divided into four groups (n = 5), namely the STZ PBS group, STZ siALOX@H18a-LNP group, STZ siALOX@O14-LNP group, and STZ siALOX@MC3-LNP group. All mice were administered *via* tail vein injection at a dosage of 0.25 mg/kg on days 1, 8, 15, and 22 after grouping. During the whole experimental period, the STZ model mice were continuously fed a high-fat diet, while the control mice were maintained on standard chow.

Following each administration, mice were fasted for 6 h on the 6th day, and their body weight and fasting blood glucose (collected from tail veins) were measured on the 7th day. All animals were euthanized on day 28. Whole blood and serum samples were collected for routine blood examination and serum biochemical analysis. Major organs including the heart, liver, spleen, lung and kidney were isolated and fixed in 4% paraformaldehyde for histological evaluation. Partial liver tissues were frozen and stored for subsequent mRNA detection. Oil Red O staining was performed to evaluate lipid accumulation in the liver. In addition, pancreatic tissues were harvested, and the expression levels of insulin and glucagon in pancreatic islets were detected by immunofluorescence staining.

### 2.18 Treatment of db/db mice

Twenty-five male db/db spontaneous diabetic mice were randomly divided into five groups (n=5 per group): db/db PBS group, db/db siACSL4-L group (0.25 mg/kg), db/db siACSL4-H group (0.5 mg/kg), db/db siKeap1-L group (0.25 mg/kg), and db/db siKeap1-H group (0.5 mg/kg). Another five age-matched littermate m/m mice were set as the blank control group (m/m PBS). The siACSL4 and siKeap1 used in this study were target-specific siRNAs encapsulated by H18a-LNPs.

All mice received tail vein injection once a week for 4 consecutive weeks. Body weight was recorded every 7 days throughout the experiment. On the 5th day after each administration, mice were fasted for 6 h, and blood glucose levels were measured using tail venous blood samples. All animals were euthanized on the second day after the final treatment. Whole blood and serum were collected for routine blood examination and serum biochemical analysis. Partial liver tissues were isolated and frozen for subsequent detection of target gene mRNA expression. In addition, major organs including the heart, liver, spleen, lung and kidney were harvested and fixed in 4% paraformaldehyde for histopathological analysis.

### 2.19 Statistical Analysis

All data are presented as the mean ± standard deviation (SD) from no fewer than three independent experiments. Statistical evaluations were conducted via one-way or two-way analysis of variance (ANOVA), followed by Tukey’s multiple comparison test. A difference of *P* < 0.05 was defined as statistically significant. All statistical analyses and graph plotting were performed using Origin Pro 2021 (Origin Lab, MA, USA).

## 3. Results and discussion

### 3.1 Design and Synthesis of Novel Ionizable Lipids with Dissymmetric Hydrophobic Tails

Herein, inspired by naturally occurring phospholipids with dissymmetric hydrophobic tails in cell membranes (e.g., sphingomyelins) [20], we designed a series of ionizable lipids featuring dissymmetric hydrophobic moieties to develop LNP delivery systems with enhanced siRNA delivery efficacy. All designed ionizable lipids consist of three modular components: (1) a hydrophilic head group, including propane-1,3-diamine (D3), 3-aminopropan-1-ol (E3), N^1^-(3-aminopropyl)-N^1^-methylpropane-1,3-diamine (K6), and 3,3’-(piperazine-1,4-diyl)bis(propan-1-amine) (P6); (2) a linker moiety, represented by tertiary amine, 2-hydroxyalkylamine, ester group, or amide functional group (denoted by A, C, O, and H, respectively); and (3) alkyl tails of varying lengths, denoted by the number of carbon atoms following the linker designation. This modular design allowed us to systematically investigate the effects of hydrophilic head groups, linker chemistries, and hydrophobic tail structures on the endosomal escape capacity and transfection efficiency of the resulting LNPs.

Specifically, we designed and synthesized 34 novel dissymmetric ionizable lipids across 10 distinct series (Fig. 2), including the dissymmetric four-tail series (4 tail), dissymmetric ester-linked three-tailed series (E3-H8), dissymmetric amide-linked three-tailed series (D3-H8), dissymmetric ester amine-linked three-tailed series (O18-D3, O16-D3, O14-D3, O12-D3), K6 head group series (O12-K6), P6 head group series (O12-P6), and unsaturated hydrophobic tail series (O12-D3n). The design focused primarily on modulating the type, length, and degree of unsaturation of the hydrophobic tails. In parallel, two symmetric ionizable lipids (2C12-D3 and 2O12-D3) with identical hydrophobic tails were synthesized as controls (Fig. 2).

**Fig. 2.**
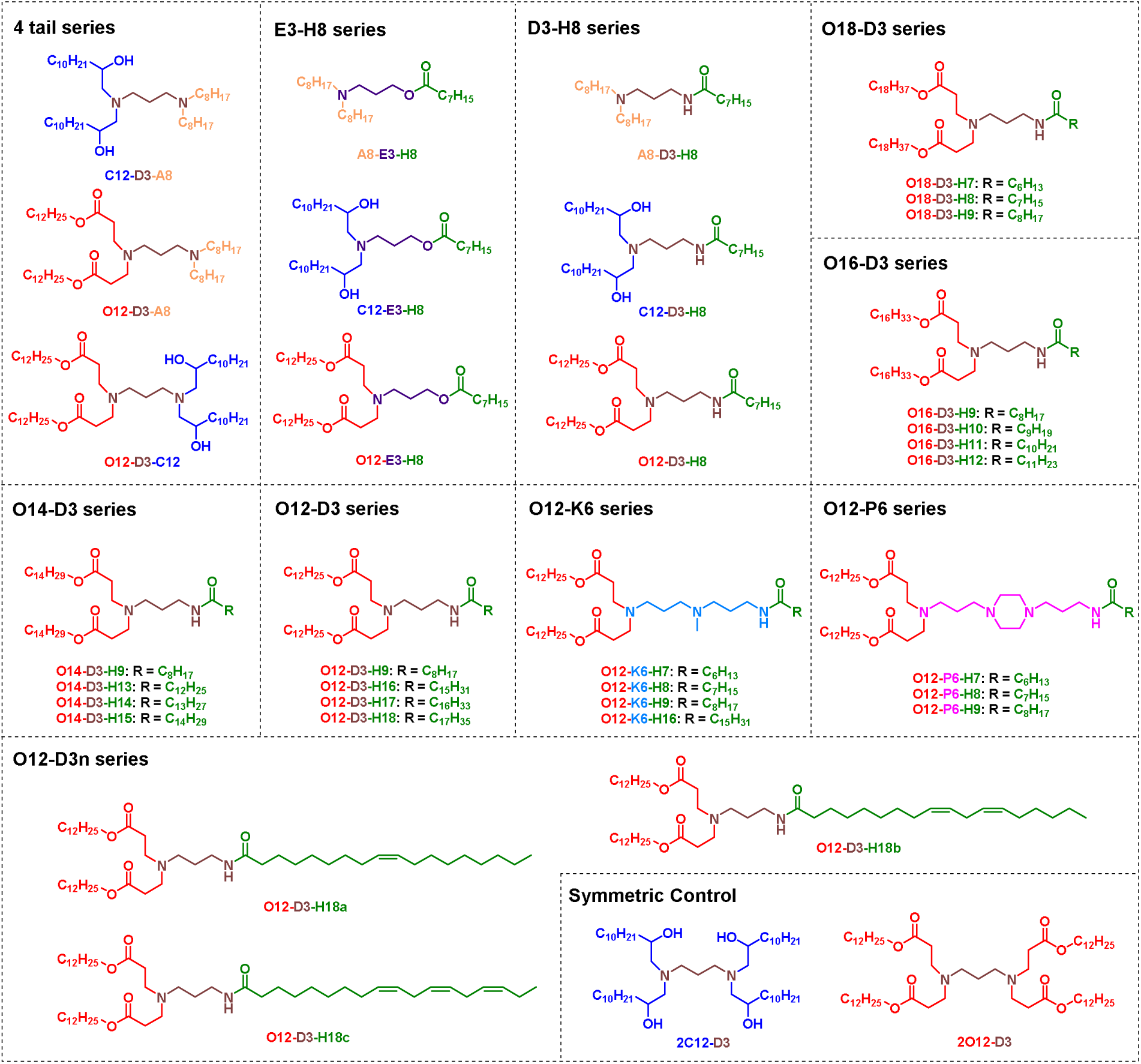
Chemical structures of 34 novel dissymmetric ionizable lipids and 2 symmetric control ionizable lipids.

Detailed synthetic procedures for all compounds are provided in the Supporting Information. Briefly, various key intermediates (e.g., C12-D3, O12-D3, A8-E3) were first prepared *via* alkylation, amidation, epoxide ring-opening, and tert-butyloxycarbonyl (Boc) protection/deprotection reactions. Synthetic protocols for several intermediates were adapted from our previously reported procedures [36]. Subsequently, the corresponding intermediates were reacted with alkyl halides of varying chain lengths (e.g., 1-bromooctane), carboxylic acids/acid chlorides (e.g., octanoic acid, palmitic acid), or unsaturated fatty acid derivatives under basic conditions, using sodium carbonate (Na_2_CO_3_), N,N-diisopropylethylamine (DIPEA), or triethylamine (TEA) as bases and O-(7-azabenzotriazol-1-yl)-N,N,N’,N’-tetramethyluronium hexafluorophosphate (HATU) or 1-ethyl-3-(3-dimethylaminopropyl) carbodiimide (EDCI) as coupling agents. Hydrophobic tails or hydrophilic head groups were introduced *via* alkylation, amidation, or esterification reactions. All synthesized compounds were purified by silica gel column chromatography and fully characterized *via* ^1^H NMR spectroscopy, ^13^C NMR spectroscopy, and MALDI or ESI mass spectrometry. The corresponding NMR spectra and mass spectra are provided in the Supporting Information.

### 3.2 Preparation and Characterization of LNPs

LNPs were prepared *via* the ethanol injection method. The lipid formulation consisted of ionizable lipid, DSPC, CHO, and DMG-PEG2000 at a molar ratio of 50:10:38.5:1.5 (Fig. 1A) [13, 36]. The particle size before and after encapsulation of siNC, PDI and zeta potential of siNC-loaded LNPs were systematically characterized (Fig. 3). The particle size of blank LNPs ranged from 84 to 163 nm, with the majority concentrated in the optimal range of 90-120 nm for nucleic acid delivery (Fig. 3A). Following siNC encapsulation, the particle size of all LNPs increased to varying degrees, with an average increase of approximately 45 nm, and most samples exhibited a size increment of 20-60 nm (Fig. S1). All LNPs displayed PDI values below 0.25, and more than 90% of the formulations showed PDI values lower than 0.2, indicating good monodispersity and homogeneity of the prepared nanoparticles (Fig. 3A). TEM observations revealed that siNC-loaded LNPs exhibited a spherical morphology with smooth and clear edges and no obvious aggregation, with a particle size of approximately 100 nm, which was consistent with the hydrodynamic particle size results determined by dynamic light scattering (Fig. 3B).

**Fig. 3.**
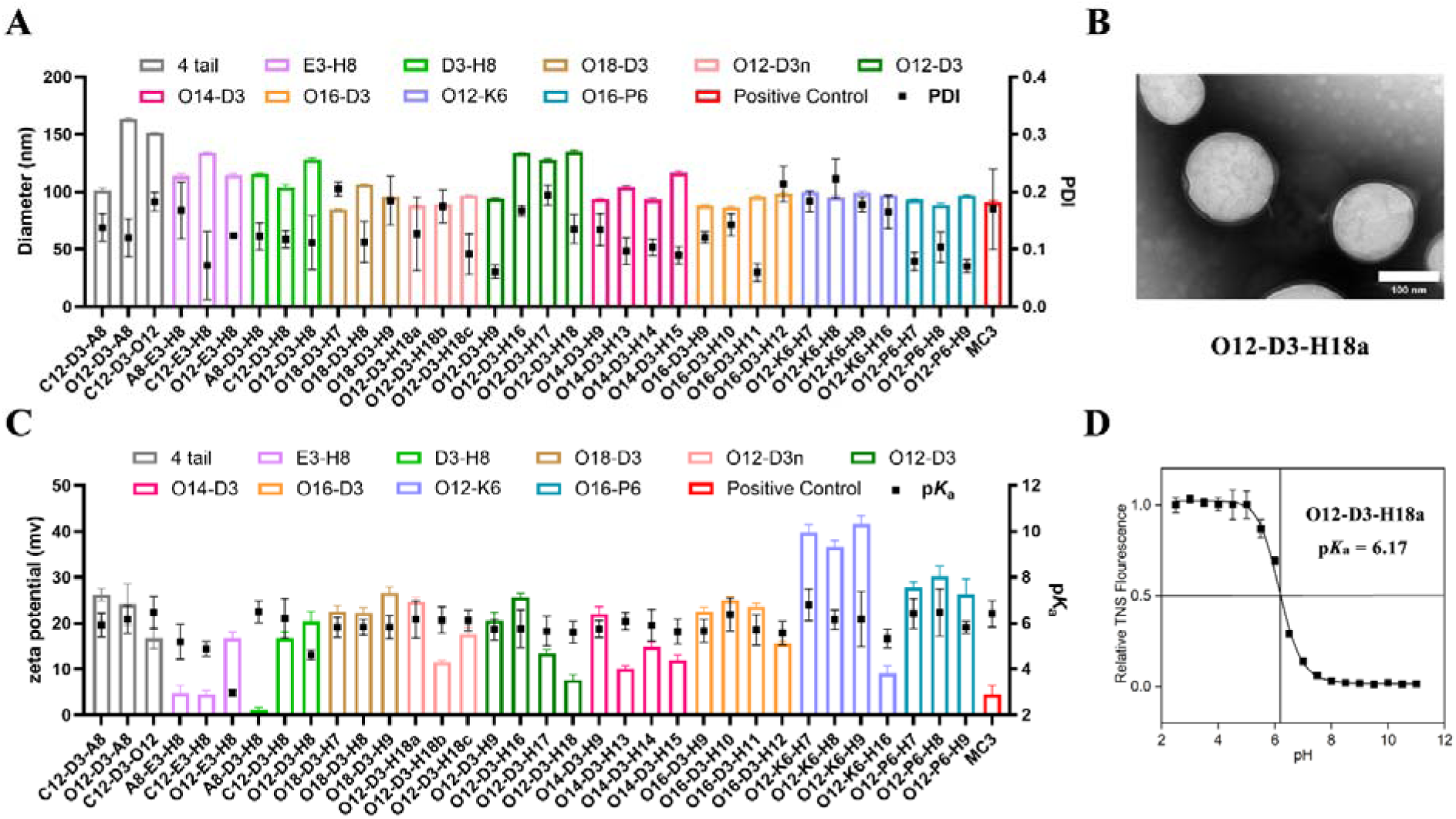
Physicochemical characterization of LNPs formulated with 34 dissymmetric ionizable lipids. (A) Particle size of blank LNPs (left Y-axis) and polydispersity index (PDI) of siNC-loaded LNPs (right Y-axis). (B) Representative transmission electron microscopy (TEM) image of siNC-loaded LNPs formulated with the ionizable lipid O12-D3-H18a. Scale bar = 100 nm. (C) Zeta potential (left Y-axis) and apparent p*K*_a_ values (right Y-axis) of LNPs. (D) Representative p*K*_a_ fitting curve for LNPs formulated with O12-D3-H18a. Data are presented as mean ± SD (n=3).

The zeta potentials of the LNPs ranged from 1.2 to 41.7 mV (Fig. 3C). The type of hydrophilic head group was the core factor determining zeta potential. The K6 and P6 series, which contain multiple amino groups (3-4 ionizable amines), exhibited higher surface positive charge density and consequently higher zeta potential values than other series. While elevated surface charge can inhibit particle aggregation and enhance LNP stability through electrostatic repulsion, excessively high positive charge may lead to non-specific adsorption of plasma proteins, potentially reducing circulation half-life and increasing systemic toxicity or hemolysis. In contrast, the E3 series, featuring a single ionizable amino group, displayed the lowest zeta potential, consistent with their structural characteristics. The D3 series, with two amino groups, exhibited intermediate zeta potential values. Additionally, the length, degree of unsaturation, and linkage type of the hydrophobic tails also influenced the zeta potential.

LNPs with high E.E. can effectively protect nucleic acids from nuclease degradation to enhance *in vivo* delivery efficiency, and an E.E. of more than 80% is generally required for nucleic acid delivery [9]. In this study, the E.E. of siNC-loaded LNPs was determined using the RiboGreen fluorescence assay. The results showed that the E.E. of LNPs formed by the 34 synthesized dissymmetric ionizable lipids ranged from 78% to 99%, with the vast majority exceeding 80%, meeting the basic requirements for siRNA delivery (Fig. S2). The p*K*_a_ is a critical factor affecting nucleic acid delivery and endosomal escape. LNPs with an appropriate p*K*_a_ (6.0-6.9) remain electrically neutral and stable at physiological pH (7.4), and become ionized in the acidic endosomal environment (pH 5.0-6.0), thereby promoting membrane fusion and enhancing endosomal escape [11, 37–39]. In this study, the apparent p*K*_a_ values of the prepared LNPs were measured using the TNS fluorescence method. The results showed that the apparent p*K*_a_ values of the prepared LNPs ranged from 2.95 to 6.80, among which 21 formulations had p*K*_a_ values in the range of 5.8-6.9 (Fig. 3C and 3D), theoretically possessing good endosomal escape potential and nucleic acid delivery prospects [37–39]. Compared with amide-containing lipids, ester-containing lipids exhibited lower p*K*_a_ values, which may be attributed to the electron-withdrawing effect of the ester group which reduces the basicity of the amino group and thus leads to a lower p*K*_a_ value [40,41]. The length and unsaturation degree of the hydrophobic tails exerted relatively minor effects on p*K*_a_ values. However, p*K*_a_ first rose and then declined with the elongation of hydrophobic tail chains. Moreover, an increase in the number of unsaturated bonds within the hydrophobic tails led to a gradual decrease in p*K*_a_ values.

Collectively, the LNPs prepared from 34 dissymmetric ionizable lipids exhibited favorable physicochemical properties: the majority of formulations demonstrated particle sizes concentrated in the range of 90-120 nm, PDI values below 0.2, positive zeta potential values, and E.E. exceeding 80%. Notably, 21 of the LNPs exhibited apparent p*K*_a_ values within the optimal range of 5.8-6.9. These findings indicate that these LNPs are well-suited for siRNA delivery, laying a solid foundation for subsequent *in vitro* and *in vivo* experiments.

### 3.3 Dissymmetric Hydrophobic-Tailed Ionizable Lipids Endow LNPs with Superior *In Vitro* Cellular Activities and *In Vivo* Delivery Capacities Compared with MC3 and Symmetric Lipids

To systematically evaluate the siRNA delivery potential of LNPs formulated from the above 34 newly synthesized dissymmetric hydrophobic-tailed ionizable lipids, we performed multidimensional biological assessments at cellular and animal levels. Using benchmark siRNA-delivering lipid MC3, commercial transfection reagent Lipo 2000, and symmetric four-tail lipids (2C12-D3 and 2O12-D3) as controls, we sequentially characterized the cellular uptake efficiency, *in vitro* gene silencing efficacy and *in vivo* biodistribution profiles of the resultant LNPs. Meanwhile, the SAR between lipid molecular structure and delivery performance was thoroughly analyzed.

#### 3.3.1 *In vitro* cellular uptake evaluation and SAR analysis

Flow cytometry was utilized to quantify the *in vitro* cellular uptake of all resultant LNPs. Cy5-siRNA was encapsulated in LNPs or Lipo 2000. This siRNA lacked sequence homology with any known mammalian genes and served as siNC in this study. HepG2 cells were incubated with 50 nM Cy5-siRNA@LNPs or Cy5-siRNA@Lipo 2000 for 4 h prior to flow cytometric detection of intracellular fluorescent signals. Overlaid flow cytometry histograms are presented in Fig. S3, and quantitative data of mean fluorescence intensity (MFI) for each group are summarized in Fig. 4A. Apart from O12-P6-H7 LNP, the remaining 33 newly developed LNPs achieved higher cellular uptake than MC3-LNP, among which 17 formulations exhibited statistically improved uptake efficiency.

**Fig. 4.**
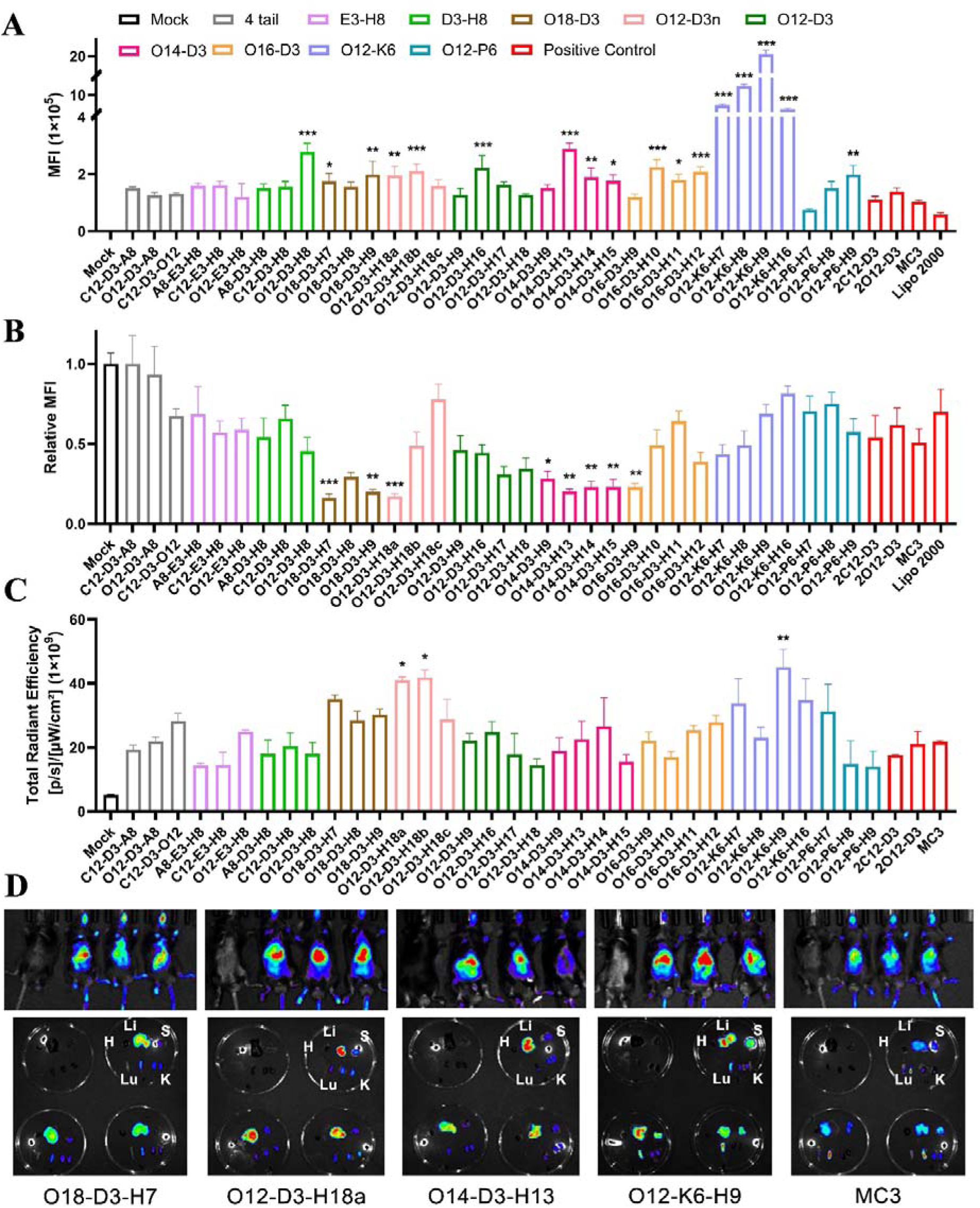
Evaluation of *in vitro* cellular uptake, gene silencing activity and *in vivo* biodistribution of dissymmetric hydrophobic-tailed ionizable lipid-based LNPs. (A) Mean fluorescence intensity (MFI) of HepG2 cells after incubation with 50 nM Cy5-siRNA@LNPs for 4 h, detected by flow cytometry. (B) Relative luciferase activity in HepG2-Luc cells after incubation with siLuc@LNPs for 48 h. (C) Average total radiant efficiency of C57BL/6 mice at 4 h after intravenous injection of Cy5-siRNA@LNPs *via* the tail vein at a siRNA dose of 1 mg/kg. (D) Representative *in vivo* fluorescence images (upper panel) and *ex vivo* organ images (lower panel) of mice treated with representative LNPs (O18-D3-H7, O12-D3-H18a, O14-D3-H13, O12-K6-H9) and the benchmark control MC3-LNP. In the *ex vivo* organ imaging, each culture dish displays the organs in the following order: heart, liver, spleen in the first row from left to right, and lung, kidney in the second row from left to right. Mock represents the untreated blank control group. Data are presented as mean ± SD from three independent experiments (n = 3). \*\*\**P* < 0.005, \*\**P* < 0.01, \**P* < 0.05 for test groups compared with the MC3-LNP group.

Subsequent SAR analysis revealed that the structure of hydrophobic tails strongly influenced cellular uptake. LNPs derived from dissymmetric-tailed lipids afforded comparable or superior internalization relative to symmetric controls 2C12-D3 and 2O12-D3, as exemplified by enhanced uptake of O12-D3-H8 relative to its symmetric counterpart 2O12-D3. In addition, three-tailed dissymmetric lipids generated LNPs with higher uptake than the four-tail series (4 tail). The O12-D3n series bearing unsaturated alkyl tails displayed robust overall uptake, with an average MFI 1.8-fold higher than MC3-LNP. By contrast, alkyl carbon chain length exerted marginal impacts on cellular uptake within the tested scope.

With regard to linker moiety, lipids functionalized with both amide and ester linkers conferred superior cellular uptake. Notably, O12-D3-H8 and O14-D3-H13 stood out with uptake levels 2.7-fold and 2.8-fold that of MC3-LNP, respectively. Consistently, D3-series lipids containing amide linkers generally outperformed E3-series lipids with solely ester linkers, such as the higher uptake of O12-D3-H8 over O12-E3-H8. Hydrophilic head groups also modulated uptake performance. O12-P6-H9 bearing piperazine-containing head groups and methylamino-based O12-K6 series LNPs both enabled prominent cellular uptake. Across all tested formulations, O12-K6 series LNPs achieved the highest internalization, and O12-K6-H9 exhibited uptake 19.5-fold higher than MC3-LNP and 35.2-fold higher than commercial Lipo 2000.

Collectively, LNPs constructed from these newly synthesized dissymmetric hydrophobic-tailed ionizable lipids exhibited overall improved cellular uptake compared with MC3 and symmetric lipid-based formulations, verifying the advantages of dissymmetric structural design to improve LNP cellular internalization. SAR results confirmed that three-tailed dissymmetric hydrophobic scaffolds, linkers bearing both amide and ester bonds, and polyamino hydrophilic head groups constitute critical structural motifs to maximize cellular uptake efficiency.

#### 3.3.2 *In vitro* transfection activity evaluation and SAR analysis

The HepG2-Luc cell line stably expressing firefly luciferase was used to evaluate the *in vitro* gene silencing efficacy of LNPs. Lower residual luciferase activity in the system indicated better siRNA-mediated target gene knockdown efficiency (Fig. 4B). Our results demonstrated that 19 of the novel dissymmetric hydrophobic-tailed lipid-based LNPs exhibited comparable or improved gene silencing effects relative to the control MC3-LNP, and 8 of these formulations showed statistically significant superiority.

Further SAR analysis revealed that the effect of hydrophobic tail structure on transfection activity followed a similar trend to that on cellular uptake. LNPs formulated from dissymmetric hydrophobic-tailed lipids outperformed those from symmetric lipid-based LNPs, and three-tailed dissymmetric lipids were superior to the four-tail series. However, the effect of hydrophobic tail unsaturation on transfection activity differed from that on cellular uptake. Specifically, transfection activity decreased with increasing hydrophobic tail unsaturation (O12-D3n series). O12-D3-H18a bearing a monounsaturated tail exhibited the most prominent transfection activity, with a gene silencing efficiency 1.7-fold that of MC3-LNP, while the efficiencies of diunsaturated and triunsaturated analogs decreased sequentially, indicating that moderate unsaturation is beneficial for transfection activity.

With regard to linker moiety, LNPs prepared from D3-series lipids containing both amide and ester bonds displayed higher silencing efficiency than those derived from E3-series lipids bearing only ester linker. The O18-D3 and O14-D3 series displayed excellent overall performance, and LNPs formulated from all seven lipids across the two series produced gene knockdown efficiencies exceeding 70%. LNPs formulated from O18-D3-H7 and O14-D3-H13 exhibited the optimal gene silencing activity, with knockdown efficiencies of 84% and 80%, respectively, which were 1.7-fold and 1.6-fold that of MC3-LNP. Hydrophilic head group architecture also had a significant effect on transfection activity, but its regulatory trend was not completely consistent with that on cellular uptake. Among the O12-K6 series with methylamino head groups and O12-P6 series bearing piperazine ring head groups, only LNPs formulated from O12-K6-H7 and O12-K6-H8 exhibited gene silencing efficiencies superior to or comparable to MC3-LNP.

Further analysis incorporating apparent p*K*_a_ data revealed that the 8 LNPs with significantly improved silencing efficiency relative to MC3-LNP had apparent p*K*_a_ values ranging from 5.7 to 6.2. Although this range was slightly lower than the commonly reported optimal pKa window (6.0-6.9) for ionizable lipid-based LNPs, it could still trigger the proton sponge effect in acidic endosomes and promote membrane fusion as well as efficient cytoplasmic release of siRNA. Notably, gene silencing activity was not completely proportional to cellular uptake efficiency. For example, the O12-K6-H9 LNP, which had the highest cellular uptake (19.5-fold that of MC3-LNP), showed a knockdown efficiency of 54%, comparable to MC3; whereas the O18-D3-H7 LNP with moderate cellular uptake (1.7-fold that of MC3-LNP) exhibited the highest knockdown efficiency (84%). This phenomenon fully demonstrated that gene silencing efficiency was jointly determined by cellular uptake efficiency and endosomal escape efficiency, and simply increasing uptake is not sufficient to ensure efficient cytoplasmic siRNA delivery.

#### 3.3.3 *In vivo* biodistribution evaluation and SAR analysis

To investigate the *in vivo* delivery potential of LNPs formulated from dissymmetric hydrophobic-tailed ionizable lipids, Cy5-siRNA@LNPs were intravenously administered to C57BL/6 mice *via* tail vein injection. *In vivo* fluorescence imaging was performed at 4 h post-injection (Fig. S4), and the average whole-body fluorescence intensity of each mouse group was quantified (Fig. 4C). Subsequently, mice were euthanized, and major organs (heart, liver, spleen, lung, kidney) were harvested for *ex vivo* imaging to characterize the tissue distribution profiles of LNPs (Fig.s S5 and Fig. 4D). The results revealed that 20 of the dissymmetric hydrophobic-tailed lipid-based LNPs exhibited comparable or higher fluorescence intensity relative to MC3-LNP, which implied these formulations possessed promising potential for *in vivo* siRNA delivery. Among them, three LNPs (O12-D3-H18a, O12-D3-H18b and O12-K6-H9) showed statistically significantly higher average whole-body fluorescence intensity than MC3-LNP, with 1.9-, 1.9- and 2.1-fold increases, respectively.

Regarding the relationship between lipid structure and *in vivo* fluorescence intensity, the SAR trends shared both similarities and differences with those from *in vitro* cellular uptake and transfection assays. For the similarities, LNPs derived from dissymmetric hydrophobic-tailed lipids outperformed symmetric counterparts, three-tailed dissymmetric lipids exhibited better performance than four-tail lipids, and D3-series lipids containing amide and ester bonds achieved superior *in vivo* efficacy compared with E3-series lipids bearing only ester linker. The primary differences lay in the effects of hydrophobic tail unsaturation and hydrophilic head groups. *In vitro* gene silencing activity declined with the increase in hydrophobic tail unsaturation. By contrast, the monounsaturated O12-D3-H18a and diunsaturated O12-D3-H18b exhibited comparable *in vivo* fluorescence intensity, both significantly higher than that of MC3-LNP, while the triunsaturated O12-D3-H18c showed inferior fluorescence intensity. Moreover, O12-K6-H9, a representative of the K6 series with weak *in vitro* transfection activity, yielded the highest *in vivo* fluorescence intensity, which was likely attributed to its excellent cellular uptake capability (Fig.s S4 and Fig. 4C).

*Ex vivo* organ imaging revealed that all LNPs displayed dominant hepatic fluorescence with limited accumulation in the spleen, lung and kidney, consistent with the well-documented clearance pathway of LNPs *via* the reticuloendothelial system [7, 8]. The organ distribution patterns were similar among all LNPs, and no obvious organ-specific targeting differences were observed. Collectively, this comprehensive SAR analysis established that the dissymmetric hydrophobic tail architecture was the key structural advantage, universally enhancing LNP performance relative to symmetric lipid scaffolds. Notably, D3-series lipids bearing both amide and ester bonds consistently outperformed single ester-based E3-series lipids across all cellular and *in vivo* evaluations, while moderate hydrophobic tail unsaturation balanced *in vitro* gene silencing efficacy and *in vivo* delivery activity (Fig.s S5 and Fig. 4D).

Based on the comprehensive multidimensional evaluation results of *in vitro* cellular uptake, gene silencing and *in vivo* biodistribution, we identified four lead ionizable lipids with distinct advantages for subsequent *in vivo* evaluation: O14-D3-H13 and O18-D3-H7 with the most prominent transfection activity, O12-D3-H18a with the most balanced *in vitro* and *in vivo* performance, and O12-K6-H9 with the best cellular uptake capacity and *in vivo* delivery efficacy. These lipids are abbreviated as O14, O18, H18a and K6 in the following experiments.

### 3.4 Preliminary *In Vitro* and *In Vivo* Safety Evaluation of Optimized LNPs

Safety is a critical prerequisite and major bottleneck for the clinical translation of nucleic acid drug delivery systems [42]. To evaluate the biocompatibility of four previously screened ionizable lipids (O14, O18, H18a, and K6), we systematically performed preliminary *in vitro* and *in vivo* safety assessments of the corresponding LNP formulations from multiple perspectives, including *in vitro* cytotoxicity, hemocompatibility, *in vivo* hematology and serum biochemistry in mice, and histopathological examination of major organs *via* hematoxylin-eosin (H&E) staining (Fig. 5 and Fig. S6). The objective was to verify the safety potential of candidate LNPs and lay an experimental foundation for subsequent pharmacodynamic studies and future clinical translation.

**Fig. 5.**
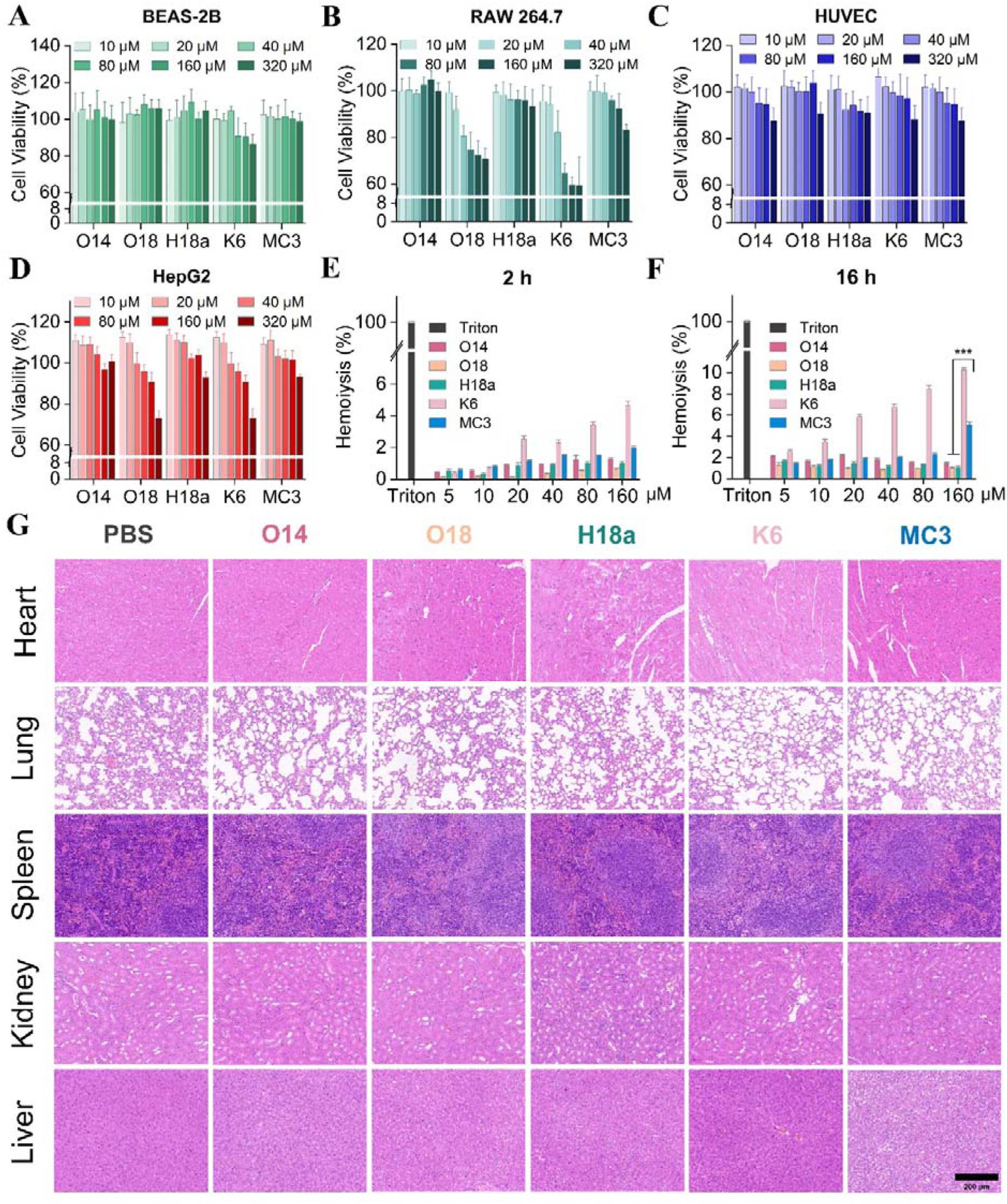
*In vitro* biocompatibility and *in vivo* systemic safety characterization of four lead ionizable lipid-based LNPs. (A-D) Cell viability of BEAS-2B, RAW 264.7, HUVEC and HepG2 cells after incubation with gradient concentrations (10, 20, 40, 80, 160, 320 μM) of O14-LNP, O18-LNP, H18a-LNP, K6-LNP and benchmark MC3-LNP for 24 h. (n = 3) (E, F) Quantitative hemolysis ratios of rabbit erythrocytes incubated with serially diluted LNPs after 2 h (E) and 16 h (F) incubation; Triton X-100 served as the positive control (n = 3). (G) Representative H&E stained histological micrographs of mouse heart, lung, spleen, kidney and liver harvested at day 7 post single intravenous injection of different LNPs (3 mg/kg); scale bar = 200 μm; (n = 5). Data are presented as mean ± SD, and \*\*\**P* < 0.005.

#### 3.4.1 *In vitro* cytotoxicity and hemocompatibility evaluation

The cytotoxicity of candidate LNPs at concentrations of 10, 20, 40, 80, 160, and 320 μM was evaluated using the CCK-8 assay in BEAS-2B, RAW 264.7, HUVEC, and HepG2 cells. LNPs formulated with the clinically benchmark ionizable lipid MC3 were used as the positive control (Fig. 5A-5D).

Different cell lines exhibited distinct sensitivities to LNP-induced cytotoxicity. In BEAS-2B cells, none of the four candidate LNPs showed significant cytotoxicity over the 10-320 μM concentration range, with cell viability consistently maintained above 80%. In RAW 264.7 macrophages, O14-LNP, H18a-LNP, and MC3-LNP retained cell viability above 80% at all tested concentrations. Specifically, O14-LNP and H18a-LNP maintained cell viability of 99% and 93%, respectively, at the maximum concentration of 320 μM, outperforming MC3-LNP (83%). In contrast, O18-LNP and K6-LNP exhibited significant cytotoxicity, with cell viability falling below 80% at 80 μM and declining further at higher concentrations. This observation is consistent with the biological characteristics of macrophages as professional phagocytes that are more prone to non-specific nanoparticle uptake. In HUVECs, all four candidate LNPs showed no significant cytotoxicity across the tested concentrations, with cell viability remaining above 80%. In HepG2 cells, O14-LNP, H18a-LNP, and MC3-LNP maintained cell viability above 80% at all tested concentrations, whereas O18-LNP and K6-LNP showed acceptable viability at lower concentrations but dropped below 80% at 320 μM, indicating potential hepatotoxicity. Collectively, these cytotoxicity results demonstrated that O14-LNP and H18a-LNP exhibited comparable or even better cytocompatibility than MC3-LNP across all four cell types, with a wider *in vitro* safety window.

Hemocompatibility is a core safety attribute for intravenously administered nanomedicines, as it directly determines their suitability for systemic *in vivo* administration. In this study, the hemolytic toxicity of candidate LNPs was evaluated using fresh rabbit erythrocyte suspensions (Fig. S6A and Fig. 5E, F). Following incubation with various concentrations of LNPs (5, 10, 20, 40, 80, and 160 μM) for 2 h and 16 h, hemolysis rates were calculated by quantifying hemoglobin release. PBS and 1% Triton X-100 served as negative and positive controls, respectively, with the clinically acceptable hemolysis threshold set at 5% [9, 42].

Macroscopic observation of post-centrifugation supernatants revealed that the Triton X-100 positive control group showed obvious red transparent hemolysis (Fig. S6A). K6-LNP exhibited visible erythrocyte rupture and hemolysis at multiple concentrations. MC3-LNP showed slight reddening of the supernatant at 160 μM after 16 h of incubation. In contrast, the supernatants of the O14-LNP, O18-LNP, and H18a-LNP groups remained colorless and clear, with no obvious signs of hemolysis. Quantitative analysis showed that after 2 h of incubation, the hemolysis rates of all candidate LNPs were below 5% at all tested concentrations (Fig. 5E). Among them, the hemolysis rates of O14-LNP, O18-LNP, H18a-LNP, and MC3-LNP were all below 2%, well below the clinical safety threshold. After 16 h of incubation, the hemolysis rates of O14-LNP, O18-LNP, and H18a-LNP remained below 2.5% at all concentrations (Fig. 5F). In contrast, K6-LNP exceeded the 5% threshold at a concentration as low as 20 μM. Notably, MC3-LNP showed a hemolysis rate of 5.1% at 160 μM, which was slightly above the clinical threshold and significantly higher than those of the O14-LNP, O18-LNP, and H18a-LNP groups (\*\*\**P* < 0.005). These results confirmed that O14-LNP, O18-LNP, and H18a-LNP possessed excellent hemocompatibility, outperforming the clinical benchmark MC3-LNP.

#### 3.4.2 *In Vivo* hematology, serum biochemistry, and histopathological evaluation

Thirty ICR mice were randomly divided into six groups and administered intravenously *via* the tail vein with PBS or 3 mg/kg of siNC@O14-LNP, siNC@O18-LNP, siNC@H18a-LNP, siNC@K6-LNP, or siNC@MC3-LNP. The mice were euthanized 7 days after administration. Blood samples were collected for routine hematology and serum biochemical analyses, and major organs (heart, liver, spleen, lung, and kidney) were harvested for H&E staining and histopathological examination (Fig. S6B, C and Fig. 5G).

Routine blood analysis showed that core hematological parameters, including RBC count, white blood cell (WBC) count, hemoglobin (HGB) level, platelet (PLT) count, neutrophil percentage (NEUT%), and lymphocyte percentage (LYMPH%), were within normal physiological ranges in all LNP-treated groups, with no statistically significant differences compared with the PBS group (Fig. S6B). Serum biochemical analysis revealed no abnormal elevations in liver function markers (alanine aminotransferase, ALT; aspartate aminotransferase, AST; albumin, ALB) or renal function markers (blood urea nitrogen, BUN; creatinine, CREA) in any LNP-treated group, with levels comparable to those in the PBS group (Fig. S6C).

Histopathological examination showed that all organs in the PBS group had normal structures with no pathological alterations (Fig. 5G). The tissue morphology of all organs in the O14-LNP, H18a-LNP, and MC3-LNP groups was highly consistent with that of the PBS group: myocardial fibers were neatly arranged with clear striations, alveolar walls showed no thickening or inflammatory infiltration, hepatocyte cords were regularly arranged with intact hepatic lobule structures, splenic red and white pulp were clearly demarcated, and glomeruli and renal tubules showed no degeneration or inflammation. The O18-LNP group showed essentially normal tissue structures with no obvious pathological damage. The K6-LNP group exhibited scattered mild inflammatory cell infiltration in some liver sections and occasional slight congestion in the renal interstitium, though the severity was relatively mild.

Collectively, the comprehensive *in vitro* and *in vivo* safety evaluation results demonstrated that O14-LNP and H18a-LNP exhibited favorable safety profiles in terms of *in vitro* cytotoxicity, hemocompatibility, *in vivo* hematological parameters, and histopathology, which were comparable or superior to those of the clinical benchmark MC3-LNP. Therefore, O14-LNP and H18a-LNP were selected for subsequent evaluations.

### 3.5 Mechanism on Intracellular Delivery of Candidate LNPs

Given the excellent *in vitro* and *in vivo* delivery performance and favorable biocompatibility of O14-LNP and H18a-LNP, we further investigated the molecular mechanisms underlying their high transfection efficiency, with a particular focus on endosomal escape capacity and cellular uptake pathways. This mechanistic dissection provides a theoretical foundation and scientific rationale for the subsequent therapeutic application of these candidate LNPs.

#### 3.5.1 Evaluation of Endosomal Escape Capacity

Efficient endosomal escape of siRNA into the cytoplasm is a critical step for achieving gene silencing [39, 43]. To evaluate the endosomal escape capacity of candidate LNPs (O14-LNP and H18a-LNP), confocal laser scanning microscopy (CLSM) was used to observe the intracellular distribution of Cy5-siRNA@LNPs in HepG2 cells, with Hoechst 33342 staining nuclei and LysoTracker Red labeling lysosomes. The Pearson’s correlation coefficient between Cy5-siRNA and LysoTracker Red fluorescence was quantified using ImageJ software as a measure of endosomal escape efficiency, with a lower coefficient indicating more effective escape [36].

Confocal fluorescence imaging results showed no obvious Cy5 fluorescence signal in the Mock group, while the Cy5 fluorescence intensity in O14-LNP and H18a-LNP groups was significantly higher than that in Lipo 2000 and MC3-LNP groups (Fig. 6A). Moreover, in Lipo 2000 and MC3-LNP groups, the green Cy5 fluorescence largely overlapped with the red lysosomal fluorescence, indicating that most siRNA was trapped in lysosomes. In contrast, Cy5 fluorescence in O14-LNP and H18a-LNP groups exhibited a diffuse cytoplasmic distribution, with significantly reduced colocalization with lysosomes. Quantitative analysis of intracellular MFI demonstrated that the MFI values of both O14-LNP and H18a-LNP groups were 1.7-fold those of the MC3-LNP group (Fig. 6B), confirming their excellent cellular uptake capacity, which was consistent with previous flow cytometry results (Fig. 4A and Fig. S3). Quantitative colocalization analysis revealed that the Pearson’s correlation coefficient between siRNA and lysosomes in the MC3-LNP group was 0.81, while those in O14-LNP and H18a-LNP groups decreased to 0.46 and 0.50, respectively, both significantly lower than that of MC3-LNP (Fig. 6C). These results indicated that O14-LNP and H18a-LNP could escape from acidic endosomes into the cytoplasm more efficiently, thereby effectively preventing siRNA degradation by lysosomal nucleases. This was one of the key molecular mechanisms underlying their superior transfection efficiency compared with MC3-LNP.

**Fig. 6.**
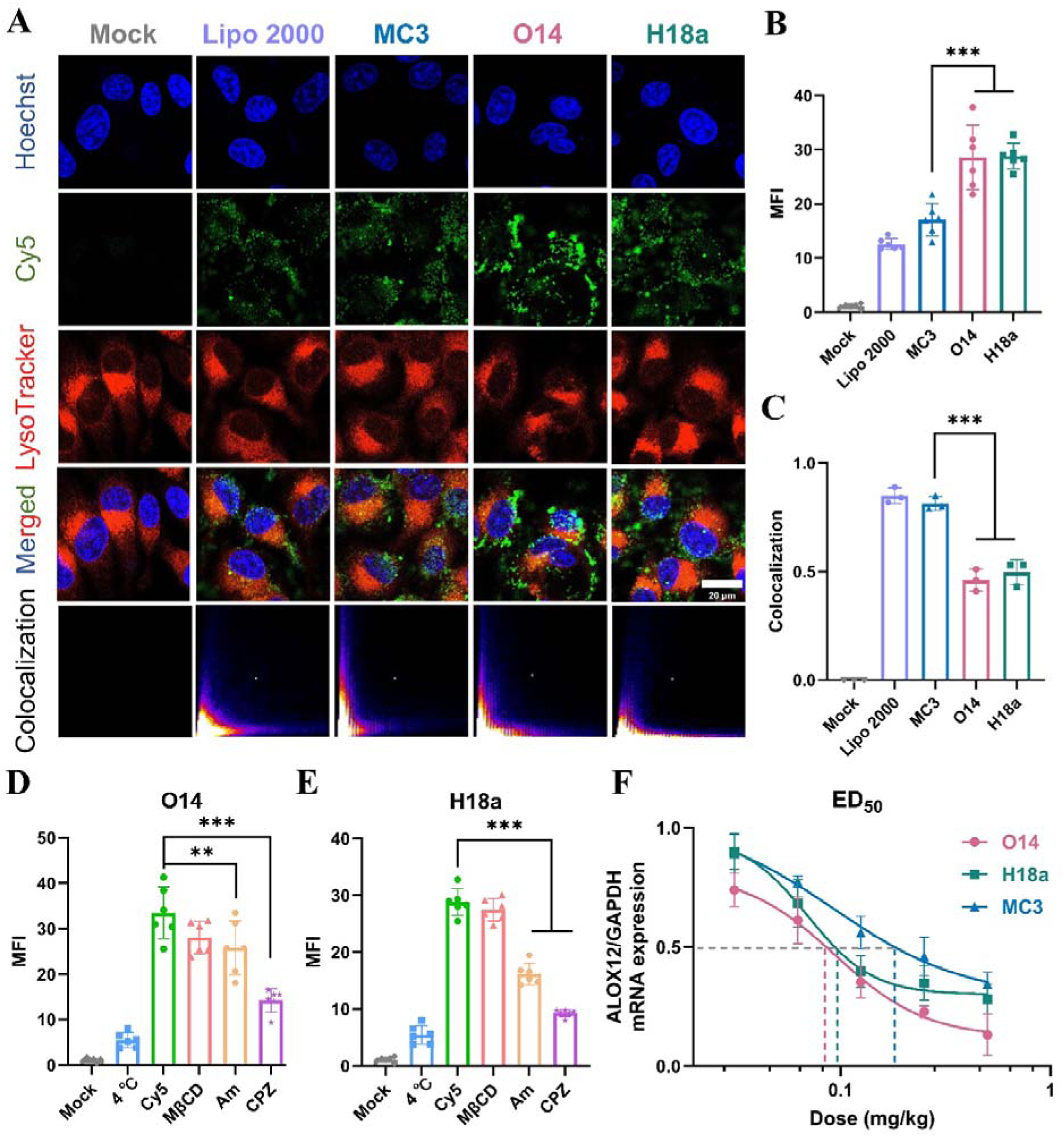
Endosomal escape, intracellular delivery and *in vivo* gene silencing performance of O14-LNP and H18a-LNP. (A) Representative confocal laser scanning microscopy (CLSM) images displaying the intracellular distribution of Cy5-siRNA (green) delivered by Mock, Lipo 2000, MC3-LNP, O14-LNP and H18a-LNP in HepG2 cells. Nuclei were stained with Hoechst 33342 (blue), and lysosomes were labeled with LysoTracker Red (red). Scale bar = 20 μm. (B) Quantitative analysis of intracellular Cy5 MFI for each formulation (n = 3). (C) Statistical quantification of Pearson’s colocalization coefficients between Cy5-siRNA and lysosomal signals (n = 3). (D, E) Quantitative MFI analysis of Cy5-siRNA internalized by O14-LNP (D) and H18a-LNP (E) under 4 °C cold treatment or pretreatment with methyl-β-cyclodextrin (MβCD), amiloride (Am), and chlorpromazine (CPZ) endocytosis inhibitors (n = 3). (F) Dose-response curves of ALOX12 mRNA knockdown mediated by MC3-LNP, O14-LNP and H18a-LNP in mice (n = 6). Data are presented as mean ± SD. \*\*\**P* < 0.005, \*\**P* < 0.01, for test groups compared with the MC3-LNP group or Cy5 group.

#### 3.5.2 Analysis of Cellular Internalization Pathways

In addition to efficient endosomal escape, differences in cellular internalization pathways also critically influence LNP transfection efficiency. To further elucidate the entry routes of the candidate LNPs, we first verified the energy dependence of their uptake *via* temperature-dependent experiments to exclude passive diffusion or non-specific adsorption. Subsequently, cells were pretreated with three specific endocytosis inhibitors: CPZ (clathrin-mediated endocytosis inhibitor), Am (macropinocytosis inhibitor), and MβCD (caveolae-mediated endocytosis inhibitor) [44]. Cellular uptake efficiency was quantitatively analyzed by CLSM to identify their predominant internalization pathways.

As shown in Fig.S6D, 6E and S7, the MFI values of both O14-LNP and H18a-LNP groups at 4°C decreased by more than 80% compared with the normal control group (Cy5 group), confirming that cellular internalization of both LNPs was an energy-dependent active process rather than passive diffusion. Pretreatment with CPZ reduced the MFI of O14-LNP and H18a-LNP by 57.4% and 67.7%, respectively, compared with their respective non-inhibited controls (\*\*\**P* < 0.005 for both). Am pretreatment decreased the MFI by 22.9% for O14-LNP (\*\**P* < 0.01) and 43.8% for H18a-LNP (\*\*\**P* < 0.005). In contrast, MβCD pretreatment had no significant effect on the uptake of either LNP (*P* > 0.05). These results demonstrated that O14-LNP and H18a-LNP were internalized into HepG2 cells primarily *via* clathrin-mediated endocytosis and macropinocytosis, while caveolae-mediated endocytosis was essentially not involved. Subtle differences in the dependence on specific endocytic pathways were observed between the two LNPs. H18a-LNP utilized both clathrin-mediated endocytosis and macropinocytosis, with clathrin-mediated endocytosis being the dominant route and macropinocytosis also contributing substantially. In contrast, O14-LNP relied predominantly on clathrin-mediated endocytosis as the core pathway, with macropinocytosis playing an auxiliary role. Notably, both pathways deliver nanoparticles to acidic endosomes, and the efficient endosomal escape described above allows effective siRNA release into the cytoplasm [45, 46]. The synergistic effect of these two processes collectively constituted the cellular basis for the superior transfection efficiency of O14-LNP and H18a-LNP.

### 3.6 Quantitative Evaluation of *In Vivo* Gene Silencing Activity of Candidate LNPs

Having elucidated the intracellular delivery mechanism underlying the high transfection efficiency of O14-LNP and H18a-LNP, we further carried out *in vivo* evaluation of transfection and gene silencing efficacy for ferroptosis-mediated T2D. As mentioned above, ALOX12 acts as a key functional gene regulating cellular ferroptosis and lipid peroxidation. Its aberrant overexpression exacerbates hyperglycemia-induced metabolic disorders as well as oxidative damage in pancreatic islets and liver, serving as a critical pathogenic target driving the progression of T2D [29, 30]. Based on this, we established an *in vivo* gene silencing model targeting ALOX12 and quantitatively evaluated the transfection performance of the two candidate LNPs *via* multi-dose gradient experiments, with MC3-LNP serving as the benchmark to precisely quantify their pharmacodynamic advantages.

Dose-response curve analysis showed that all three LNPs significantly downregulated ALOX12 mRNA expression in a dose-dependent manner (Fig. 6F). At the low dose of 0.06 mg/kg, MC3-LNP achieved only 23% ALOX12 knockdown, while O14-LNP and H18a-LNP already achieved 39% and 32% knockdown, respectively. At a dose of 0.125 mg/kg, the knockdown efficiency of MC3-LNP was 44%, whereas those of O14-LNP and H18a-LNP increased to 65% and 60%, respectively. At the highest dose of 0.5 mg/kg, both O14-LNP and H18a-LNP achieved knockdown efficiencies exceeding 75%, significantly higher than the 66% achieved by MC3-LNP. The half-maximal effective dose (ED_50_) was calculated *via* nonlinear regression fitting of the dose-response curves. The ED_50_ value of MC3-LNP was approximately 0.17 mg/kg, while those of O14-LNP and H18a-LNP were 0.08 mg/kg and 0.10 mg/kg, representing 53% and 41% reductions compared with MC3-LNP, respectively. These results confirmed that O14-LNP and H18a-LNP possessed superior *in vivo* gene silencing activity, enabling potent target gene knockdown at significantly lower doses. This advantage laid a solid pharmacodynamic foundation for their subsequent application in the treatment of ferroptosis-related diseases.

### 3.7 Evaluation of the *In Vitro* and *In Vivo* Therapeutic Effects of Candidate LNPs in Delivering Ferroptosis-related siRNA for Treating Type 2 Diabetes

Ferroptosis has been established as one of the core pathological mechanisms mediating pancreatic β-cell dysfunction, hepatic lipid metabolic disturbance and insulin resistance in T2D [25–27]. Targeted modulation of key ferroptosis genes *via* siRNA holds great promise as a novel therapeutic strategy for T2D intervention. Previous experiments confirmed that O14-LNP and H18a-LNP exhibited excellent cellular uptake efficiency and endosomal escape capacity *in vitro*, with transfection activity significantly superior to that of the classic benchmark carrier MC3-LNP. *In vivo*, both LNPs achieved robust silencing of the ferroptosis regulatory gene ALOX12, with markedly reduced ED_50_ values relative to MC3-LNP, demonstrating greatly enhanced *in vivo* nucleic acid delivery efficacy and therapeutic potential. However, gene silencing performance at the molecular level only reflects the nucleic acid delivery capability of the carriers. Whether they can effectively mitigate oxidative stress damage, ameliorate glucose and lipid metabolic disorders, and alleviate pathological damage to target organs during T2D progression by targeting the ferroptosis pathway still requires systematic therapeutic validation at both cellular and whole-animal levels. Accordingly, we systematically evaluated the therapeutic efficacy of candidate LNPs delivering ferroptosis-targeted siRNAs in a high glucose-induced ferroptosis model of RAW 264.7 macrophages, a STZ-induced T2D mouse model, and a db/db spontaneous diabetic mouse model, to comprehensively elucidate their application value and translational potential as a nucleic acid delivery platform for ferroptosis-based therapy.

#### 3.7.1 Evaluation of the Activity of LNPs in Inhibiting LPO in the Ferroptosis Cell Model

To preliminarily validate the therapeutic potential of candidate LNPs in ferroptosis-related diseases, we evaluated at the cellular level the inhibitory effect of siALOX12 (siALOX) delivered by these LNPs on LPO, a key hallmark of ferroptosis [47]. A ferroptosis cell model was established by stimulating RAW 264.7 macrophages with 30 mM glucose, and intracellular LPO levels were detected using C11-BODIPY (581/591), a specific fluorescent probe for LPO. Confocal imaging combined with quantitative analysis was employed to compare the intervention efficacy of different LNPs loaded with siALOX.

As shown in Fig. S8, the fluorescence intensity of oxidized C11-BODIPY in the high glucose stimulation group was significantly elevated compared with the PBS group (^###^*P* < 0.005), indicating that high glucose successfully induced cellular lipid peroxidation damage. After treatment with siALOX-loaded LNPs, intracellular LPO levels in all three groups were markedly reduced relative to the high glucose model group (\*\*\**P* < 0.005), confirming that all three carriers effectively suppressed high glucose-induced ferroptosis by delivering siALOX. Intergroup comparison revealed that the LPO level in the siALOX@H18a-LNP group was comparable to that in the siALOX@MC3-LNP positive control group, whereas the LPO level in the siALOX@O14-LNP group was slightly higher than that in the siALOX@MC3-LNP group (\**P* < 0.05), suggesting its moderately weaker ferroptosis-inhibitory activity than that of the classical control carrier.

From a mechanistic perspective, LPO accumulation is not only a core biological feature of ferroptosis but also a critical pathological basis for islet injury and hepatic metabolic dysfunction under diabetic conditions [48–50]. ALOX12, a positive regulator of ferroptosis, directly promotes LPO generation by catalyzing the oxidation of polyunsaturated fatty acids in cell membranes. Its aberrant upregulation under high glucose conditions serves as a key driver of cellular ferroptosis and metabolic damage. Therefore, silencing this gene can block the initiation and progression of ferroptosis at the source [29, 30]. In this study, all three LNPs effectively reversed high glucose-induced oxidative damage by delivering siALOX, functionally validating the regulatory role of the ALOX12 target in macrophage ferroptosis. Notably, H18a-LNP exhibited *in vitro* activity comparable to that of MC3-LNP. Collectively, these cellular-level findings provide a solid cytological basis for subsequent *in vivo* evaluations of the therapeutic efficacy of these LNPs in diabetic models.

#### 3.7.2 Evaluation of the Therapeutic Efficacy in the STZ-induced Type 2 Diabetic Mouse Model

To verify the therapeutic potential of candidate LNPs delivering ferroptosis-related siRNA (siALOX) in T2D, we established a T2D mouse model *via* a 50-day high-fat diet followed by a single intraperitoneal injection of STZ (100 mg/kg) [51]. The diabetic mice were then treated with siALOX@O14-LNP, siALOX@H18a-LNP or siALOX@MC3-LNP *via* weekly tail vein injection at 0.25 mg/kg for 4 consecutive weeks, with the STZ model group and normal C57BL/6 group serving as controls (Fig. 7A). The therapeutic effects were comprehensively evaluated from multiple dimensions, including the regulation of glucose and lipid metabolism, restoration of islet function, amelioration of hepatic pathology, and modulation of the ferroptosis pathway.

**Fig. 7.**
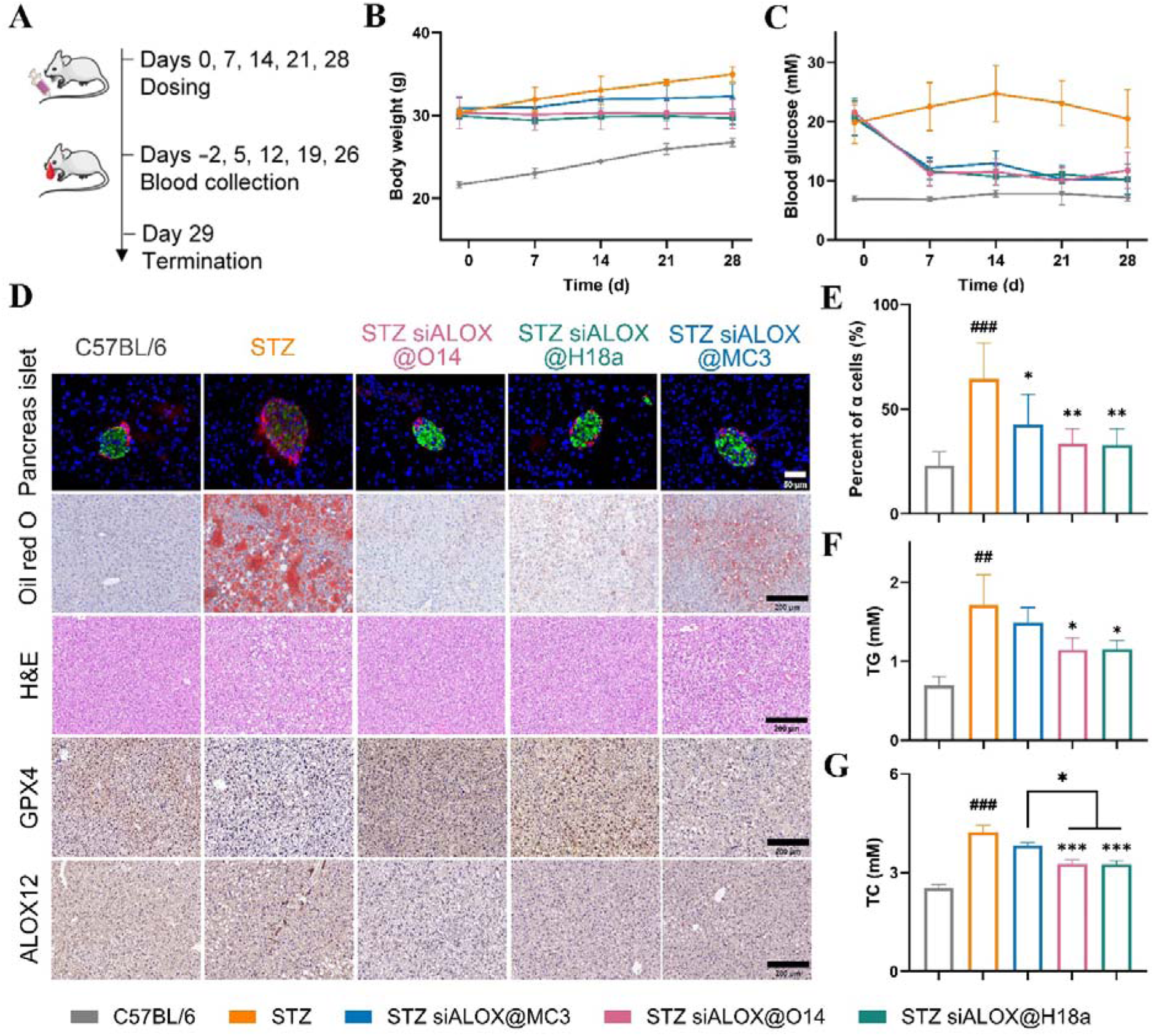
Therapeutic efficacy of siALOX-loaded LNPs in STZ-induced type 2 diabetic mice. (A) Schematic diagram of the experimental protocol. (B) Dynamic changes in body weight of mice in each group during the 28-day treatment period. (C) Time-course changes in fasting blood glucose levels across all groups. (D) Representative images of pancreatic islet immunofluorescence staining (green: insulin; red: glucagon; scale bar = 20 μm), hepatic Oil Red O staining (scale bar = 200 μm), hepatic H&E staining (scale bar = 200 μm), and immunohistochemical staining of GPX4 and ALOX12 in liver tissues (scale bar = 200 μm). (E) Quantitative analysis of α-cell to β-cell ratio in pancreatic islets. (F) Serum triglyceride (TG) levels in each group after 4 weeks of treatment. (G) Serum total cholesterol (TC) levels in each group after 4 weeks of treatment. Data are presented as the mean ± SD from five independent experiments (n = 5), and ^###^*P* < 0.005, ^##^*P* < 0.01, STZ group compared to the C56BL/6 group; \*\*\**P* < 0.005, \*\**P* < 0.01, \**P* < 0.05, test groups compared to the STZ group.

Body weight and blood glucose monitoring showed that mice in the STZ-induced diabetic group had significantly higher body weight than the normal controls, exhibiting a classic T2D phenotype of obesity combined with hyperglycemia (Fig. 7B, C). After intervention with siALOX@LNPs, the body weight growth rate of each treatment group was slower than that of the model group. On day 28, the body weights of the siALOX@O14-LNP and siALOX@H18a-LNP groups were 30.18 ± 1.75 g and 29.70 ± 0.79 g, respectively, both lower than that of the siALOX@MC3-LNP group (32.30 ± 1.50 g). In terms of blood glucose, the fasting blood glucose in the STZ model group was markedly elevated and persistently maintained above 20 mM. In contrast, the fasting blood glucose in all three siALOX@LNPs groups significantly decreased to 11-12 mM on day 7 after the first injection and remained stable throughout the experiment, with comparable glucose-lowering efficacy among the three groups. These results indicated that the two candidate LNPs delivering siALOX effectively suppressed excessive weight gain and reversed hyperglycemia in STZ-induced diabetic mice, with overall performance comparable to or even better than that of MC3-LNP.

Islet dysfunction is a core pathological feature of T2D, mainly characterized by excessive glucagon secretion and an imbalanced α-cell to β-cell (α/β cell) ratio [52]. Immunofluorescence staining of pancreatic tissues showed markedly upregulated glucagon expression in the STZ model group, which was reduced to varying degrees after siALOX@LNPs treatment (Fig. 7D). Quantitative analysis of islet cell proportions further showed that the α/β cell ratio in normal islets was 22.9%, whereas it significantly increased to 64.5% in the STZ model group (^###^*P* < 0.005), consistent with the pathological features of abnormal α-cell proliferation and glucagon overactivity in diabetes (Fig. 7E). After treatment, the α/β cell ratio declined to 33.5% and 32.9% in the siALOX@O14-LNP and siALOX@H18a-LNP groups, respectively (\*\**P* < 0.01 for both), and to 42.6% in the siALOX@MC3-LNP group (\**P* < 0.05). These findings suggested that O14-LNP and H18a-LNP were more effective at reversing islet cell imbalance and restoring islet structure and function.

The liver acts as the central organ for glucose and lipid metabolism and a key target for ferroptosis regulation, and is also the primary site of LNP accumulation *in vivo*. Oil Red O staining showed massive lipid droplet accumulation and severe ectopic lipid deposition in the liver parenchyma of STZ model mice, consistent with hepatic steatosis associated with T2D (Fig. 7D). After siALOX@LNPs treatment, both the number and area of hepatic lipid droplets were markedly reduced, with the siALOX@O14-LNP and siALOX@H18a-LNP groups exhibiting no evident large-area lipid droplet aggregation, outperforming the siALOX@MC3-LNP group. H&E staining further confirmed that the STZ model and siALOX@MC3-LNP groups displayed vacuolar changes due to lipid accumulation, whereas the livers of the siALOX@O14-LNP and siALOX@H18a-LNP groups showed normal tissue architecture without obvious pathological alterations (Fig. 7D). The marked amelioration of hepatic pathology confirmed that candidate LNPs exerted metabolic regulation and organ protection *via* liver-targeted delivery.

At the molecular level, gene and protein detection in liver tissues further verified the targeted modulation of the ferroptosis pathway by the formulations. Immunohistochemistry and qPCR results revealed that the expression of GPX4, a key ferroptosis defense protein [35], was downregulated in the livers of STZ model mice, indicating hyperactivation of the hepatic ferroptosis pathway in diabetes (Fig. 7D). After siALOX@LNPs intervention, both ALOX12 mRNA and protein levels were significantly reduced, and GPX4 protein expression was markedly restored in all treatment groups (Fig. 7D, S9A). These findings demonstrated that candidate LNPs inhibited hepatic ferroptosis by efficiently silencing the hepatic target gene ALOX12 and upregulating the ferroptosis defense molecule GPX4, which represented the core mechanism underlying their effects on glucose and lipid metabolism regulation and organ protection.

In addition, serum lipid profiling showed that triglyceride (TG) and total cholesterol (TC) levels were significantly elevated in the STZ model group, reflecting diabetes-associated dyslipidemia (Fig. 7F, G). After treatment with candidate LNPs, both TG and TC levels decreased remarkably, with superior improvements to those observed in the MC3-LNP group, demonstrating the corrective effect of the formulations on systemic lipid metabolism disorders. There were no significant intergroup differences in hepatic and renal function markers, including AST, BUN, CREA and ALB, nor were there significant differences in any routine blood parameters (Fig. S9B-G). ALT levels were comparable between the STZ model and normal control groups, yet were significantly lowered in the siALOX@MC3-LNP and siALOX@H18a-LNP groups compared with the STZ group. Combined with the absence of overt pathological abnormalities in H&E-stained major organs (heart, spleen, lung, kidney), these results indicated that severe organic impairment of liver and kidney function had not developed at this disease stage. Notably, repeated administration of LNP formulations did not induce additional hepatic or renal injury or histopathological lesions even under diabetic pathological conditions, further verifying the favorable *in vivo* biocompatibility of the formulations (Fig. S9H).

Collectively, in the STZ-induced T2D mouse model, O14-LNP and H18a-LNP delivering siALOX modulated the ferroptosis pathway by efficiently silencing hepatic ALOX12, and exerted therapeutic benefits at multiple levels including glucose and lipid metabolism regulation, islet structural restoration, hepatic lipid deposition improvement, and molecular pathway modulation. Their overall therapeutic efficacy was superior to that of the classic benchmark carrier MC3-LNP, providing reliable *in vivo* evidence for ferroptosis-targeted nucleic acid therapy against T2D.

#### 3.7.3 Evaluation of the Therapeutic Efficacy and Other Targets Expansion Verification in the db/db Spontaneous Type 2 Diabetic Mouse Model

To further evaluate the therapeutic potential of candidate LNPs in a model with a genetic background more analogous to that of human T2D, and to explore the feasibility of targeting additional key regulators of ferroptosis, we employed leptin receptor-deficient db/db spontaneous diabetic mice in this study [53]. H18a-LNP was used as the delivery vector to encapsulate siRNAs targeting ACSL4 and Keap1, two critical mediators of the ferroptosis pathway [31–34]. Separate low-dose (0.25 mg/kg) and high-dose (0.5 mg/kg) groups were established for siACSL4@H18a-LNP and siKeap1@H18a-LNP, with db/db PBS mice and wild-type m/m PBS mice set as controls. All formulations were administered *via* tail vein injection once weekly for 4 consecutive weeks, and therapeutic outcomes were comprehensively assessed in terms of systemic glucose and lipid homeostasis, target gene silencing efficiency, and histopathological improvement.

Monitoring of body weight and fasting blood glucose revealed that db/db model mice exhibited significantly higher body weight than m/m normal controls, consistent with the classic obese-hyperglycemic phenotype driven by hyperphagia and metabolic dysregulation resulting from leptin receptor deficiency (Fig. 8A, B). Following intervention with H18a-LNP-encapsulated targeted siRNAs, the rate of body weight gain was reduced in all treatment groups compared with the db/db model group, and the effect was more pronounced in the high-dose groups, exhibiting a moderate dose-dependent trend. For glycemic control, fasting blood glucose in the db/db model group remained persistently elevated above 25 mM. In contrast, fasting blood glucose in all treatment groups declined notably by day 7 after the first administration and was maintained at lower levels throughout the treatment period. The glucose-lowering effect was more robust in the 0.5 mg/kg high-dose groups. By the end of the intervention, blood glucose in both the siACSL4 and siKeap1 high-dose groups stably decreased to approximately 14 mM, demonstrating a clear dose-dependent pattern. These results indicated that targeted silencing of either ACSL4 or Keap1 effectively ameliorated obesity and hyperglycemia in db/db mice, and H18a-LNP exerted reliable regulatory effects on glucose and lipid metabolism *in vivo* when delivering siRNAs against distinct ferroptosis targets.

**Fig. 8.**
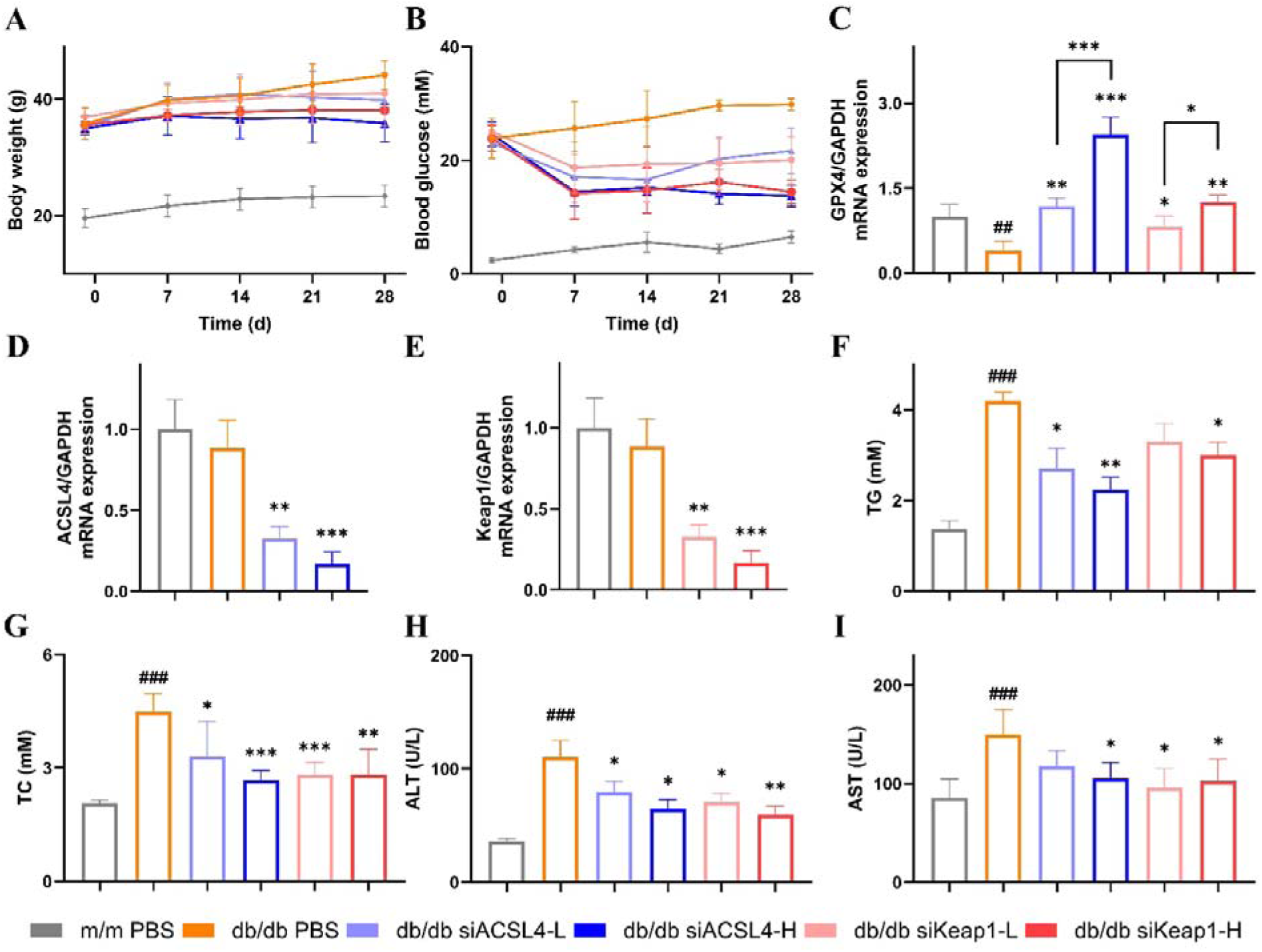
Therapeutic efficacy of siACSL4 and siKeap1 delivered by H18a-LNP in db/db spontaneous type 2 diabetic mice. (A, B) Time-course changes in body weight (A) and fasting blood glucose (B) across groups during the 28-day treatment. (C-E) Relative mRNA expression of GPX4 (C), ACSL4 (D), and Keap1 (E) in liver tissues determined by qPCR. (F-I) Serum biochemical parameters: triglyceride (TG, F), total cholesterol (TC, G), alanine aminotransferase (ALT, H), and aspartate aminotransferase (AST, I). Data are presented as the mean ± SD from five independent experiments (n = 5), and ^###^*P* < 0.005, ^##^*P* < 0.01, db/db group compared to the m/m group; \*\*\**P* < 0.005, \*\**P* < 0.01, \**P* < 0.05, test groups compared to the db/db group.

qPCR analysis of liver tissues further confirmed target gene silencing and pathway modulation at the molecular level. Relative to m/m normal controls, db/db model mice exhibited significantly lower hepatic GPX4 mRNA expression, which indicated hyperactivation of the ferroptosis pathway under spontaneous diabetic conditions (Fig. 8C). After treatment with H18a-LNP-delivered siRNAs, the mRNA levels of target genes (ACSL4 or Keap1) were dose-dependently reduced in all treatment groups, along with a marked restoration of GPX4 mRNA expression (Fig. 8C, D, E). These findings verified that H18a-LNP efficiently delivered siRNAs to hepatic target cells, effectively silenced ACSL4 or Keap1, and restored the expression of the ferroptosis defense molecule GPX4, thereby suppressing excessive ferroptosis pathway activation at the transcriptional level. Moreover, these results demonstrated the versatile delivery capacity of H18a-LNP as a carrier for siRNAs with distinct sequences.

Serum biochemical analyses showed that db/db model mice had significantly elevated levels of TG, TC, ALT and AST compared with m/m controls (Fig. 8F-I). Consistent with this, H&E staining of liver sections revealed extensive vacuolar alterations resulting from lipid accumulation, reflecting severe lipid dysregulation and hepatic functional impairment in the diabetic model (Fig. S10). Following siACSL4 or siKeap1 treatment, serum lipid profiles and liver enzyme levels were dose-dependently improved, and both the number and area of hepatic lipid vacuoles were markedly reduced, indicating substantial alleviation of hepatic pathological damage. In addition, BUN levels were significantly lower in the db/db model group than in wild-type m/m controls, and showed partial recovery after treatment (Fig. S10). No significant intergroup differences were observed in CREA levels or any routine blood parameters, and no overt pathological abnormalities were detected by H&E staining in major organs including the heart, spleen, lung and kidney (Fig. S10). These results demonstrated that ferroptosis-targeted intervention effectively mitigated diabetes-associated hepatic lipid deposition and functional injury, and the formulations did not cause additional tissue damage within the effective dose range, supporting their favorable *in vivo* biocompatibility.

In summary, in the db/db model of spontaneous T2D, H18a-LNP-mediated delivery of siACSL4 or siKeap1 effectively silenced hepatic target genes, upregulated GPX4 expression, and consequently improved hyperglycemia, corrected lipid dysregulation, and attenuated hepatic steatosis in a dose-dependent manner. These findings corroborated previous results from the STZ-induced model, further strengthening the experimental basis for ferroptosis intervention as a novel therapeutic strategy for T2D. Moreover, this study validated the broad applicability of H18a-LNP for delivering siRNAs targeting different ferroptosis molecules, providing critical preclinical support for future multi-target combination therapy and further expansion of this delivery platform.

## 4. Conclusion

In this work, we developed a library of 34 novel ionizable lipids featuring dissymmetric hydrophobic tails, a design inspired by the dissymmetric structural properties of natural cell membrane phospholipids that facilitate membrane fusion and non-lamellar phase transition. Systematic physicochemical characterization and multidimensional biological evaluations established clear SAR: three-tailed dissymmetric hydrophobic scaffolds, linkers bearing both amide and ester bonds, and moderate hydrophobic tail unsaturation conferred optimal delivery performance. Two lead formulations, O14-LNP and H18a-LNP, exhibited significantly enhanced cellular uptake, more efficient endosomal escape, and superior *in vivo* gene silencing activity compared with the clinical benchmark MC3-LNP, along with favorable biocompatibility. Mechanistic studies confirmed that both LNPs are internalized primarily *via* clathrin-mediated endocytosis and macropinocytosis. Furthermore, we validated the therapeutic potential of the lead LNP platform in two T2D mouse models, demonstrating that delivery of siRNAs targeting distinct ferroptosis regulators effectively improved glucose and lipid homeostasis, restored pancreatic islet structure, and mitigated hepatic steatosis. These findings not only provide a rational design paradigm for next-generation ionizable lipids, but also offer a robust and versatile delivery platform for ferroptosis-targeted nucleic acid therapeutics.

## CRediT authorship contribution statement

**Han Zhang:** Writing-original draft, Conceptualization, Software, Investigation, Formal analysis, Visualization, Methodology, Data curation, Project administration; **Yuanyuan Liu:** Conceptualization, Methodology, Software, Data curation, Investigation, Formal analysis, Writing-original draft, Visualization; **Fengyang He:** Methodology, Software, Data curation, Investigation, Formal analysis; **Genping Xue:** Investigation, Data curation, Formal analysis; **Yi Kang:** Investigation, Formal analysis, Data curation; **Zepu Zhang:** Validation, Investigation; **Jixin Ma:** Validation; **Junhai Xiao:** Conceptualization, Methodology, Supervision, Writing-review & editing; **Qingbin Meng:** Writing-review & editing, Conceptualization, Methodology, Supervision, Project administration, Resources.

## Data Availability Statement

Data will be made available on reasonable request.

## References

[1] K. A. Whitehead, R. Langer, D. G. Anderson, “Knocking Down Barriers: Advances in siRNA Delivery,” Nat. Rev. Drug Discov., 8 (2009), pp. 129–138, 10.1038/nrd2742.

[2] S. M. Elbashir, J. Harborth, W. Lendeckel, A. Yalcin, K. Weber, T. Tuschl, “Duplexes of 21-Nucleotide RNAs Mediate RNA Interference in Cultured Mammalian Cells,” Nature, 411 (2001), pp. 494–498, 10.1038/35078107.

[3] M. Friedrich, A. Aigner, “Therapeutic siRNA: State-of-the-Art and Future Perspectives,” BioDrugs, 36 (2022), pp. 549–571, 10.1007/s40259-022-00549-3.

[4] B. Hu, L. Zhong, Y. Weng, L. Peng, Y. Huang, Y. Zhao, X. J. Liang, “Therapeutic siRNA: State of the Art,” Signal Transduct. Target. Ther., 5 (2020), pp. 101–126, 10.1038/s41392-020-0207-x.

[5] R. Kanasty, J. R. Dorkin, A. Vegas, D. Anderson, “Delivery Materials for siRNA Therapeutics,” Nat. Mater., 12 (2013), pp. 967–977, 10.1038/nmat3765.

[6] X. Hou, T. Zaks, R. Langer, Y. Dong, “Lipid Nanoparticles for mRNA Delivery,” Nat. Rev. Mater., 6 (2021), pp. 1078–1094, 10.1038/s41578-021-00358-0.

[7] J. A. Kulkarni, P. R. Cullis, R. van der Meel, “Lipid Nanoparticles Enabling Gene Therapies: From Concepts to Clinical Utility,” Nucleic Acid Ther., 28 (2018), pp. 146–157, 10.1089/nat.2018.0721.

[8] J. A. Kulkarni, D. Witzigmann, S. Chen, P. R. Cullis, R. van der Meel, “Lipid Nanoparticle Technology for Clinical Translation of siRNA Therapeutics,” Acc. Chem. Res., 52 (2019), pp. 2435–2444, 10.1021/acs.accounts.9b00368.

[9] C. H. Albertsen, J. A. Kulkarni, D. Witzigmann, M. Lind, K. Petersson, J. B. Simonsen, “The Role of Lipid Components in Lipid Nanoparticles for Vaccines and Gene Therapy,” Adv. Drug Deliv. Rev., 188 (2022), Article 114416, 10.1016/j.addr.2022.114416.

[10] X. Han, H. Zhang, K. Butowska, K. L. Swingle, M. G. Alameh, D. Weissman, M. J. Mitchell, “An Ionizable Lipid Toolbox for RNA Delivery,” Nat. Commun., 12 (2021), Article 7233, 10.1038/s41467-021-27493-0.

[11] S. C. Semple, A. Akinc, J. Chen, A. P. Sandhu, B. L. Mui, C. K. Cho, D. W. Y. Sah, D. Stebbing, E. J. Crosley, E. Yaworski, I. M. Hafez, J. R. Dorkin, J. Qin, K. Lam, K. G. Rajeev, K. F. Wong, L. B. Jeffs, L. Nechev, M. L. Eisenhardt, M. Jayaraman, M. Kazem, M. A. Maier, M. Srinivasulu, M. J. Weinstein, Q. Chen, R. Alvarez, S. A. Barros, S. De, S. K. Klimuk, T. Borland, V. Kosovrasti, W. L. Cantley, Y. K. Tam, M. Manoharan, M. A. Ciufolini, M. A. Tracy, A. de Fougerolles, I. MacLachlan, P. R. Cullis, T. D. Madden, M. J. Hope, “Rational Design of Cationic Lipids for siRNA Delivery,” Nat. Biotechnol., 28 (2010), pp. 172–176, 10.1038/nbt.1602.

[12] A. Akinc, M. A. Maier, M. Manoharan, K. Fitzgerald, M. Jayaraman, S. Barros, S. Ansell, X. Du, M. J. Hope, T. D. Madden, B. L. Mui, S. C. Semple, Y. K. Tam, M. Ciufolini, D. Witzigmann, J. A. Kulkarni, R. van der Meel, P. R. Cullis, “The Onpattro Story and the Clinical Translation of Nanomedicines Containing Nucleic-Acid-Based Drugs,” Nat. Nanotechnol., 14 (2019), pp. 1084–1087, 10.1038/s41565-019-0591-y.

[13] D. Adams, A. G. Duarte, W. D. O’Riordan, C. C. Yang, M. Ueda, A. V. Kristen, I. Tournev, H. H. Schmidt, T. Coelho, J. L. Berk, K. P. Lin, G. Vita, S. Attarian, V. P. Bordeneuve, M. M. Mezei, J. M. Campistol, J. Buades, T. H. Brannagan, B. J. Kim, J. Oh, Y. Parman, Y. Sekijima, P. N. Hawkins, S. D. Solomon, M. Polydefkis, P. J. Dyck, P. J. Gandhi, S. Goyal, J. Chen, A. L. Strahs, S. V. Nochur, M. T. Sweetser, P. P. Garg, A. K. Vaishnaw, J. A. Gollob, O. B. Suhr, “Patisiran, an RNAi Therapeutic, for Hereditary Transthyretin Amyloidosis,” N. Engl. J. Med., 379 (2018), pp. 11–21, 10.1056/NEJMoa1716153.

[14] M. Jayaraman, S. M. Ansell, B. L. Mui, Y. K. Tam, J. Chen, X. Du, D. Butler, L. Eltepu, S. Matsuda, J. K. Narayanannair, K. G. Rajeev, I. M. Hafez, A. Akinc, M. A. Maier, M. A. Tracy, P. R. Cullis, T. D. Madden, M. Manoharan, M. J. Hope, “Maximizing the Potency of siRNA Lipid Nanoparticles for Hepatic Gene Silencing In Vivo,” Angew. Chem. Int. Ed., 51 (2012), pp. 8529–8533, 10.1002/anie.201203263.

[15] Y. Eygeris, M. Gupta, J. Kim, G. Sahay, “Chemistry of Lipid Nanoparticles for RNA Delivery,” Acc. Chem. Res., 55 (2022), pp. 2–12, 10.1021/acs.accounts.1c00544.

[16] Y. Dong, D. J. Siegwart, D. G. Anderson, “Strategies, Design, and Chemistry in siRNA Delivery Systems,” Adv. Drug Deliv. Rev., 144 (2019), pp. 133–147, 10.1016/j.addr.2019.05.004.

[17] P. R. Cullis, B. de Kruijff, “Lipid Polymorphism and the Functional Roles of Lipids in Biological Membranes,” Biochim. Biophys. Acta, 559 (1979), pp. 399–420, 10.1016/0304-4157(79)90012-1.

[18] I. M. Hafez, N. Maurer, P. R. Cullis, “On the Mechanism Whereby Cationic Lipids Promote Intracellular Delivery of Polynucleic Acids,” Gene Ther., 8 (2001), pp. 1188–1196, 10.1038/sj.gt.3301506.

[19] D. Zhang, E. N. Atochina-Vasserman, J. Lu, D. S. Maurya, Q. Xiao, M. Liu, J. Adamson, N. Ona, E. K. Reagan, H. Ni, D. Weissman, V. Percec, “The Unexpected Importance of the Primary Structure of the Hydrophobic Part of One-Component Ionizable Amphiphilic Janus Dendrimers in Targeted mRNA Delivery Activity,” J. Am. Chem. Soc., 144 (2022), pp. 4746–4753, 10.1021/jacs.2c00273.

[20] J. Zhang, H. Fan, D. A. Levorse, L. S. Crocker, “Interaction of Cholesterol-Conjugated Ionizable Amino Lipids with Biomembranes: Lipid Polymorphism, Structure-Activity Relationship, and Implications for siRNA Delivery,” Langmuir, 27 (2011), pp. 9473–9483, 10.1021/la201464k.

[21] K. G. M. M. Alberti, P. Zimmet, “Global Burden of Disease--Where Does Diabetes Mellitus Fit In?” Nat. Rev. Endocrinol., 9 (2013), pp. 258–260, 10.1038/nrendo.2013.54.

[22] P. Saeedi, I. Petersohn, P. Salpea, B. Malanda, S. Karuranga, N. Unwin, S. Colagiuri, L. Guariguata, A. A. Motala, K. Ogurtsova, J. E. Shaw, D. Bright, R. Williams, R. Almutairi, P. A. Montoya, A. Basit, S. Besancon, C. Bommer, W. Borgnakke, E. Boyko, “Global and Regional Diabetes Prevalence Estimates for 2019 and Projections for 2030 and 2045: Results from the International Diabetes Federation Diabetes Atlas, 9th Edition,” Diabetes Res. Clin. Pract., 157 (2019), pp. 107843–107853, 10.1016/j.diabres.2019.107843.

[23] T. A. Chowdhury, S. Shaho, A. Moolla, “Complications of Diabetes: Progress, but Significant Challenges Ahead,” Ann. Transl. Med., 2 (2014), pp. 120–124, 10.3978/j.issn.2305-5839.2014.08.12.

[24] P. Saeedi, P. Salpea, S. Karuranga, I. Petersohn, B. Malanda, E. W. Gregg, N. Unwin, S. H. Wild, R. Williams, “Mortality Attributable to Diabetes in 20-79 Years Old Adults, 2019 Estimates: Results from the International Diabetes Federation Diabetes Atlas, 9th Edition,” Diabetes Res. Clin. Pract., 162 (2020), Article 108086, 10.1016/j.diabres.2020.108086.

[25] S. J. Dixon, K. M. Lemberg, M. R. Lamprecht, R. Skouta, E. M. Zaitsev, C. E. Gleason, D. N. Patel, A. J. Bauer, A. M. Cantley, W. S. Yang, B. Morrison, B. R. Stockwell, “Ferroptosis: An Iron-Dependent Form of Nonapoptotic Cell Death,” Cell, 149 (2012), pp. 1060–1072, 10.1016/j.cell.2012.03.042.

[26] S. Altamura, S. Kopf, J. Schmidt, K. Müdder, A. da Silva, P. Nawroth, M. Muckenthaler, “Uncoupled Iron Homeostasis in Type 2 Diabetes Mellitus,” J. Mol. Med., 95 (2017), pp. 1387–1398, 10.1007/s00109-017-1596-3.

[27] M. K. Prasad, S. Mohandas, R. Kunka Mohanram, “Role of Ferroptosis Inhibitors in the Management of Diabetes,” BioFactors, 49 (2023), pp. 270–296, DOI:10.1002/biof.1920.

[28] J. Wang, H. Wang, “Oxidative Stress in Pancreatic Beta Cell Regeneration,” Oxid. Med. Cell. Longev., 2017 (2017), pp. 1930261–1930271, 10.1155/2017/1930261.

[29] V. E. Kagan, G. Mao, F. Qu, J. P. Friedmann Angeli, S. Doll, C. St Croix, H. H. Dar, B. Liu, V. A. Tyurin, V. B. Ritov, A. A. Kapralov, A. A. Amoscato, J. Jiang, T. Anthonymuthu, D. Mohammadyani, Q. Yang, B. Proneth, J. Klein-Seetharaman, S. Watkins, I. Bahar, J. Greenberger, R. K. Mallampalli, B. R. Stockwell, Y. Y. Tyurina, M. Conrad, H. Bayır, “Oxidized Arachidonic and Adrenic PEs Navigate Cells to Ferroptosis,” Nat. Chem. Biol., 13 (2017), pp. 81–90, 10.1038/nchembio.2238.

[30] R. Shintoku, Y. Takigawa, K. Yamada, C. Kubota, Y. Yoshimoto, T. Takeuchi, I. Koshiishi, S. Torii, “Lipoxygenase-Mediated Generation of Lipid Peroxides Enhances Ferroptosis Induced by Erastin and RSL3,” Cancer Sci., 108 (2017), pp. 2187–2194, 10.1111/cas.13380.

[31] S. J. Dixon, G. E. Winter, L. S. Musavi, E.D. Lee, B. Snijder, M. Rebsamen, G. Superti-Furga, B. R. Stockwell, “Human Haploid Cell Genetics Reveals Roles for Lipid Metabolism Genes in Nonapoptotic Cell Death,” ACS Chem. Biol., 10 (2015), pp. 1604–1609, 10.1021/acschembio.5b00245.

[32] S. Doll, B. Proneth, Y. Y. Tyurina, E. Panzilius, S. Kobayashi, I. Ingold, M. Irmler, J. Beckers, M. Aichler, A. Walch, H. Prokisch, D. Trümbach, G. Mao, F. Qu, H. Bayir, J. Füllekrug, C. H. Scheel, W. Wurst, J. A. Schick, V. E. Kagan, J. P. Angeli, M. Conrad, “ACSL4 Dictates Ferroptosis Sensitivity by Shaping Cellular Lipid Composition,” Nat. Chem. Biol., 13 (2017), pp. 91–98, 10.1038/nchembio.2239.

[33] A. Uruno, Y. Furusawa, Y. Yagishita, T. Fukutomi, H. Muramatsu, T. Negishi, A. Sugawara, T. W. Kensler, M. Yamamoto, “The Keap1-Nrf2 System Prevents Onset of Diabetes Mellitus,” Mol. Cell. Biol., 33 (2013), pp. 2996–3010, 10.1128/MCB.00225-13.

[34] R. Pillai, M. Hayashi, A.-M. Zavitsanou, T. Papagiannakopoulos, “NRF2: KEAPing Tumors Protected,” Cancer Discov., 12 (2022), pp. 625–643, 10.1158/2159-8290.CD-21-0922.

[35] W. S. Yang, R. SriRamaratnam, M. E. Welsch, K. Shimada, R. Skouta, V. S. Viswanathan, J. H. Cheah, P. A. Clemons, A. F. Shamji, C. B. Clish, L. M. Brown, A. W. Girotti, V. W. Cornish, S. L. Schreiber, B. R. Stockwell, “Regulation of Ferroptotic Cancer Cell Death by GPX4,” Cell, 156 (2014), pp. 317–331, 10.1016/j.cell.2013.12.010.

[36] Y. Liu, F. He, L. Chen, H. Zhang, J. Xiao, Q. Meng, “Imidazolyl Lipids Enhanced LNP Endosomal Escape for Ferroptosis RNAi Treatment of Cancer,” Small, 20 (2024), Article e2402362, 10.1002/smll.202402362.

[37] P. Patel, N. M. Ibrahim, K. Cheng, “The Importance of Apparent pKa in the Development of Nanoparticles Encapsulating siRNA and mRNA,” Trends Pharmacol. Sci., 42 (2021), pp. 448–460, 10.1016/j.tips.2021.03.002.

[38] J. Zhang, H. Fan, D. A. Levorse, “Ionization Behavior of Amino Lipids for siRNA Delivery: Determination of Ionization Constants, SAR, and the Impact of Lipid pKa on Cationic Lipid-Biomembrane Interactions,” Langmuir, 27 (2011), pp. 1907–1914, 10.1021/la104590k.

[39] J. Gilleron, W. Querbes, A. Zeigerer, A. Borodovsky, G. Marsico, U. Schubert, K. Manygoats, S. Seifert, C. Andree, M. Stöter, H. Epstein-Barash, L. Zhang, V. Koteliansky, K. Fitzgerald, E. Fava, M. Bickle, Y. Kalaidzidis, A. Akinc, M. Maier, M. Zerial, “Image-Based Analysis of Lipid Nanoparticle-Mediated siRNA Delivery, Intracellular Trafficking and Endosomal Escape,” Nat. Biotechnol., 31 (2013), pp. 638–646, 10.1038/nbt.2612.

[40] D. Zhi, S. Zhang, S. Cui, Y. Zhao, Y. Wang, D. Zhao, “The Headgroup Evolution of Cationic Lipids for Gene Delivery,” Bioconjug. Chem., 24 (2013), pp. 487–519, 10.1021/bc300381s.

[41] Y. Sato, K. Hashiba, K. Sasaki, M. Maeki, M. Tokeshi, H. Harashima, “Understanding Structure-Activity Relationships of pH-Sensitive Cationic Lipids Facilitates the Rational Identification of Promising Lipid Nanoparticles for Delivering siRNAs In Vivo,” J. Control. Release, 295 (2019), pp. 140–152, 10.1016/j.jconrel.2019.01.001.

[42] E. Samaridou, J. Heyes, P. Lutwyche, “Lipid Nanoparticles for Nucleic Acid Delivery: Current Perspectives,” Adv. Drug Deliv. Rev., 154 (2020), pp. 37–63, 10.1016/j.addr.2020.06.002.

[43] G. Sahay, W. Querbes, C. Alabi, A. Eltoukhy, S. Sarkar, C. Zurenko, E. Karagiannis, K. Love, D. Chen, R. Zoncu, Y. Buganim, A. Schroeder, R. Langer, D. G. Anderson, “Efficiency of siRNA Delivery by Lipid Nanoparticles Is Limited by Endocytic Recycling,” Nat. Biotechnol., 31 (2013), pp. 653–658, 10.1038/nbt.2614.

[44] J. Rejman, V. Oberle, I. S. Zuhorn, D. Hoekstra, “Size-Dependent Internalization of Particles via the Pathways of Clathrin-and Caveolae-Mediated Endocytosis,” Biochem. J., 377 (2004), pp. 159–169, 10.1042/BJ20031253.

[45] H. Hillaireau, P. Couvreur, “Nanocarriers’ Entry into the Cell: Relevance to Drug Delivery,” Cell. Mol. Life Sci., 66 (2009), pp. 2873–2896, 10.1007/s00018-009-0053-z.

[46] G. S. Chahal, K. J. Helbig, R. G. Parton, E. A. Monson, “The Biology of Endosomal Escape: Strategies for Enhanced Delivery of Therapeutics,” ACS Nano, 20 (2026), pp. 1789–1813, 10.1021/acsnano.5c18112.

[47] S. Cui, J. Ye, “Ferroptosis: The Demise of Cells Through Phospholipid Peroxidation,” Adv. Sci., 13 (2026), Article e24387, 10.1002/advs.202524387.

[48] R. Wang, Y. Chen, “Mechanisms of Ferroptosis in Pancreatic β-Cell Dysfunction in Diabetes and Exploration of Therapeutic Targets,” J. Pharm. Pharmacol., 13 (2026), pp. 1–26, 10.17265/2328-2150/2026.01.001.

[49] Y. Yu, Y. Yao, “Lipid Metabolism in Ferroptosis: Mechanistic Insights and Therapeutic Potential,” Front. Immunol., 16 (2025), Article 1545339, 10.3389/fimmu.2025.1545339.

[50] L. Wu, W. Lai, L. Li, S. Yang, F. Li, C. Yang, X. Gong, L. Wu, “Autophagy Regulates Ferroptosis-Mediated Diabetic Liver Injury by Modulating the Degradation of ACSL4,” J. Diabetes Res., 2024 (2024), Article 7146054, 10.1155/jdr/7146054.

[51] X. Zhou, J. Zhou, Q. Ban, M. Zhang, B. Ban, “Effects of Metformin on the Glucose Regulation, Lipid Levels and Gut Microbiota in High-Fat Diet with Streptozotocin Induced Type 2 Diabetes Mellitus Rats,” Endocrine, 86 (2024), pp. 163–172, 10.1007/s12020-024-03843-y.

[52] M. Salehi, “Alpha-Cell Secretion Across the Spectrum of Glucose Tolerance,” J. Clin. Endocrinol. Metab., 109 (2024), pp. 1456–1457, 10.1210/clinem/dgad686.

[53] R. Singh, M. Gholipourmalekabadi, S. H. Shafikhani, “Animal Models for Type 1 and Type 2 Diabetes,” Front. Endocrinol., 15 (2024), Article 1359685, 10.3389/fendo.2024.1359685.

