## Supplementary material for "Novel Dissymmetric Ionizable Lipid-Assembled Lipid Nanoparticles for Delivery of Ferroptosis-Related siRNA in Diabetic Treatment": Supporting Information.docx

^*^ Corresponding authors:

**CONTENTS**

**1.** Synthesis of dissymmetric four-tailed series lipids…………………………………..…...…………3

**2.** Synthesis of E3-H8 series lipids……..………….…………..……..…………………………..…….4

**3.** Synthesis of D3-H8 series lipids.…..…………………….....……..……………………………...…6

**4.** Synthesis of O18-D3, O12-D3n, O12-D3, O14-D3 and O16-D3 series lipids………………..……..7

**5.** Synthesis of O12-K6 series lipids………….………………………………………..…………..…11

**6.** Synthesis of O12-P6 series lipids………….……………………………..…………….………..…13

**7.** ^1^H NMR, ^13^C NMR and MS data for all synthesized final compounds…..…………………………15

**8.** **FIGURE S1.** Particle size changes of LNPs before and after siNC encapsulation.…………….…..50

**9. FIGURE S2.** Encapsulation efficiency (E.E.) of siNC-loaded LNPs..……………….……..…..…50

**10. FIGURE S3.** Overlaid flow cytometry histograms of intracellular Cy5 fluorescence intensity.....51

**11. FIGURE S4.** *In vivo* fluorescence imaging of 34 dissymmetric hydrophobic-tailed ionizable lipid-based LNPs in mice.…………………………………………………………………………………..52

**12. FIGURE S5.** *Ex vivo* organ fluorescence imaging of 34 dissymmetric hydrophobic-tailed ionizable lipid-based LNPs in mice..……………………………………………………………………………53

**13. FIGURE S6.** Supplementary safety evaluation data of candidate LNPs.……..……...………...…54

**14. FIGURE S7.** Representative confocal laser scanning microscopy (CLSM) micrographs of cellular internalization pathways for O14-LNP (A) and H18a-LNP (B).…..……………………………...…..55

**15. FIGURE S8.** Inhibitory effects of siALOX12-loaded LNPs on high glucose-induced lipid peroxidation in RAW 264.7 macrophages...……………………………………………...………..…56

**16. FIGURE S9.** Target validation and in vivo biocompatibility of siALOX12-loaded LNPs.……………………………………………………………………………….……………...…57

**17. FIGURE S10.** *In vivo* biocompatibility evaluation of siRNA-loaded H18a-LNP formulations.………..………………………………………………………………………………..58

**Chemical Synthesis**

**Synthesis of dissymmetric four-tailed series lipids**

Synthesis of 1,1’-((3-(dioctylamino)propyl)azanediyl)bis(dodecan-2-ol) (C12-D3-A8)

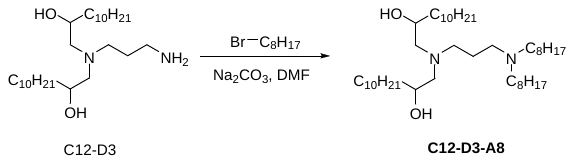

The synthetic method of the intermediate C12-D3 has been reported in our previous work. C12-D3 (300 mg, 0.7 mmol) and Na_2_CO_3_ (574 mg, 5.4 mmol) were dissolved in DMF. The mixture was stirred at room temperature for 2 h, followed by the addition of 1-bromooctane (A8, 327 mg, 1.7 mmol). The resulting reaction system was heated to reflux and maintained for 18 h. After the reaction was confirmed complete by thin-layer chromatography (TLC) monitoring, the crude product was washed three times with saturated sodium chloride solution and then purified *via* column chromatography. Finally, a pale-yellow oily liquid, designated as C12-D3-A8 (0.36 g), was obtained with a yield of 79.5%. ^1^H NMR (600 MHz, CDCl_3_) δ 3.57 (s, 2H), 2.93 (dd, *J* = 10.6, 5.2 Hz, 4H), 2.59 (t, *J* = 14.5 Hz, 2H), 2.51 – 2.42 (m, 2H), 2.30 (d, *J* = 11.9 Hz, 2H), 2.11 (d, *J* = 12.6 Hz, 2H), 1.82 – 1.73 (m, 2H), 1.67 (d, *J* = 9.4 Hz, 4H), 1.46 – 1.36 (m,6H), 1.35 – 1.11 (m, 50H), 0.84 (t, *J* = 7.0 Hz, 12H); ^13^C NMR (150 MHz, CDCl_3_) δ 66.84, 59.45, 50.99, 50.71, 50.02, 34.19, 30.90, 30.64, 28.76, 28.69, 28.61, 28.60, 28.33, 28.03, 28.00, 25.76, 24.78, 21.75, 21.68, 21.56, 13.10, 13.03; MS (MALDI) m/z: C_43_H_90_N_2_O_2_ (M^+^), calcd 667.700; found 667.577 [M+H]^+^.

Synthesis of didodecyl 3,3’-((3-(dioctylamino)propyl)azanediyl)dipropionate (O12-D3-A8)

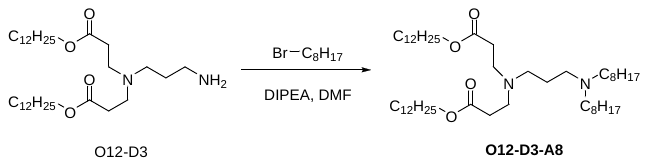

The synthetic method of the intermediate O12-D3 has been reported in our previous work. O12-D3 (300 mg, 0.5 mmol) and N, N-diisopropylethylamine (DIPEA, 312 mg, 2.4 mmol) were dissolved in DMF. The mixture was stirred at room temperature for 2 h, followed by the addition of A8 (234 mg, 1.0 mmol). The reaction was then continued at room temperature for another 18 h. After the reaction completion was verified by TLC monitoring, the crude reaction mixture was washed three times with saturated sodium chloride solution and subsequently purified by column chromatography. Finally, a pale-yellow oily liquid, O12-D3-A8 (80.6 mg), was obtained with a yield of 20.8%. ^1^H NMR (600 MHz, CDCl_3_) δ 7.97 (t, *J* = 5.3 Hz, 1H), 7.53 (s, 1H), 6.81 (s, 1H), 3.64 – 3.45 (m, 1H), 3.26 (q, *J* = 6.4 Hz, 2H), 3.09 – 2.80 (m, 2H), 2.42 (dt, *J* = 39.5, 7.3 Hz, 6H), 1.71 – 1.53 (m, 2H), 1.45 – 1.36 (m, 4H), 1.25 (m, 20H), 0.85 (t, *J* = 7.0 Hz, 6H); ^13^C NMR (150 MHz, CDCl_3_) δ 173.78, 134.13, 54.26, 52.95, 51.11, 37.23, 30.83, 28.55, 28.31, 26.58, 25.56, 21.64, 13.09; MS (MALDI) m/z: C_49_H_98_N_2_O_4_ (M^+^), calcd 779.333; found 779.575 [M+H]^+^.

Synthesis of didodecyl 3,3’-((3-(bis(2-hydroxydodecyl)amino)propyl)azanediyl) dipropionate (O12-D3-C12)

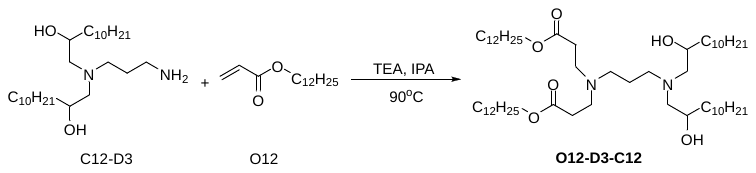

Triethylamine (TEA, 1.45 g, 14.4 mmol) was slowly added dropwise to a solution of C12-D3 (1.00 g, 2.2 mmol) in isopropanol (IPA, 15 mL). After the reaction was stirred for 1 h at room temperature, O12 (3.45 g, 14.4 mmol) was added, and the reaction temperature was raised to 90 °C for reflux reaction for 5 h. After the reaction was confirmed complete by TLC monitoring, the crude reaction mixture was washed three times with saturated sodium chloride solution, and the organic phase was extracted with DCM. Purification by column chromatography afforded a pale-yellow oily liquid, O12-D3-C12 (0.60 g), with a yield of 28.8%. ^1^H NMR (600 MHz, CDCl_3_) δ 4.04 (t, *J* = 6.8 Hz, 4H), 3.76 (s, 2H), 2.79 (t, *J* = 7.0 Hz, 2H), 2.74 (t, *J* = 7.1 Hz, 4H), 2.66 (d, *J* = 11.6 Hz, 2H), 2.58 (dd, *J* = 21.4, 7.4 Hz, 2H), 2.47 (t, *J* = 6.3 Hz, 2H), 2.42 (d, *J* = 7.1 Hz, 4H), 1.77 – 1.65 (m,2H), 1.62 – 1.55 (m, 4H), 1.41 (d, *J* = 9.4 Hz, 4H), 1.30 – 1.21 (m, 68H), 0.91 – 0.82 (m, 12H); ^13^C NMR (150 MHz, CDCl_3_) δ 171.65, 67.72, 63.76, 61.53, 53.30, 49.89, 47.79, 34.05, 34.00, 30.96, 30.92, 28.76, 28.72, 28.67, 28.64, 28.62, 28.36, 28.31, 27.63, 24.94, 24.61, 21.69, 13.11; MS (MALDI) m/z: C_57_H_114_N_2_O_6_ (M^+^), calcd 923.868; found 923.572 [M+H]^+^.

**Synthesis of E3-H8 series lipids**

Synthesis of 3-(dioctylamino)propyl octanoate (A8-E3-H8)

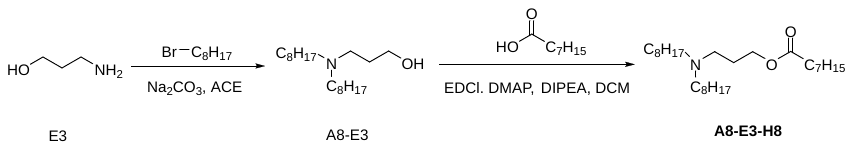

The synthetic method of the intermediate A8-E3 was analogous to that of C12-D3-A8, with the only difference being the use of acetonitrile (ACE) as the solvent. Subsequently, A8-E3 (500 mg, 1.7 mmol), octanoic acid (H8, 722 mg, 5.0 mmol), 4-dimethylaminopyridine (DMAP) (224 mg, 1.8 mmol), 1-ethyl-3-(3-dimethyl aminopropyl) carbodiimide (EDCI) (351 mg, 1.8 mmol) and DIPEA (237 mg, 1.8 mmol) were dissolved in DCM (20 mL). The reaction mixture was heated to 50 °C under reflux for 28 h. After the reaction was confirmed complete by TLC monitoring, the mixture was subjected to reduced pressure distillation, washed three times with saturated sodium bicarbonate solution, and extracted with DCM. Subsequent purification *via* column chromatography afforded a pale-yellow viscous liquid, A8-E3-H8 (570 mg), with a yield of 80.2%. ^1^H NMR (600 MHz, CDCl_3_) δ 4.03 (t, *J* = 6.5 Hz, 2H), 2.46 – 2.38 (m, 2H), 2.36 – 2.26 (m, 4H), 2.22 (t, *J* = 7.6 Hz, 2H), 1.68 (p, *J* = 6.6 Hz, 2H), 1.55 (p, *J* = 7.6 Hz, 2H), 1.34 (p, *J* = 7.2 Hz, 4H), 1.27 – 1.11 (m, 28H), 0.81 (t, *J* = 7.0 Hz, 9H); ^13^C NMR (150 MHz, CDCl_3_) δ 172.87, 61.75, 53.18, 49.45, 33.39, 30.88, 30.68, 28.60, 28.34, 28.15, 27.96, 26.57, 26.07, 25.42, 24.03, 21.67, 21.61, 13.09; MS (ESI) m/z: C_27_H_55_NO_2_ (M^+^), calcd 426.42; found 426.43 [M+H]^+^.

Synthesis of 3-(bis(2-hydroxydodecyl)amino)propyl octanoate (C12-E3-H8)

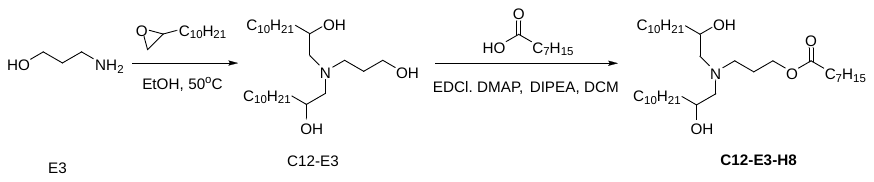

3-Aminopropan-1-ol (E3, 1.0 g, 13.3 mmol) and 2-decyloxirane (C12, 5.4 g, 29.3 mmol) were dissolved in EtOH (15 mL). The reaction mixture was stirred at 50 °C overnight. After the reaction was confirmed complete by TLC monitoring, the mixture was subjected to reduced pressure distillation, followed by purification *via* column chromatography to afford a white solid, C12-E3 (570 mg), with a yield of 84.7%. Subsequently, C12-E3-H8 was synthesized from C12-E3 (100 mg, 0.2 mmol) using a procedure analogous to that described for the synthesis of A8-E3-H8. A pale-yellow oily liquid, C12-E3-H8 (53 mg), was obtained with a yield of 41.6%. ^1^H NMR (600 MHz, CDCl_3_) δ 4.14 – 4.06 (m, 2H), 3.68 – 3.55 (m, 2H), 2.70 – 2.62 (m, 2H), 2.56 (dd, *J* = 13.2, 3.4 Hz, 2H), 2.40 (t, *J* = 5.4 Hz, 1H), 2.29 (t, *J* = 8.0 Hz, 2H), 1.80 (p, *J* = 6.4 Hz, 2H), 1.61 (p, *J* = 7.5 Hz, 2H), 1.48 – 1.35 (m, 6H), 1.33 – 1.20 (m, 40H), 0.87 (t, *J* = 7.0 Hz, 9H); ^13^C NMR (150 MHz, CDCl_3_) δ 172.88, 68.53, 66.79, 61.63, 61.32, 60.08, 51.68, 50.71, 34.06, 33.86, 33.29, 30.90, 30.66, 28.76, 28.61, 28.60, 28.12, 27.93, 25.35, 24.64, 23.95, 21.68, 21.59, 13.10; MS (ESI) m/z: C_35_H_71_NO_4_ (M^+^), calcd 570.54; found 570.54 [M+H]^+^.

Synthesis of didodecyl 3,3’-((3-(octanoyloxy)propyl)azanediyl)dipropionate (O12-E3-H8)

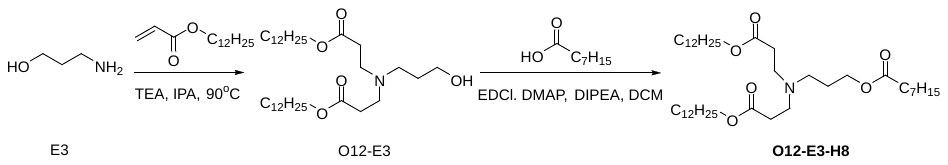

The synthetic method of the intermediate O12-E3 was analogous to that of O12-D3-C12. Subsequently, O12-E3-H8 was synthesized from O12-E3 (500 mg, 0.2 mmol) following a procedure analogous to that employed for A8-E3-H8. A pale-yellow oily liquid, O12-E3-H8 (300 mg), was obtained with a yield of 48.8%. ^1^H NMR (600 MHz, CDCl_3_) δ 4.14 – 3.96 (m, 6H), 2.74 (t, *J* = 7.0 Hz, 4H), 2.47 (t, *J* = 6.8 Hz, 2H), 2.40 (t, *J* = 7.0 Hz, 4H), 2.27 (t, *J* = 7.6 Hz, 2H), 1.73 (p, *J* = 6.5 Hz, 2H), 1.59 (p, *J* = 6.8 Hz, 6H), 1.34 – 1.12 (m, 44H), 0.86 (t, *J* = 7.0 Hz, 9H); ^13^C NMR (150 MHz, CDCl_3_) δ 172.83, 171.63, 63.63, 61.28, 52.41, 49.14, 48.28, 33.34, 31.71, 30.91, 30.67, 28.66, 28.64, 28.29, 27.94, 27.63, 25.57, 24.94, 24.00, 21.68, 21.60, 13.10; MS (ESI) m/z: C_41_H_79_NO_6_ (M^+^), calcd 682.59; found 682.57 [M+H]^+^.

**Synthesis of D3-H8 series lipids**

Synthesis of A8-D3-H8, C12-D3-H8 and O12-D3-H8

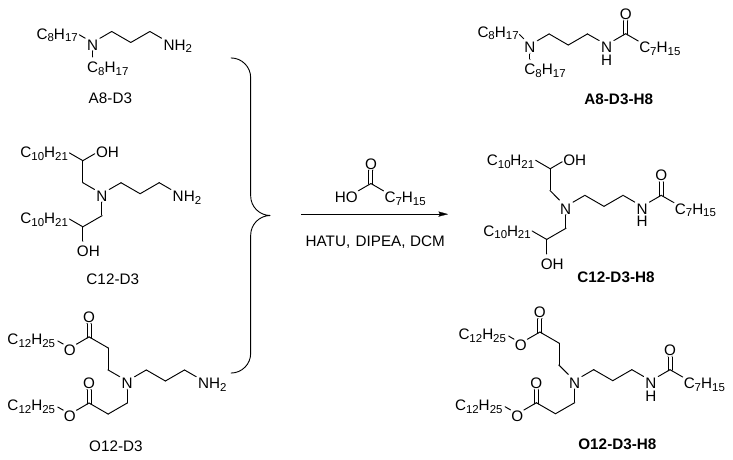

The synthetic methods of the intermediates A8-D3, C12-D3 and O12-D3 have been reported in our previous work. A8-D3 (400 mg, 1.3 mmol), C12-D3 (500 mg, 1.1 mmol) or O12-D3 (555 mg, 1.0 mmol), H8 (579 mg, 4.0 mmol), O-(7-azabenzotriazol-1-yl)-N, N, N’, N’-tetramethyluronium hexafluorophosphate (HATU, 1.0 g, 2.7 mmol) and DIPEA (692 mg, 5.37 mmol) were dissolved in DCM (15 mL). The mixture was stirred at room temperature for 18 h. After the reaction was confirmed complete by TLC monitoring, the solvent was removed by reduced pressure distillation. The residue was washed three times with saturated sodium chloride solution and extracted with DCM. Purification *via* column chromatography afforded pale-yellow oily liquids, namely A8-D3-H8 (200 mg, 42.6% yield), C12-D3-H8 (130 mg, 32.5% yield) and O12-D3-H8 (132 mg, 44.1% yield), respectively.

A8-D3-H8 (N-(3-(dioctylamino)propyl)octanamide): ^1^H NMR (600 MHz, CDCl_3_) δ 6.99 (t, *J* = 5.8 Hz, 1H), 3.36 – 3.27 (m, 2H), 3.10 (d, *J* = 5.5 Hz, 2H), 3.03 (d, *J* = 8.3 Hz, 4H), 2.31 – 2.25 (m, 2H), 2.02 (s, 2H), 1.68 (p, *J* = 7.6 Hz, 4H), 1.65 – 1.55 (m, 2H), 1.39 – 1.22 (m, 30H), 0.88 (t, *J* = 7.0 Hz, 9H); ^13^C NMR (150 MHz, CDCl_3_) δ 178.68, 52.84, 50.95, 39.17, 37.00, 35.10, 34.05, 31.67, 31.66, 29.15, 29.08, 29.02, 29.01, 28.95, 28.88, 26.58, 25.96, 24.80, 23.69, 23.42, 22.61, 22.59, 14.07, 14.04; MS (ESI) m/z: C_27_H_56_N_2_O (M^+^), calcd 425.439; found 425.447 [M+H]^+^.

C12-D3-H8 (N-(3-(bis(2-hydroxydodecyl)amino)propyl)octanamide): ^1^H NMR (600 MHz, CDCl_3_) δ 6.69 (s, 1H), 3.84 (d, *J* = 7.5 Hz, 2H), 3.28 (q, *J* = 6.2 Hz, 2H), 2.98 (t, *J* = 7.1 Hz, 2H), 2.78 (d, *J* = 8.5 Hz, 4H), 2.16 – 2.06 (m, 2H), 1.83 (p, *J* = 6.6 Hz, 2H), 1.60 – 1.51 (m, 2H), 1.45 – 1.28 (m, 6H), 1.27 – 1.13 (m, 40H), 0.89 – 0.71 (m, 9H); ^13^C NMR (150 MHz, CDCl_3_) δ 173.27, 66.83, 61.23, 52.80, 35.67, 35.65, 34.21, 33.99, 30.91, 30.73, 28.65, 28.62, 28.34, 28.09, 24.82, 24.48, 21.68, 21.63, 13.11; MS (ESI) m/z: C_35_H_72_N_2_O_3_ (M^+^), calcd 569.554; found 569.562 [M+H]^+^.

O12-D3-H8 (didodecyl 3,3’-((3-octanamidopropyl)azanediyl)dipropionate): ^1^H NMR (600 MHz, CDCl_3_) δ 6.60 (s, 1H), 4.05 (t, *J* = 6.8 Hz, 4H), 3.25 (q, *J* = 5.9 Hz, 2H), 2.78 (s, 4H), 2.49 (d, *J* = 22.3 Hz, 3H), 2.24 – 2.11 (m, 2H), 1.61 (dt, *J* = 14.2, 6.4 Hz, 8H), 1.37 – 1.17 (m, 44H), 0.87 (t, *J* = 7.0 Hz, 9H); ^13^C NMR (150 MHz, CDCl_3_) δ 176.08, 172.54, 171.59, 63.92, 50.02, 48.09, 37.60, 35.78, 30.91, 30.72, 28.69, 28.65, 28.63, 28.59, 28.53, 28.34, 28.28, 28.06, 27.60, 24.92, 24.88, 21.68, 21.62; MS (ESI) m/z: C_41_H_80_N_2_O_5_ (M^+^), calcd 681.61; found 681.62 [M+H]^+^.

**Synthesis of O18-D3, O12-D3n, O12-D3, O14-D3 and O16-D3 series lipids**

Synthesis of O18-D3-H7, O18-D3-H8, O18-D3-H9, O12-D3-H18a, O12-D3-H18b, O12-D3-H18c, O12-D3-H9, O12-D3-H16, O12-D3-H17, O12-D3-H18, O14-D3-H9, O14-D3-H13, O14-D3-H14, O14-D3-H15, O16-D3-H9, O16-D3-H10, O16-D3-H11 and O16-D3-H12

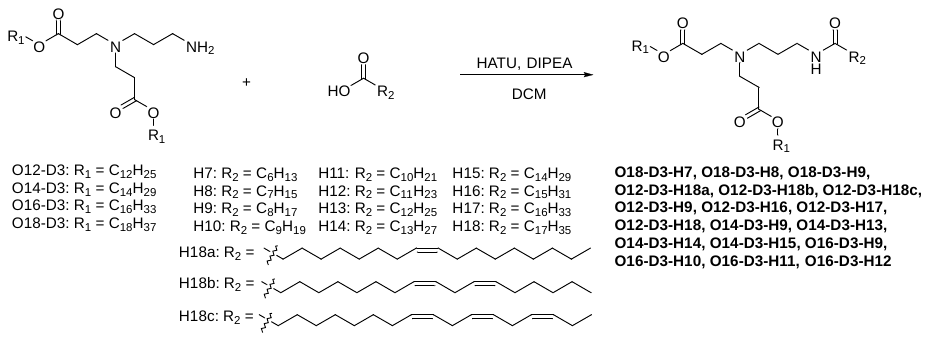

The synthetic routes of the intermediates O14-D3, O16-D3 and O18-D3 are analogous to that of O12-D3, all of which were prepared according to our previously reported procedure. The synthesis of a total of 18 ionizable lipids, including the O18-D3, O12-D3n, O12-D3, O14-D3 and O16-D3 series, was performed following the synthetic protocol established for O12-D3-H8.

O18-D3-H7 (dioctadecyl 3,3’-((3-heptanamidopropyl)azanediyl)dipropionate): Off white solid, 59.0% yield. ^1^H NMR (600 MHz, CDCl_3_) δ 6.56 (t, *J* = 5.8 Hz, 1H), 4.06 (t, *J* = 6.8 Hz, 4H), 3.26 (q, *J* = 6.1 Hz, 2H), 2.80 (s, 4H), 2.48 (s, 6H), 2.17 (t, *J* = 7.7 Hz, 2H), 1.62 (dt, *J* = 14.5, 6.9 Hz, 8H), 1.32-1.25 (m, 66H), 0.87 (t, *J* = 7.0 Hz, 9H); ^13^C NMR (150 MHz, CDCl_3_) δ172.81,172.71,65.12,51.28, 49.33, 37.45,36.87, 32.06, 31.72, 29.84, 29.80, 29.75, 29.69, 29.50, 29.43, 29.21, 28.74, 26.07, 25.95, 22.82, 22.68, 14.25, 14.18; MS (MALDI) m/z calcd for C_52_H_102_N_2_O_5_ (M^+^) 835.782; found 835.766 [M+H]^+^, 857.740 [M+Na]^+^.

O18-D3-H8 (dioctadecyl 3,3’-((3-octanamidopropyl)azanediyl)dipropionate): Off white solid, 53.2% yield. ^1^H NMR (600 MHz, CDCl_3_) δ 6.65 (t, *J* = 5.7 Hz, 1H), 4.02 (t, *J* = 6.8 Hz, 4H), 3.21 (q, *J* = 6.0 Hz, 2H), 2.73 (t, *J* = 6.9 Hz, 4H), 2.44 (dt, *J* = 27.7, 6.5 Hz, 6H), 2.18 – 2.11 (m, 2H), 1.64 – 1.56 (m, 8H), 1.30 – 1.18 (m, 68H), 0.86 – 0.82 (m, 9H); ^13^C NMR (150 MHz, CDCl_3_) δ172.87, 171.93, 64.10, 50.28, 48.41, 36.75, 36.00, 31.53, 31.20, 31.01, 28.98, 28.94, 28.88, 28.82, 28.64, 28.57, 28.33, 27.89, 25.21, 25.16,24.27, 21.96, 21.89, 13.36, 13.32; MS (ESI) m/z calcd for C_53_H_104_N_2_O_5_ (M^+^) 849.403; found 849.800 [M+H]^+^, 871.850[M+Na]^+^.

O18-D3-H9 (dioctadecyl 3,3’-((3-nonanamidopropyl)azanediyl)dipropionate): Off white solid, 43.8% yield. ^1^H NMR (600 MHz, CDCl_3_) δ 6.63 (t, *J* = 5.8 Hz, 1H), 4.04 (t, *J* = 6.8 Hz, 4H), 3.25 (q, *J* = 6.0 Hz, 2H), 2.81 (t, *J* = 7.2 Hz, 4H), 2.50 (dt, *J* = 44.6, 5.6 Hz, 6H), 2.20 – 2.12 (m, 2H), 1.72 – 1.65 (m, 2H), 1.59 (q, *J* = 7.0 Hz, 6H), 1.32 – 1.21 (m, 70H), 0.86 (t, *J* = 7.0 Hz, 9H); ^13^C NMR (150 MHz, CDCl_3_) δ 174.19, 172.82, 65.28, 51.43, 49.49, 37.55, 37.00, 32.24, 32.16,32.14, 30.02, 29.98, 29.92, 29.86, 29.72, 29.67, 29.60, 29.52, 28.91, 26.24, 26.17, 26.11, 22.99, 22.96, 14.41,14.39; MS (ESI) m/z calcd for C_54_H_106_N_2_O_5_ (M^+^) 863.430; found 863.750 [M+H]^+^, 885.900 [M+Na]^+^.

O12-D3-H18a (didodecyl 3,3’-((3-oleamidopropyl)azanediyl)dipropionate): Off white solid, 23.5% yield. ^1^H NMR (600 MHz, CDCl_3_) δ 6.54 (t, *J* = 5.7 Hz, 1H), 5.36 – 5.30 (m, 2H), 4.05 (t, *J* = 6.9 Hz, 4H), 3.24 (q, *J* = 6.0 Hz, 2H), 2.75 (t, *J* = 7.0 Hz, 4H), 2.45 (dt, *J* = 21.6, 6.7 Hz, 6H), 2.18 – 2.14 (m, 2H), 2.00 (q, *J* = 7.0 Hz, 4H), 1.61 (dt, *J* = 14.8, 6.7 Hz, 8H), 1.30 – 1.24 (m, 56H), 0.87 (t, *J* = 7.1 Hz, 9H); ^13^C NMR (150 MHz, CDCl_3_) δ 173.70,173.12, 130.31, 130.13, 65.20, 51.35, 49.55, 37.85, 37.15, 32.27, 30.12, 30.05, 30.01, 29.99, 29.95, 29.90, 29.88, 29.76, 29.70, 29.67, 29.64, 29.60,29.57, 28.97, 27.57, 26.29, 26.24, 23.04, 14.46; MS (ESI) m/z calcd for C_51_H_98_N_2_O_5_ (M^+^) 818.748; found 819.850 [M+H]^+^.

O12-D3-H18b (didodecyl 3,3’-((3-((9*Z*,12*Z*)-octadeca-9,12-dienamido)propyl) azanediyl)dipropionate): Off white solid, 56.2% yield. ^1^H NMR (600 MHz, CDCl_3_) δ 6.53 (t, *J* = 5.8 Hz, 1H), 5.38 – 5.29 (m, 4H), 4.05 (t, *J* = 6.8 Hz, 4H), 3.24 (q, *J* = 6.0 Hz, 2H), 2.75 (dt, *J* = 12.8, 6.9 Hz, 6H), 2.45 (dt, *J* = 21.6, 6.5 Hz, 6H), 2.16 (t, *J* = 7.7 Hz, 2H), 2.04 (q, *J* = 7.2 Hz, 4H), 1.62 (dp, *J* = 14.0, 6.5 Hz, 8H), 1.37– 1.25(m, 50H), 0.88 (q, *J* = 7.1, 6.6 Hz, 9H); ^13^C NMR (150 MHz, CDCl_3_) δ 172.71, 130.14, 130.00, 127.93, 127.84, 64.75, 50.90, 49.13, 37.42, 36.72, 32.40, 31.84, 31.44, 29.58, 29.55, 29.52, 29.47, 29.32, 29.27, 29.21, 29.13, 28.54, 27.14, 27.12, 25.92, 25.86, 25.80, 25.54, 22.60, 22.49, 14.03, 13.98; MS (ESI) m/z calcd for C_51_H_96_N_2_O_5_ (M^+^) 816.732; found 817.800 [M+H]^+^, 839.850 [M+Na]^+^.

O12-D3-H18c (didodecyl 3,3’-((3-((9*Z*,12*Z*,15*Z*)-octadeca-9,12,15-trienamido) propyl)azanediyl)dipropionate): Off white solid, 25.4% yield. ^1^H NMR (600 MHz, CDCl_3_) δ 6.53 (t, *J* = 5.9 Hz, 1H), 5.59 – 5.13 (m, 6H), 4.06 (t, *J* = 6.8 Hz, 4H), 3.26 (q, *J* = 6.1 Hz, 2H), 2.83 – 2.75 (m, 8H), 2.47 (s, 6H), 2.17 (t, *J* = 7.7 Hz, 2H), 2.05 (dq, *J* = 14.2, 7.3 Hz, 4H), 1.82 – 1.49 (m, 10H), 1.42 – 1.25 (m, 44H), 0.97 (t, *J* = 7.5 Hz, 3H), 0.87 (t, *J* = 6.9 Hz, 6H); ^13^C NMR (150 MHz, CDCl_3_) δ 172.98, 172.91, 132.31, 130.64, 128.64, 128.05, 127.47,65.24, 51.43, 49.57, 38.95, 37.07, 31.87, 30.48, 30.01, 29.99, 29.95, 29.89, 29.74, 29.70, 29.64, 29.56, 28.95, 27.58, 27.54, 26.28, 26.19, 25.96, 25.87, 23.03,20.90, 14.62, 14.46; MS (ESI) m/z calcd for C_51_H_94_N_2_O_5_ (M^+^) 814.716; found 815.850 [M+H]^+^, 837.800 [M+Na]^+^.

O12-D3-H9 (didodecyl 3,3’-((3-nonanamidopropyl)azanediyl)dipropionate): Off white solid, 42.6% yield. ^1^H NMR (600 MHz, CDCl_3_) δ ppm 6.57 (t, *J =* 5.7 Hz, 1H), 4.05 (t, *J =* 6.8 Hz, 4H), 3.25 (q, *J =* 6.0 Hz, 2H), 2.77 (d, *J =* 7.1 Hz, 4H), 2.50 (s, 2H), 2.45 (t, *J =* 6.9 Hz, 4H), 2.20 – 2.14 (m, 2H), 1.67 (t, *J =* 6.2 Hz, 2H), 1.63 – 1.59 (m, 6H), 1.32 – 1.24 (m, 46H), 0.89 – 0.86 (m, 9H); ^13^C NMR (150 MHz, CDCl_3_) δ ppm 173.97, 173.02, 65.25, 51.40, 49.50, 38.96, 37.12, 32.26, 32.20, 32.15, 30.00, 29.98, 29.95, 29.89, 29.77, 29.70, 29.63, 29.58, 29.55, 29.48, 29.46, 28.96, 26.28, 26.23, 25.18, 23.03, 22.98, 14.45; MS (MALDI) *m/z* calcd for C_42_H_82_N_2_O_5_ (M)^+^ 694.626; found 695.557 [M+H]^+^, 717.542 [M+Na]^+^, 733.529 [M+K]^+^.

O12-D3-H16 (didodecyl 3,3'-((3-palmitamidopropyl)azanediyl)dipropionate): Off white solid, 39.6% yield. ^1^H NMR (600 MHz, CDCl_3_) δ ppm 6.56 (t, *J =* 5.9 Hz, 1H), 4.07 (t, *J =* 6.8 Hz, 4H), 3.28 (s, 2H), 2.80 (s, 4H), 2.51 (s, 4H), 2.34 (t, *J =* 7.5 Hz, 2H), 2.18 (t, *J =* 7.7 Hz, 2H), 1.65 – 1.65 (m, 8H), 1.30 – 1.25 (m, 60H), 0.88 (t, *J =* 7.0 Hz, 9H); ^13^C NMR (150 MHz, CDCl_3_) δ ppm 178.52, 172.19, 65.03, 51.04, 48.98, 31.83, 29.63, 29.59, 29.57, 29.55, 29.52, 29.48, 29.46, 29.33, 29.27, 29.20, 28.48, 25.82, 25.74, 24.75, 14.01; MS (MALDI) *m/z* calcd for C_49_H_96_N_2_O_5_ (M^+^) 792.735; found 793.578 [M+H]^+^, 815.870 [M+Na]^+^.

O12-D3-H17 (didodecyl 3,3’-((3-heptadecanamidopropyl)azanediyl)dipropionate): Off white solid, 36.6% yield. ^1^H NMR (600 MHz, CDCl_3_) δ ppm 6.58 (t, *J =* 5.7 Hz, 1H), 4.05 (t, *J =* 6.9 Hz, 4H), 3.25 (q, *J =* 6.0 Hz, 2H), 2.76 (t, *J =* 7.0 Hz, 4H), 2.47 (dt, *J =* 26.4, 6.6 Hz, 6H), 2.17 (t, *J =* 7.7 Hz, 2H), 1.66 (t, *J =* 6.5 Hz, 2H), 1.62 (q, *J =* 6.9 Hz, 6H), 1.29 – 1.24 (m, 62H), 0.87 (t, *J =* 7.0 Hz, 9H); ^13^C NMR (150 MHz, CDCl_3_) δ ppm 178.12, 172.63, 64.81, 50.94, 49.06, 37.44, 36.72, 31.85, 29.64, 29.63, 29.59, 29.57, 29.53, 29.50, 29.48, 29.39, 29.36, 29.29, 29.22, 29.05, 28.54, 25.86, 25.83, 24.74, 22.61, 14.04; MS (MALDI) *m/z* calcd for C_50_H_98_N_2_O_5_ (M^+^) 806.751; found 807.696 [M+H]^+^, 829.679 [M+Na]^+^.

O12-D3-H18 (didodecyl 3,3’-((3-stearamidopropyl)azanediyl)dipropionate): Off white solid, 48.5% yield. ^1^H NMR (600 MHz, CDCl_3_) δ ppm 6.66 (t, *J =* 6.0 Hz, 1H), 4.06 (t, *J =* 7.1 Hz, 4H), 3.27 (t, *J =* 6.6 Hz, 2H), 2.86 (s, 4H), 2.52 (t, *J =* 7.0 Hz, 4H), 2.32 (t, *J =* 7.7 Hz, 2H), 2.18 (t, *J =* 7.7 Hz, 2H), 1.75 – 1.69 (m, 2H), 1.61 (s, 6H), 1.32 – 1.25 (m, 64H), 0.86 (t, *J =* 7.4 Hz, 9H); ^13^C NMR (150 MHz, CDCl_3_) δ ppm 174.21, 172.29, 64.99, 50.99, 48.97, 37.11, 36.59, 34.07, 31.84, 29.64, 29.61, 29.58, 29.53, 29.47, 29.39, 29.34, 29.28, 29.21, 29.05, 28.50, 25.83, 25.78, 25.51, 24.73, 22.61, 14.02; MS (ESI) *m/z* calcd for C_51_H_100_N_2_O_5_ (M^+^) 820.767; found 821.850 [M+H]^+^.

O14-D3-H9 (ditetradecyl 3,3’-((3-nonanamidopropyl)azanediyl)dipropionate): Off white solid, 38.6% yield. ^1^H NMR (600 MHz, CDCl_3_) δ ppm 6.58 (s, 1H), 4.06 (t, *J =* 6.8 Hz, 4H), 3.26 (q, *J =* 6.1 Hz, 2H), 2.76 (s, 4H), 2.46 (s, 4H), 2.34 (t, *J =* 7.5 Hz, 2H), 2.17 (t, *J =* 7.7 Hz, 2H), 1.64 – 1.58 (m, 8H), 1.33 –1.26 (m, 54H), 0.89 – 0.87 (m, 9H); ^13^C NMR (150 MHz, CDCl_3_) δ ppm 65.27, 51.39, 49.47, 37.16, 32.28, 32.21, 32.16, 31.86, 31.79, 30.05, 30.02, 29.96, 29.91, 29.78, 29.72, 29.65, 29.56, 29.46, 28.97, 26.29, 26.25, 25.11, 23.04, 22.99, 14.47; MS (ESI) m/z calcd for C_46_H_90_N_2_O_5_ (M^+^) 751.692; found: 752.700 [M+H]^+^, 773.750 [M+Na]^+^.

O14-D3-H13 (ditetradecyl 3,3’-((3-tridecanamidopropyl)azanediyl)dipropionate): Off white solid, 17.6% yield. ^1^H NMR (600 MHz, CDCl_3_) δ ppm 6.57 (t, *J =* 6.0 Hz, 1H), 4.07 (t, *J =* 6.8 Hz, 4H), 3.29 (q, *J =* 8.7 Hz, 2H), 2.89 (s, 6H), 2.53 (d, *J =* 9.9 Hz, 4H), 2.17 (t, *J =* 7.7 Hz, 2H), 1.74 (s, 2H), 1.64 – 1.58 (m, 6H), 1.33 – 1.23 (m, 62H), 0.87 (t, *J =* 7.0 Hz, 9H); ^13^C NMR (150 MHz, CDCl_3_) δ ppm 172.77, 172.60, 65.51, 51.63, 49.56, 36.98, 32.27, 30.04, 30.01, 29.96, 29.90, 29.77, 29.71, 29.64, 28.92, 26.26, 26.15, 23.03, 14.45; MS (MALDI) *m/z* calcd for C_50_H_98_N_2_O_5_ (M^+^) 806.751; found: 807.650 [M+H]^+^.

O14-D3-H14 (ditetradecyl 3,3’-((3-tetradecanamidopropyl)azanediyl)dipropionate): Off white solid, 17.3% yield. ^1^H NMR (600 MHz, CDCl_3_) δ ppm 6.58 (t, *J =* 6.5 Hz, 1H), 4.25 – 3.99 (m, 4H), 3.46 – 3.38 (m, 4H), 3.17 (s, 2H), 2.97 – 2.71 (m, 4H), 2.22 (t, *J =* 7.8 Hz, 2H), 2.10 – 1.96 (m, 2H), 1.83 – 1.43 (m, 8H), 1.32 – 1.22 (m, 64H), 0.88 (t, *J =* 7.0 Hz, 9H); ^13^C NMR (150 MHz, CDCl_3_) δ ppm 176.86, 170.72, 66.00, 51.83, 49.12, 35.99, 31.86, 29.66, 29.64, 29.61, 29.60, 29.57, 29.50, 29.49, 29.31, 29.30, 29.23, 28.37, 25.77, 25.45, 22.62, 14.04; MS (MALDI) *m/z* calcd for C_51_H_100_N_2_O_5_ (M^+^) 820.767; found: 821.644 [M+H]^+^.

O14-D3-H15 (ditetradecyl 3,3’-((3-pentadecanamidopropyl)azanediyl)dipropionate): Off white solid, 23.1% yield. ^1^H NMR (600 MHz, CDCl_3_) δ ppm 6.60 (t, *J =* 5.7 Hz, 1H), 4.05 (t, *J =* 6.9 Hz, 4H), 3.26 (q, *J =* 6.0 Hz, 2H), 2.81 (s, 4H), 2.51 (dt, *J =* 41.2, 6.9 Hz, 6H), 2.17 (t, *J =* 7.7 Hz, 2H), 1.70 – 1.59 (m, 8H), 1.29 – 1.24 (m, 66H), 0.86 (t, *J =* 7.1 Hz, 9H); ^13^C NMR (150 MHz, CDCl_3_) δ ppm 173.71, 172.46, 64.89, 51.02, 49.07, 37.24, 36.66, 33.74, 31.85, 29.62, 29.59, 29.53, 29.48, 29.41, 29.35, 29.28, 29.22, 29.08, 28.53, 25.85, 25.80, 25.71, 22.61, 14.03; MS (ESI) *m/z* calcd for C_52_H_102_N_2_O_5_ (M^+^) 834.377; found: 835.850 [M+H]^+^, 857.900 [M+Na]^+^.

O16-D3-H9 (dihexadecyl 3,3’-((3-nonanamidopropyl)azanediyl)dipropionate): Off white solid, 66.4% yield. ^1^H NMR (600 MHz, CDCl_3_) δ ppm 6.59 (t, *J =* 6.6 Hz, 1H), 4.13 (t, *J =* 6.6 Hz, 4H), 3.46 – 3.38 (m, 4H), 3.16 (s, 2H), 2.82 (d, *J =* 19.2 Hz, 4H), 2.26 – 2.20 (m, 2H), 2.04 (q, *J =* 7.1, 6.1 Hz, 2H), 1.61 (dq, *J =* 30.0, 8.1 Hz, 8H), 1.34 – 1.20 (m, 62H), 0.88 (d, *J =* 7.0 Hz, 9H); ^13^C NMR (150 MHz, CDCl_3_) δ ppm 166.21, 166.13, 66.39, 49.54, 38.96, 34.15, 32.27, 32.18, 30.05, 30.02, 29.98, 29.89, 29.71, 29.64, 29.58, 29.53, 29.48, 28.80, 26.19, 25.89, 25.20, 23.03, 22.99, 14.44, 14.43; MS (MALDI) *m/z* calcd for C_50_H_98_N_2_O_5_ (M^+^) 806.324; found: 807.387 [M+H]^+^.

O16-D3-H10 (dihexadecyl 3,3’-((3-decanamidopropyl)azanediyl)dipropionate): Off white solid, 63.7% yield. ^1^H NMR (600 MHz, CDCl_3_) δ ppm 6.73 (t, *J =* 6.5 Hz, 1H), 4.11 (t, *J =* 7.3 Hz, 4H), 3.39 (q, *J =* 6.6 Hz, 4H), 3.14 (t, *J =* 6.3 Hz, 2H), 2.82 (s, 4H), 2.20 (t, *J =* 7.8 Hz, 2H), 2.01 (p, *J =* 6.1 Hz, 2H), 1.64 – 1.53 (m, 6H), 1.31 – 1.24 (m, 66H), 0.86 (t, *J =* 7.0 Hz, 9H); ^13^C NMR (150 MHz, CDCl_3_) δ ppm 170.73, 165.65, 65.84, 51.82, 49.08, 38.47, 35.87, 31.80, 31.76, 29.59, 29.58, 29.54, 29.45, 29.38, 29.24, 29.20, 28.32, 25.72, 25.39, 22.55, 13.97; MS (MALDI) *m/z* calcd for C_51_H_100_N_2_O_5_ (M^+^) 820.320; found: 821.500 [M+H]^+^.

O16-D3-H11 (dihexadecyl 3,3’-((3-undecanamidopropyl)azanediyl)dipropionate): Off white solid, 67.6% yield. ^1^H NMR (600 MHz, CDCl_3_) δ ppm 6.59 (t, *J =* 6.5 Hz, 1H), 4.17 – 4.08 (m, 4H), 3.42 (t, *J =* 6.3 Hz, 4H), 3.19 – 3.15 (m, 2H), 2.83 (t, *J =* 15.7 Hz, 4H), 2.29 – 2.16 (m, 2H), 2.04 (p, *J =* 5.9 Hz, 2H), 1.66 – 1.54 (m, 6H), 1.43 (dd, *J =* 17.9, 6.7 Hz, 2H), 1.37 – 1.25 (m, 66H), 0.87 (t, *J =* 7.2 Hz, 9H); ^13^C NMR (150 MHz, CDCl_3_) δ ppm 176.0, 169.93, 65.23, 54.88, 51.07, 48.35, 42.78, 35.17, 34.08, 31.06, 28.85, 28.83, 28.81, 28.78, 28.70, 28.50, 28.44, 27.57, 24.98, 24.65, 23.93, 21.82, 13.25; MS (MALDI) *m/z* calcd for C_52_H_102_N_2_O_5_ (M^+^) 834.377; found: 835.519 [M+H]^+^.

O16-D3-H12 (dihexadecyl 3,3’-((3-dodecanamidopropyl)azanediyl)dipropionate): Off white solid, 63.2% yield. ^1^H NMR (600 MHz, CDCl_3_) δ ppm 6.55 (t, *J =* 5.8 Hz, 1H), 4.05 (t, *J =* 6.9 Hz, 4H), 3.25 (q, *J =* 6.0 Hz, 2H), 2.77 (s, 4H), 2.58 – 2.34 (m, 6H), 2.19 – 2.14 (m, 2H), 1.63 (tq, *J =* 14.2, 6.9, 6.5 Hz, 8H), 1.33 – 1.25 (m, 68H), 0.87 (t, *J =* 6.9 Hz, 9H); ^13^C NMR (150 MHz, CDCl_3_) δ ppm 173.80, 173.04, 65.25, 51.45, 49.57, 37.15, 32.28, 30.05, 30.02, 29.97, 29.91, 29.79, 29.71, 29.65, 28.97, 26.29, 26.24, 23.04, 14.46; MS (MALDI) *m/z* calcd for C_53_H_104_N_2_O_5_ (M^+^) 848.403; found: 849.405 [M+H]^+^.

**Synthesis of O12-K6 series lipids**

Synthesis of O12-K6-H7, O12-K6-H8, O12-K6-H9 and O12-K6-H16

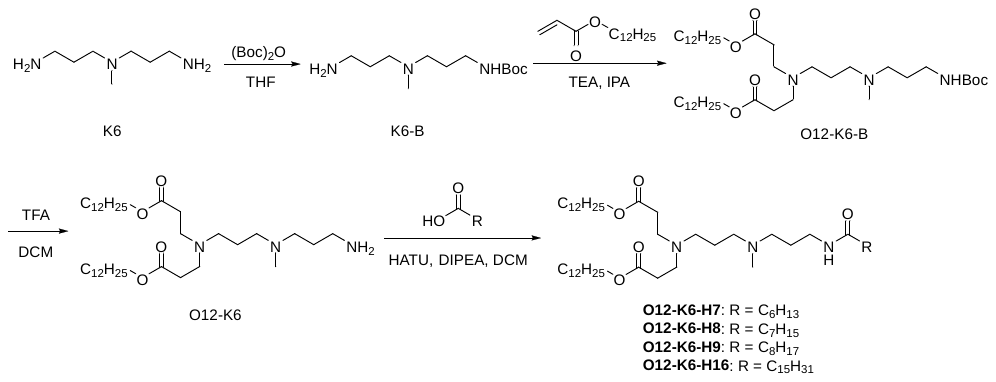

A 250 mL round-bottom flask equipped with a magnetic stir bar was charged with N^1^-(3-aminopropyl)-N^1^-methylpropane-1,3-diamine (K6, 1.5 g, 10.33 mmol) and 20 mL of THF, and the mixture was stirred to homogeneity. Separately, (Boc)_2_O (1.13 g, 5.18 mmol) was dissolved in 20 mL of THF. The (Boc)_2_O solution was then slowly added dropwise to the K6 solution *via* a constant-pressure dropping funnel under ice-bath cooling, and the reaction was maintained for 5 h while monitored by TLC. Upon reaction completion, the mixture was extracted three times with saturated sodium bicarbonate solution. The crude product was purified by column chromatography to afford K6-B as a white solid (795 mg, 3.24 mmol), with a yield of 31.4%.

The synthetic route of intermediate O12-K6-B was analogous to that of O12-D3-C12. Subsequently, 8 mL of trifluoroacetic acid (TFA) was slowly added to a solution of O12-K6-B (1.45 g, 2 mmol) in DCM under ice-bath conditions, and the resulting mixture was stirred for 1 h. After full conversion was confirmed by TLC, the solvent was removed under reduced pressure. The residue was washed three times with saturated sodium bicarbonate solution, and the solvent was again evaporated under reduced pressure to yield O12-K6 as a pale-yellow transparent liquid (1.25 g, 2 mmol), with a yield of 100.0%. Following the synthetic procedure established for O12-D3-H8, the final products O12-K6-H7, O12-K6-H8, O12-K6-H9 and O12-K6-H16 were synthesized starting from O12-K6.

O12-K6-H7 (didodecyl 3,3’-((3-((3-heptanamidopropyl)(methyl)amino)propyl) azanediyl)dipropionate): Off white solid, 27.6% yield. ^1^H NMR (600 MHz, CDCl_3_) δ ppm 6.78 (t, *J =* 6.4 Hz, 1H), 4.02 (t, *J =* 6.8 Hz, 4H), 3.32 (q, *J =* 6.2 Hz, 2H), 3.15 (t, *J =* 7.3 Hz, 2H), 3.09 (t, *J =* 7.0 Hz, 2H), 2.82 (s, 3H), 2.74 (t, *J =* 6.6 Hz, 4H), 2.59 (t, *J =* 6.0 Hz, 2H), 2.43 (t, *J =* 6.6 Hz, 4H), 2.23 (t, *J =* 7.7 Hz, 2H), 1.98 (p, *J =* 6.7 Hz, 2H), 1.90 (p, *J =* 6.7 Hz, 2H), 1.59 (dq, *J =* 12.0, 6.1, 5.4 Hz, 6H), 1.32-1.24 (m, 42H), 0.85 (q, *J =* 4.0 Hz, 9H); ^13^C NMR (150MHz, CDCl_3_) δ ppm 172.67, 172.62, 64.89, 55.19, 53.55, 51.24, 48.25, 39.94, 35.89, 35.39, 31.79, 31.36, 31.30, 29.54, 29.52, 29.49, 29.43, 29.23, 29.20, 28.78, 28.46, 25.81, 25.48, 24.23, 22.55, 22.37, 21.21, 13.97, 13.89; MS (MALDI) *m/z* calcd for C_44_H_87_N_3_O_5_ (M^+^) 737.179; found: 738.438 [M+H]^+^.

O12-K6-H8 (didodecyl 3,3’-((3-(methyl(3-octanamidopropyl)amino)propyl) azanediyl)dipropionate): Off white solid, 22.9% yield. ^1^H NMR (600 MHz, CDCl_3_) δ ppm 6.90 (t, *J =* 6.2 Hz, 1H), 4.03 (t, *J =* 6.8 Hz, 4H), 3.36 (q, *J =* 6.1 Hz, 2H), 3.08 (dt, *J =* 17.1, 7.2 Hz, 4H), 2.78 (s, 3H), 2.72 (t, *J =* 6.6 Hz, 4H), 2.56 (t, *J =* 5.9 Hz, 2H), 2.42 (t, *J =* 6.6 Hz, 4H), 2.26 – 2.23 (m, 2H), 2.03 (q, *J =* 6.1 Hz, 2H), 1.92 (dt, *J =* 12.3, 6.8 Hz, 2H), 1.60 (q, *J =* 6.5 Hz, 6H), 1.34 – 1.25 (m, *J =* 14.3 Hz, 44H), 0.87 (t, *J =* 4.5 Hz, 9H); ^13^C NMR (150 MHz, CDCl_3_) δ ppm 172.62, 64.88, 55.15, 53.76, 51.31, 48.45, 40.07, 36.21, 35.73, 31.85, 31.77, 31.64, 29.62, 29.57, 29.49, 29.29, 29.25, 29.19, 28.94, 28.54, 25.88, 25.64, 24.85, 24.19, 22.62, 22.54, 14.04, 13.99; MS (MALDI) *m/z* calcd for C_45_H_89_N_3_O_5_ (M^+^) 751.205; found: 752.453 [M+H]^+^.

O12-K6-H9 (didodecyl 3,3’-((3-(methyl(3-nonanamidopropyl)amino)propyl) azanediyl)dipropionate): Off white solid, 20.4% yield. ^1^H NMR (600 MHz, CDCl_3_) δ ppm 6.75 (t, *J =* 6.5 Hz, 1H), 4.03 (t, *J =* 6.8 Hz, 4H), 3.34 (q, *J =* 6.2 Hz, 2H), 3.14 (dt, *J =* 37.4, 7.0 Hz, 4H), 2.84 (s, 3H), 2.76 (t, *J =* 6.5 Hz, 4H), 2.61 (t, *J =* 6.1 Hz, 2H), 2.45 (t, *J =* 6.6 Hz, 4H), 2.25 (t, *J =* 7.7 Hz, 2H), 2.00 (p, *J =* 6.6 Hz, 2H), 1.92 (p, *J =* 6.7 Hz, 2H), 1.61 (p, *J =* 6.5 Hz, 6H), 1.34 – 1.25 (m, 46H), 0.87 (d, *J =* 7.2 Hz, 9H); ^13^C NMR (150 MHz, CDCl_3_) δ ppm 175.80, 171.66, 63.99, 54.23, 52.61, 50.32, 47.30, 39.06, 34.97, 34.44, 30.84, 30.75, 30.72, 30.28, 28.59, 28.57, 28.54, 28.48, 28.25, 28.22, 28.14, 28.10, 28.03, 27.50, 24.86, 24.58, 23.71, 23.24, 21.60, 21.56, 13.02, 13.00; MS (MALDI) *m/z* calcd for C_46_H_91_N_3_O_5_ (M^+^) 765.232; found: 766.450 [M+H]^+^.

O12-K6-H16 (didodecyl 3,3’-((3-(methyl(3-palmitamidopropyl)amino)propyl) azanediyl)dipropionate): Off white solid, 24.1% yield. ^1^H NMR (600 MHz, CDCl_3_) δ ppm 6.69 (t, *J =* 6.4 Hz, 1H), 4.04 (t, *J =* 6.8 Hz, 4H), 3.36 (q, *J =* 6.2 Hz, 2H), 3.15 (dt, *J =* 38.0, 7.0 Hz, 4H), 2.84 (s, 3H), 2.77 (s, 2H), 2.64 (s, 2H), 2.46 (t, *J =* 6.6 Hz, 4H), 2.34 (t, *J =* 7.5 Hz, 2H), 2.26 (t, *J =* 7.7 Hz, 2H), 2.02 (p, *J =* 6.5 Hz, 2H), 1.94 (p, *J =* 6.5 Hz, 2H), 1.62 (p, *J =* 6.9 Hz, 6H), 1.29 – 1.25 (m, 60H), 0.87 (d, *J =* 7.2 Hz, 9H); ^13^C NMR (150 MHz, CDCl_3_) δ ppm 177.24, 173.07, 65.51, 54.18, 51.82, 48.80, 40.63, 36.47, 35.90, 34.11, 32.28, 30.08, 30.06, 30.03, 30.01, 29.98, 29.95, 29.92, 29.80, 29.74, 29.72, 29.68, 29.66, 29.60, 29.43, 28.94, 26.29, 26.04, 25.08, 24.67, 23.05, 14.47; MS (MALDI) *m/z* calcd for C_53_H_105_N_3_O_5_ (M^+^) 863.809; found: 864.718 [M+H]^+^, 886.695 [M+Na]^+^.

**Synthesis of O12-P6 series lipids**

Synthesis of O12-P6-H7, O12-P6-H8 and O12-P6-H9

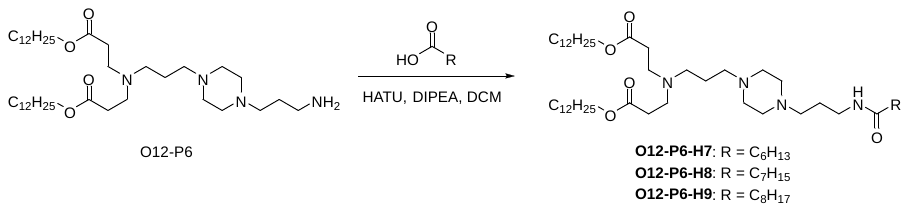

Lipids belonging to the O12-P6 series were synthesized following the same protocol as described for O12-K6-H7.

O12-P6-H7 (didodecyl 3,3’-((3-(4-(3-heptanamidopropyl)piperazin-1-yl)propyl) azanediyl)dipropionate): Off white solid, 23.5% yield. ^1^H NMR (600 MHz, CDCl_3_) δ ppm 6.95 (d, *J =* 24.4 Hz, 1H), 4.03 (t, *J =* 6.8 Hz, 4H), 3.32 (q, *J =* 5.8 Hz, 2H), 2.74 (t, *J =* 7.1 Hz, 4H), 2.70 – 2.57 (m, 4H), 2.60 – 2.51 (m, 4H), 2.42 (dt, *J =* 20.1, 7.0 Hz, 10H), 2.14 (t, *J =* 7.7 Hz, 2H), 1.72 (p, *J =* 6.4 Hz, 2H), 1.61 (tt, *J =* 13.9, 7.0 Hz, 8H), 1.31 – 1.24 (m, 42H), 0.86 (t, *J =* 6.9 Hz, 9H); ^13^C NMR (150 MHz, CDCl_3_) δ ppm 173.40, 172.78, 64.74, 56.89, 56.03, 52.67, 51.51, 49.33, 38.84, 37.11, 32.75, 32.02, 31.70, 29.80, 29.76, 29.74, 29.71, 29.65, 29.45, 29.40, 29.15, 28.75, 26.05, 25.98, 25.03, 22.79, 22.64, 14.22, 14.16; MS (ESI) *m/z* calcd for C_47_H_92_N_4_O_5_ (M^+^) 792.257; found: 793.750 [M+H]^+^.

O12-P6-H8 (didodecyl 3,3’-((3-(4-(3-octanamidopropyl)piperazin-1-yl)propyl) azanediyl) dipropionate): Off white solid, 38.8% yield. ^1^H NMR (600 MHz, CDCl_3_) δ ppm 6.89 (s, 1H), 4.03 (t, *J =* 6.8 Hz, 4H), 3.32 (q, *J =* 5.9 Hz, 2H), 2.73 (t, *J =* 7.1 Hz, 4H), 2.72 – 2.61 (m, 4H), 2.56 (t, *J =* 6.7 Hz, 4H), 2.42 (dt, *J =* 20.9, 7.1 Hz, 10H), 2.14 (t, *J =* 7.6 Hz, 2H), 1.73 (t, *J =* 6.6 Hz, 2H), 1.60 (p, *J =* 7.0 Hz, 8H), 1.36 – 1.24 (m, 44H), 0.85 (t, *J =* 6.6 Hz, 9H); ^13^C NMR (150 MHz, CDCl_3_) δ ppm 173.67, 172.99, 64.96, 58.65, 56.96, 56.21, 51.71, 49.53, 38.93, 37.31, 32.95, 32.23, 32.04, 30.61, 30.01, 29.98, 29.95, 29.92, 29.87, 29.67, 29.66, 29.61, 29.38, 28.97, 26.27, 26.23, 25.26, 24.40, 23.00, 22.94, 14.43, 14.39; MS (ESI) *m/z* calcd for C_48_H_94_N_4_O_5_ (M^+^) 806.284; found: 807.800 [M+H]^+^.

O12-P6-H9 (didodecyl 3,3’-((3-(4-(3-nonanamidopropyl)piperazin-1-yl)propyl) azanediyl)dipropionate): Off white solid, 39.1% yield. ^1^H NMR (600 MHz, CDCl_3_) δ ppm 7.03 – 6.88 (m, 1H), 4.01 (t, *J =* 6.8 Hz, 4H), 3.31 (q, *J =* 5.9 Hz, 2H), 2.72 (t, *J =* 7.2 Hz, 4H), 2.71 – 2.59 (m, 4H), 2.53 (t, *J =* 6.5 Hz, 4H), 2.41 (dt, *J =* 20.9, 7.0 Hz, 10H), 2.12 (t, *J =* 7.6 Hz, 2H), 1.71 (q, *J =* 6.4 Hz, 2H), 1.59 (tt, *J =* 14.0, 7.1 Hz, 8H), 1.30 – 1.22 (m, 46H), 0.86 – 0.83 (m, 9H); ^13^C NMR (150 MHz, CDCl_3_) δ ppm 173.37, 172.74, 64.69, 56.85, 55.99, 52.64, 51.48, 49.31, 38.80, 37.07, 32.72, 31.98, 31.91, 29.76, 29.73, 29.70, 29.67, 29.62, 29.46, 29.43, 29.36, 29.26, 28.72, 26.02, 25.99, 25.02, 24.24, 22.75, 22.71, 14.18; MS (ESI) *m/z* calcd for C_49_H_96_N_4_O_5_ (M^+^) 820.310; found: 821.800 [M+H]^+^.

**^1^H NMR, ^13^C NMR and MS data for all synthesized final compounds**

C12-D3-A8

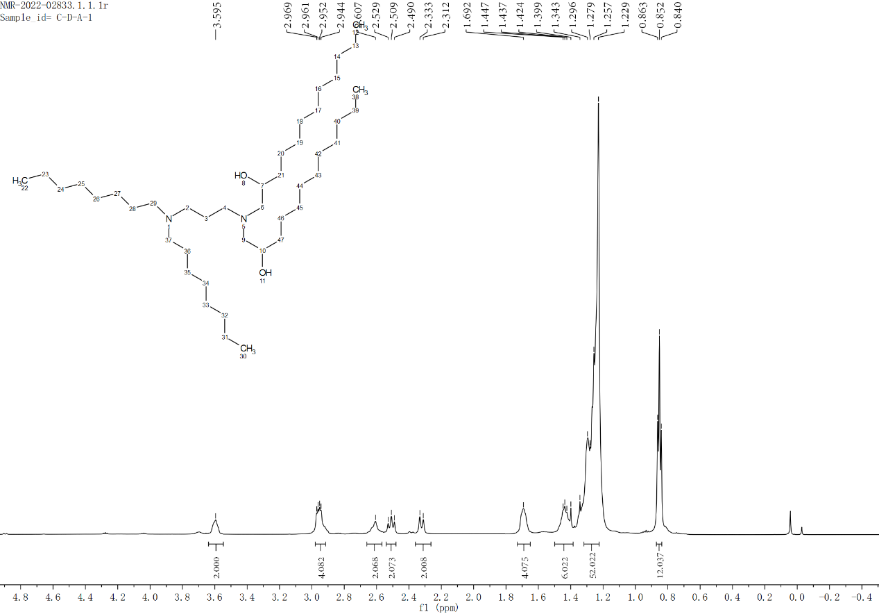

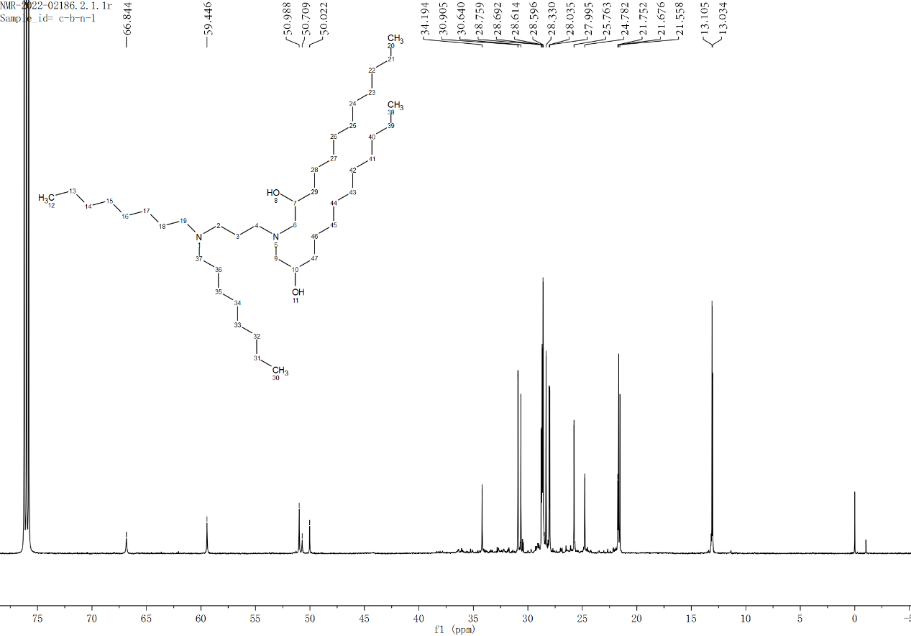

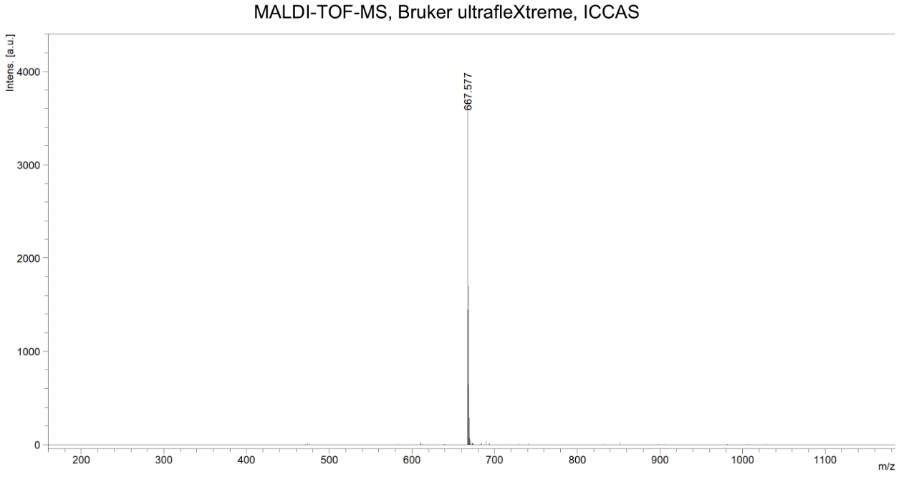

O12-D3-A8

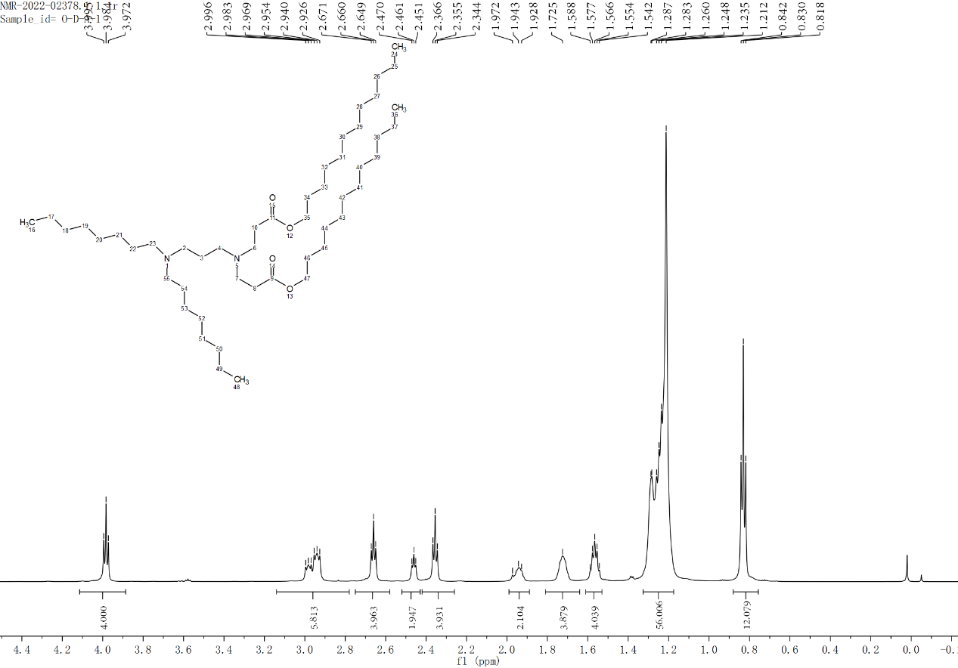

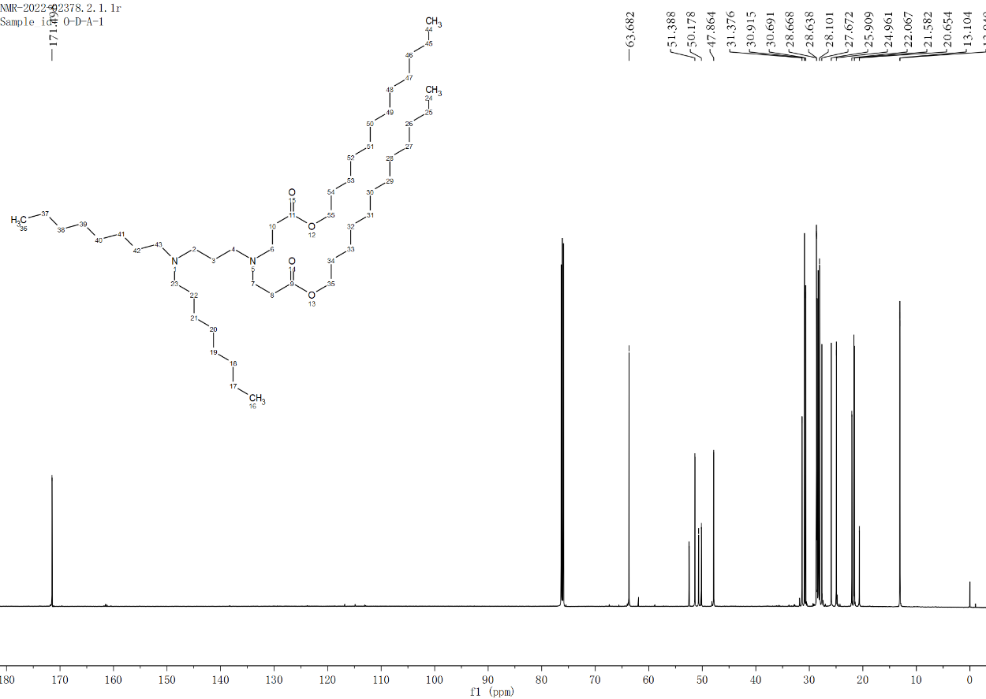

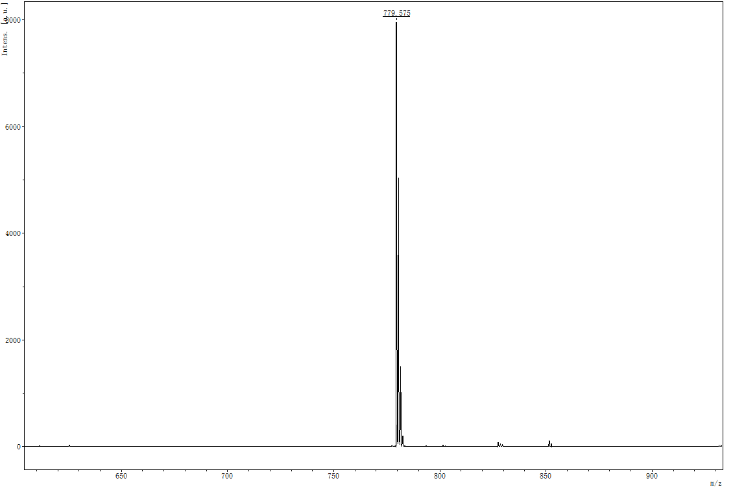

C12-D3-O12

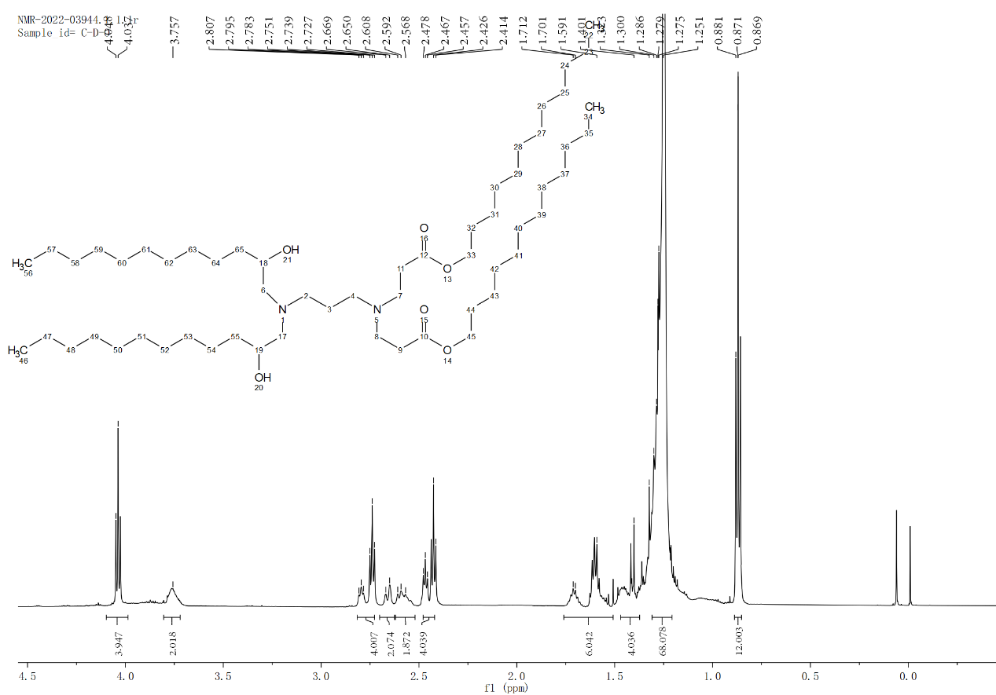

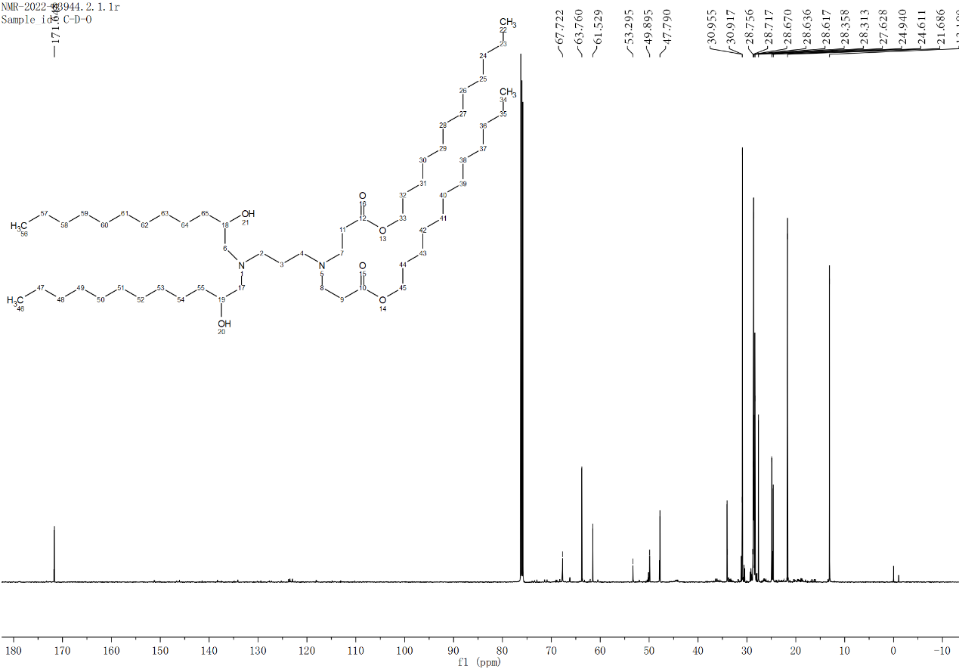

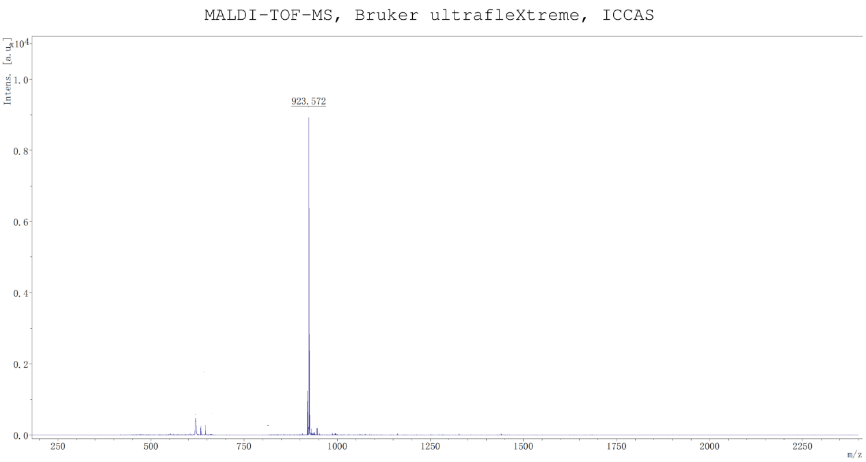

A8-E3-H8

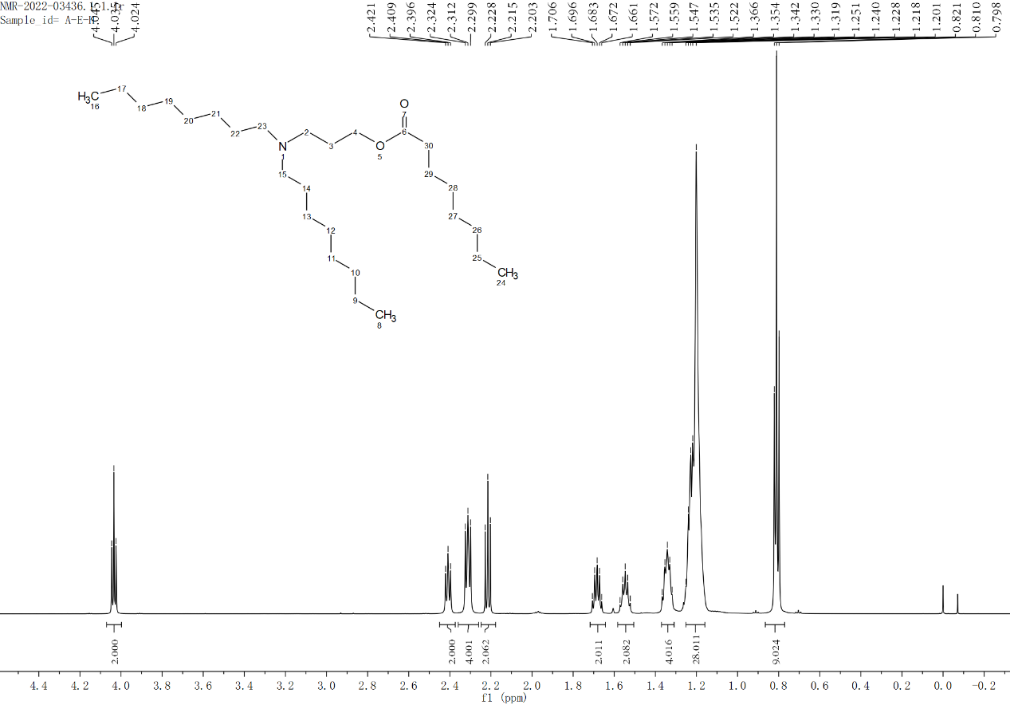

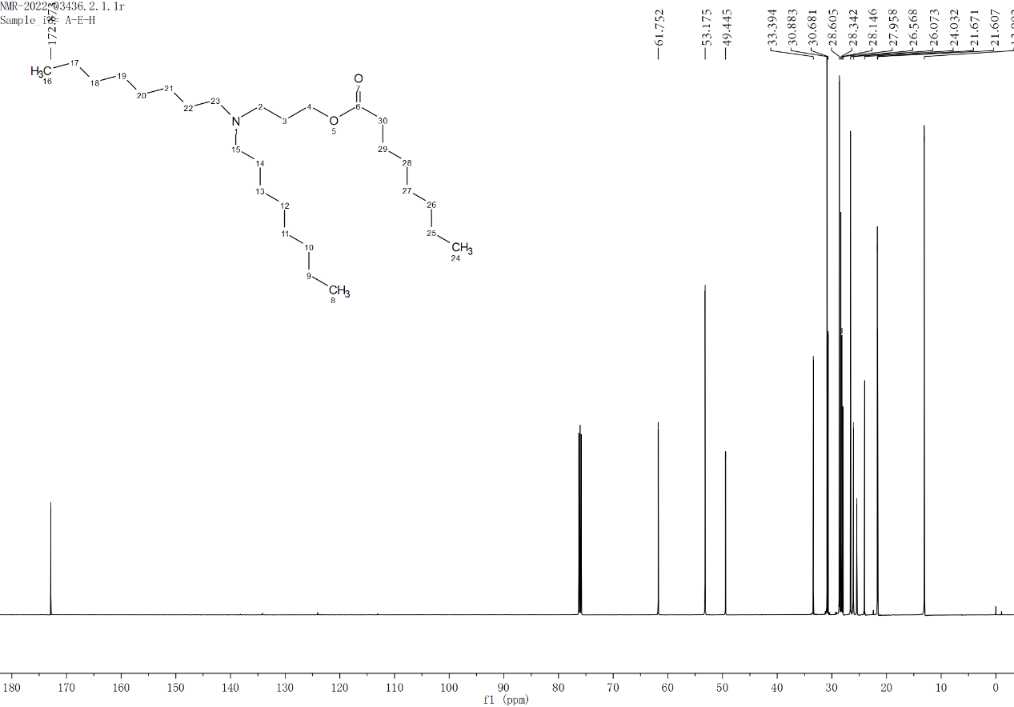

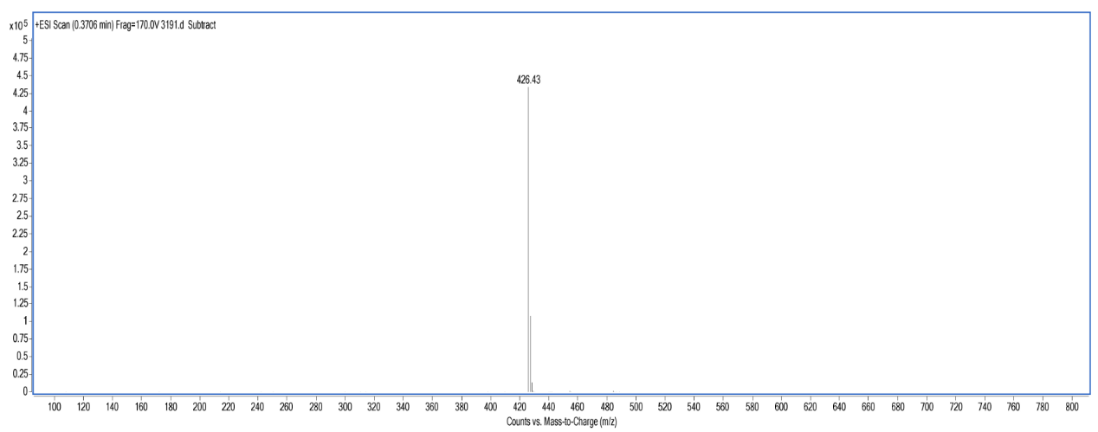

C12-E3-H8

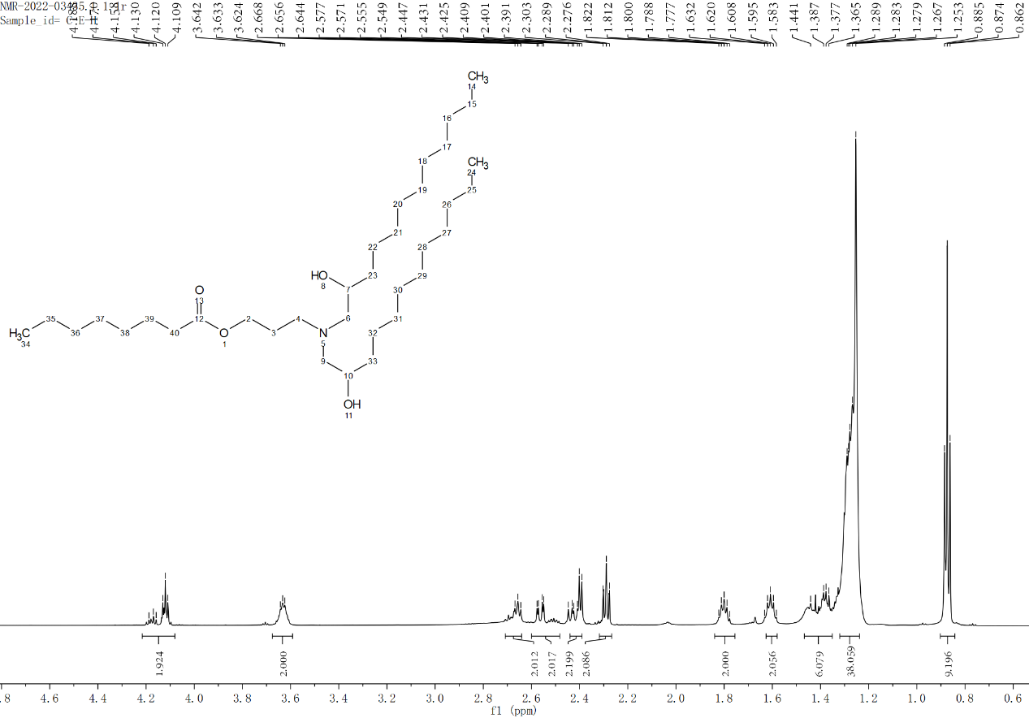

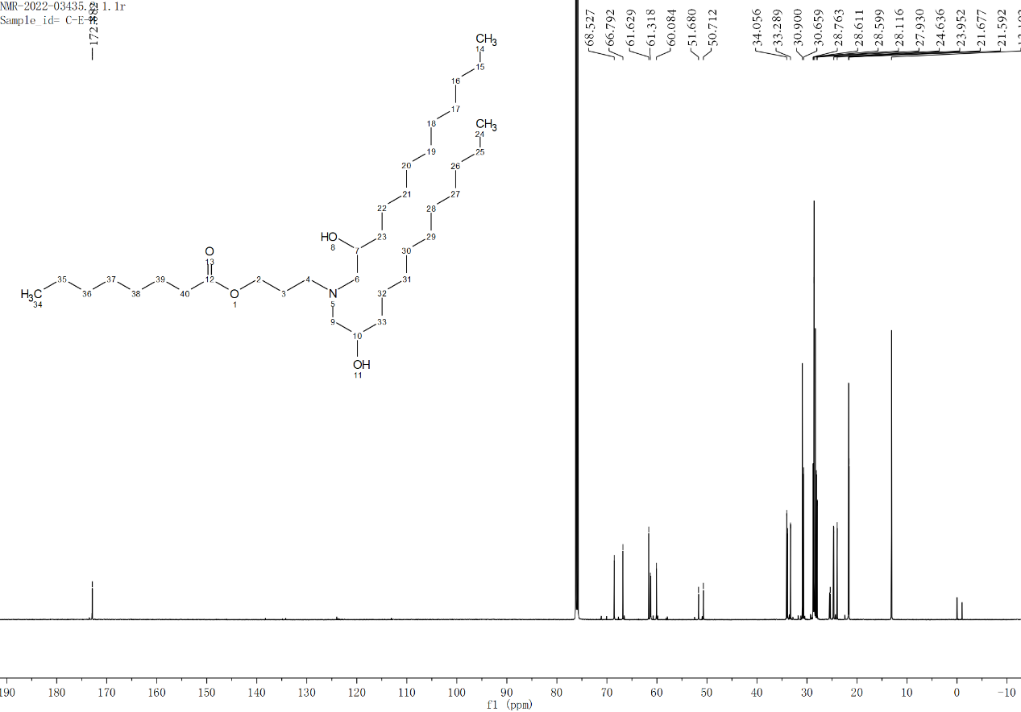

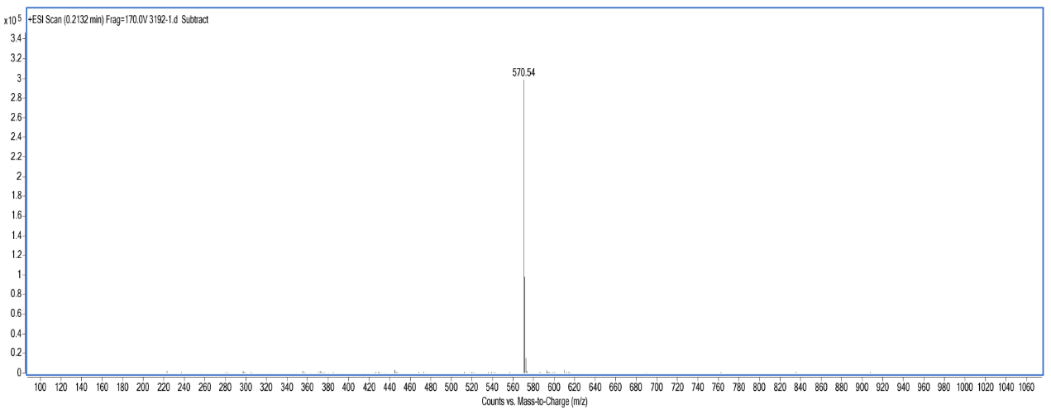

O12-E3-H8

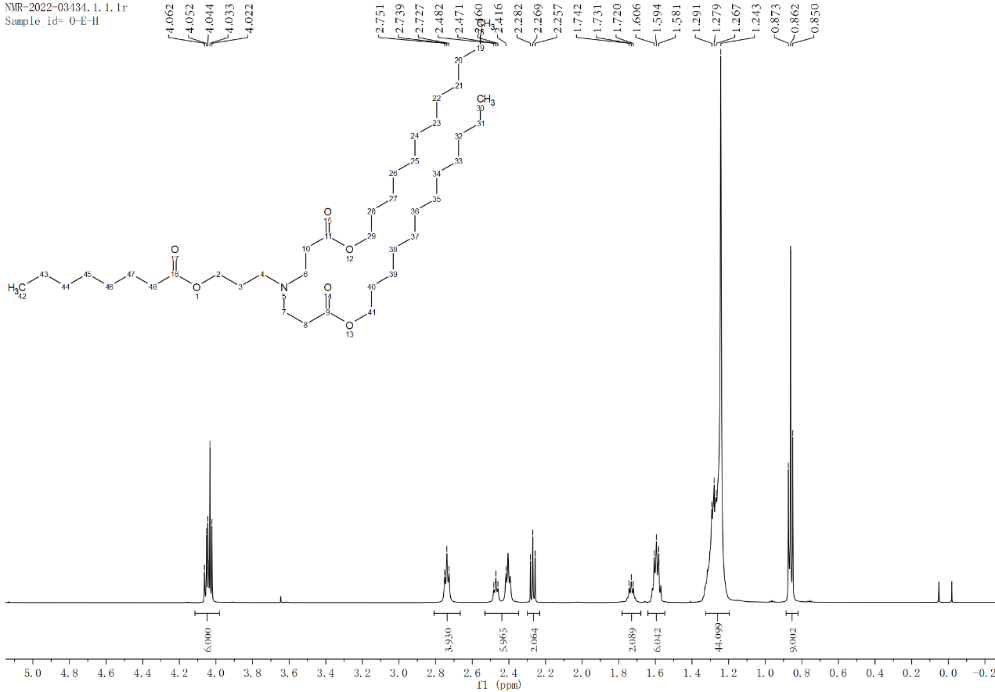

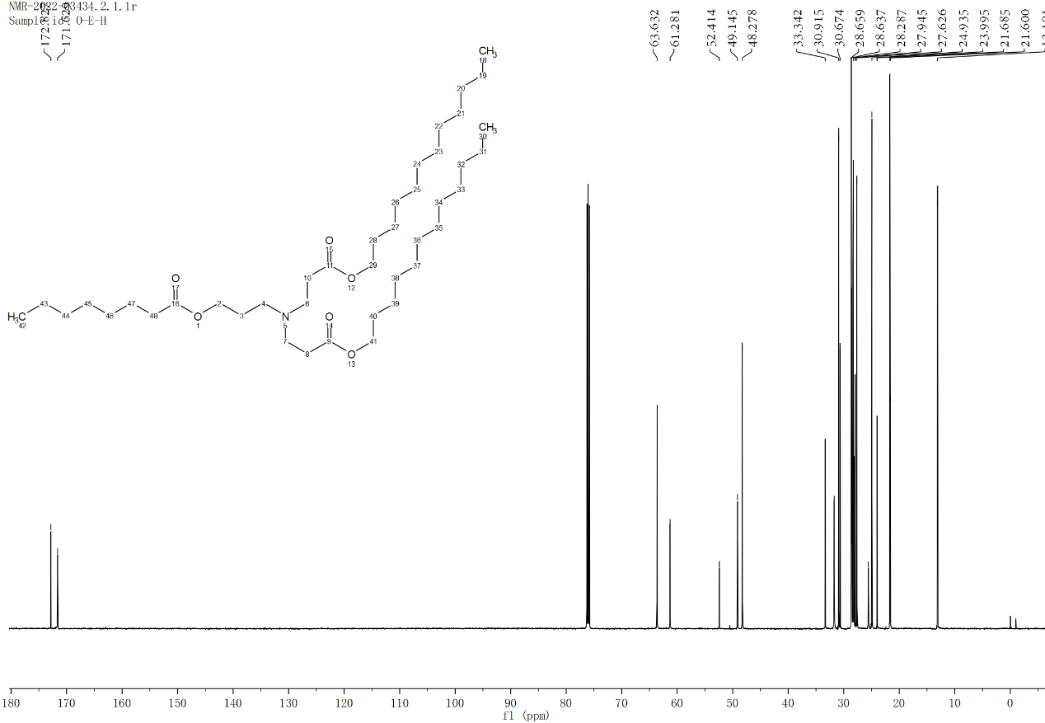

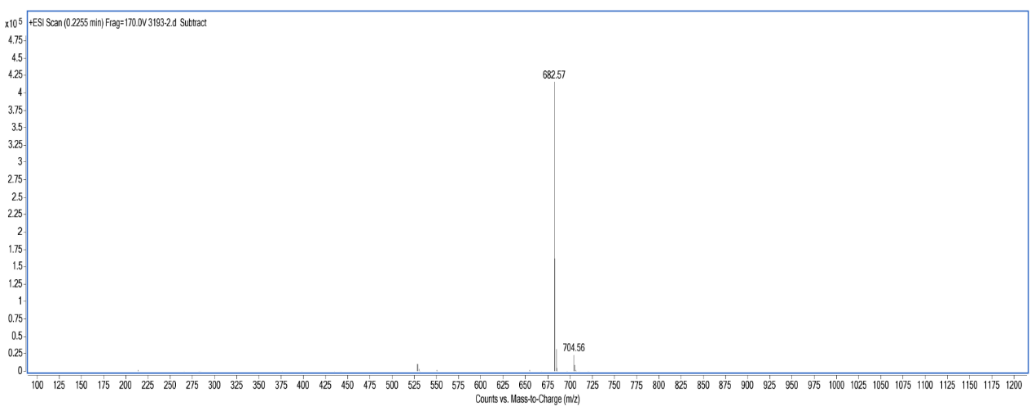

A8-D3-H8

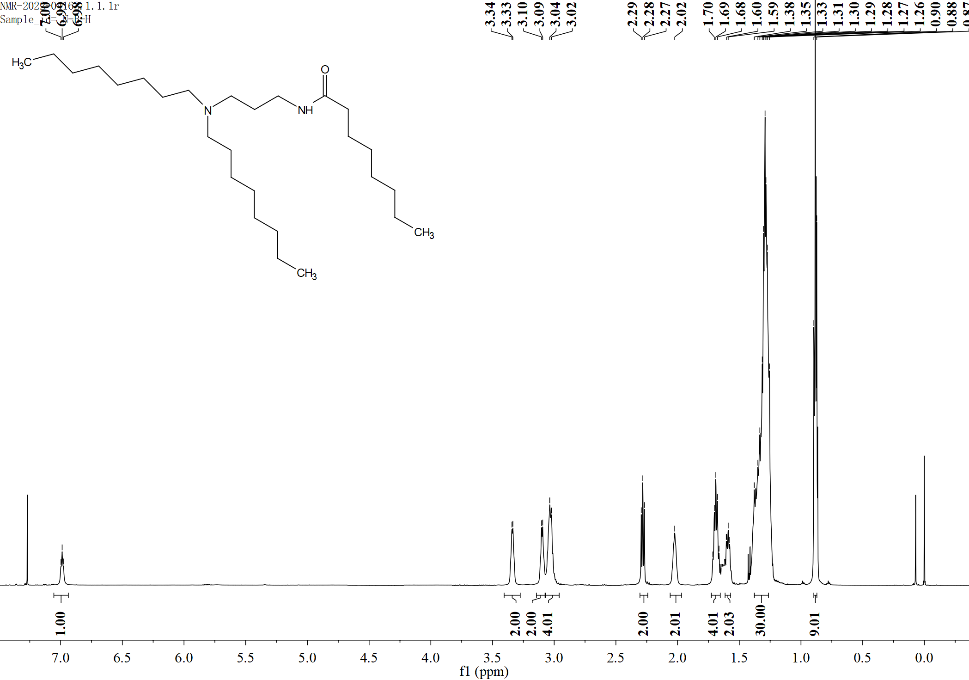

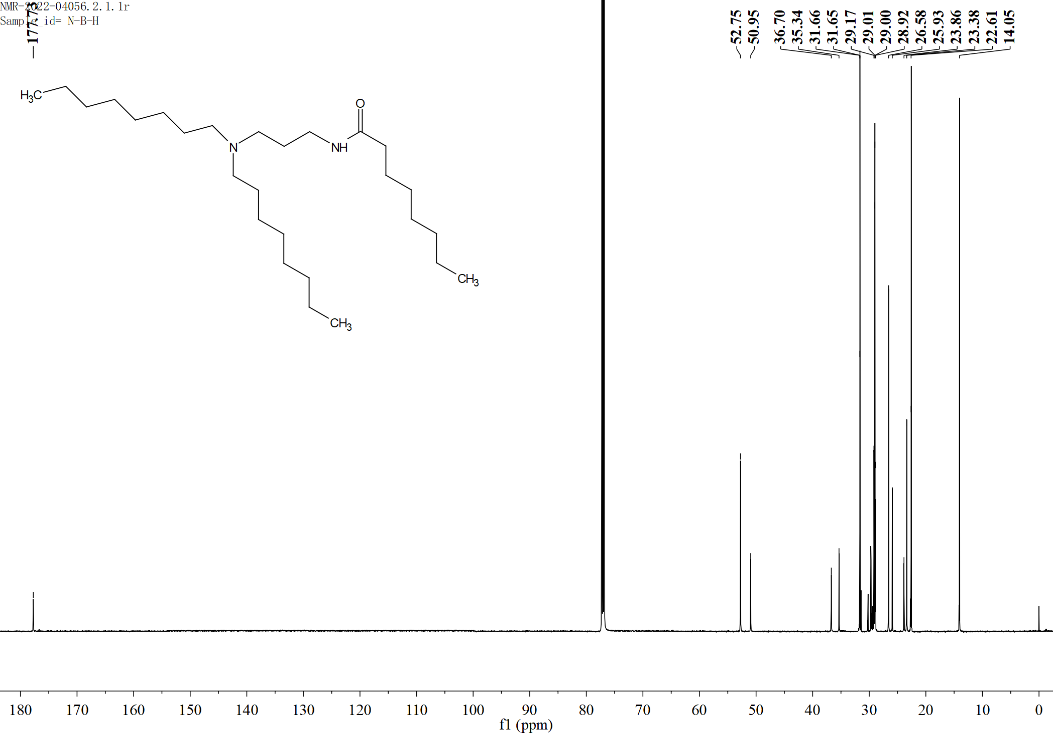

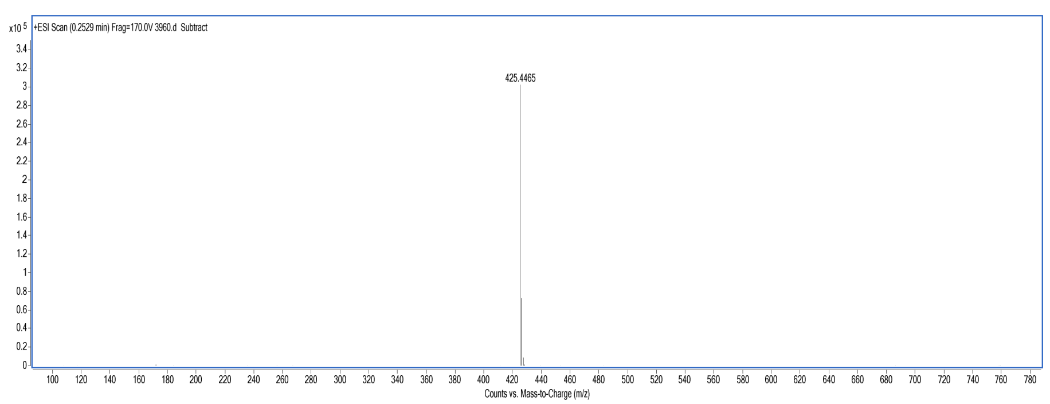

C12-D3-H8

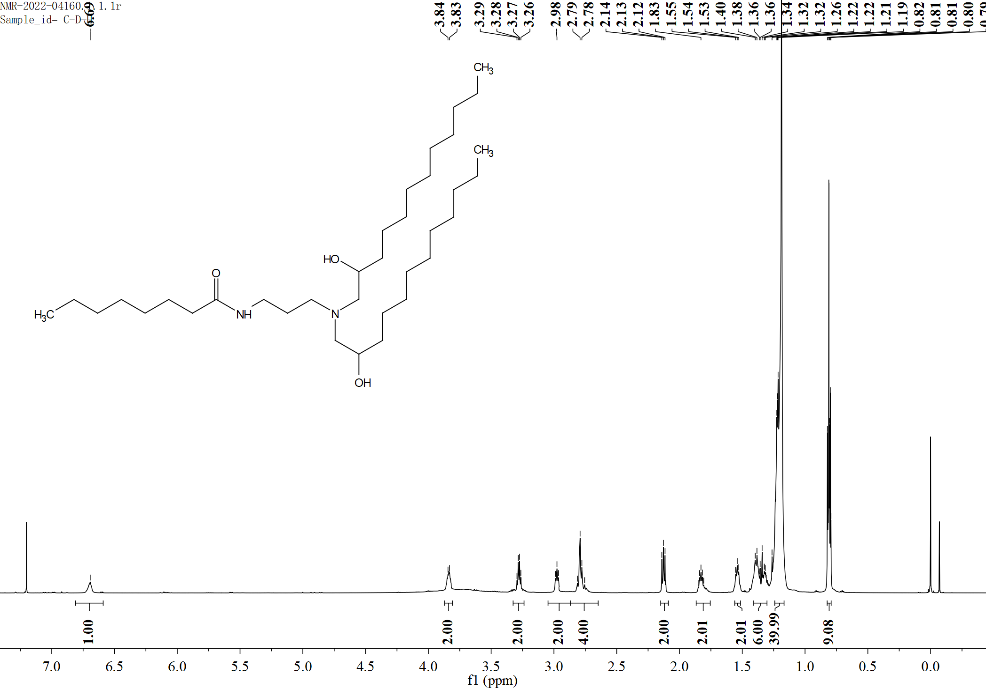

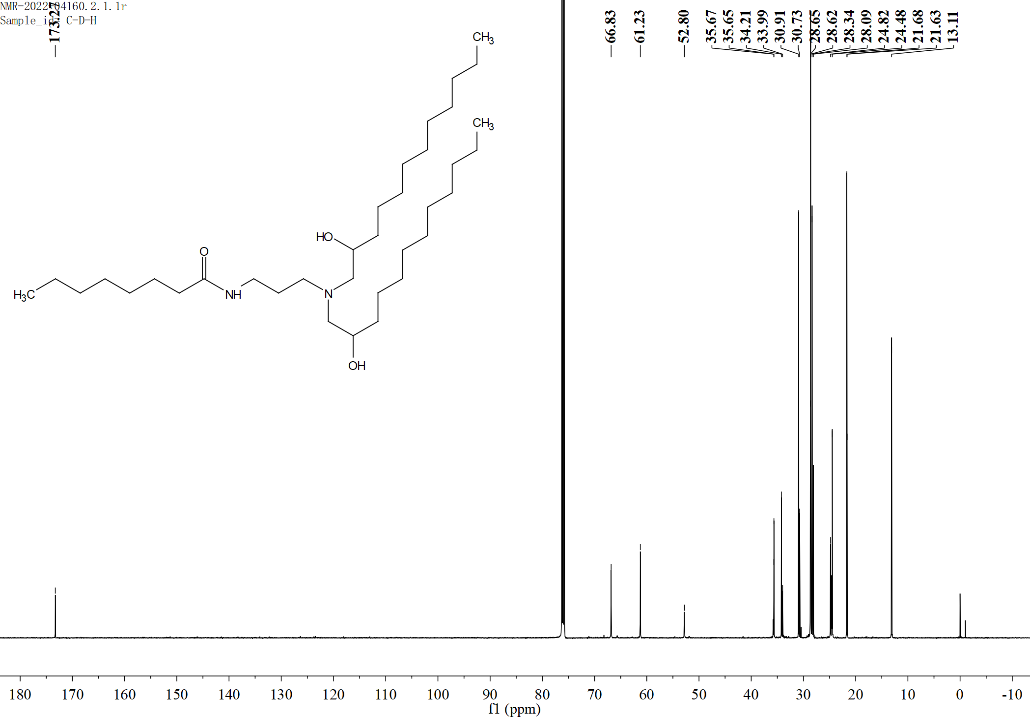

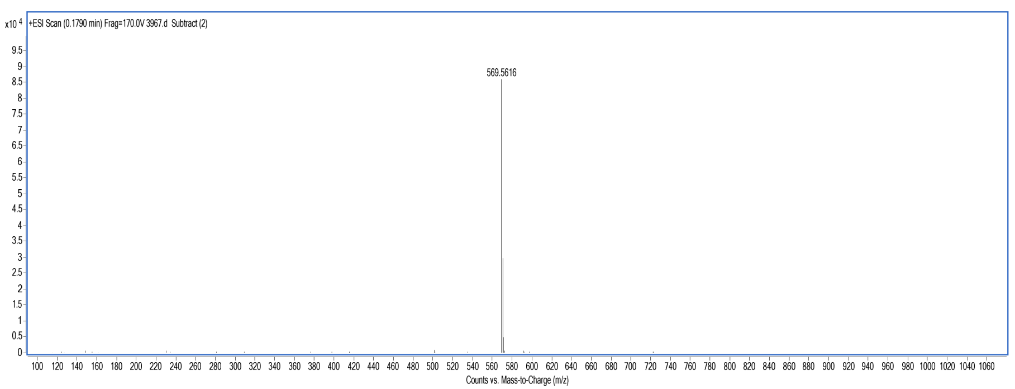

O12-D3-H8

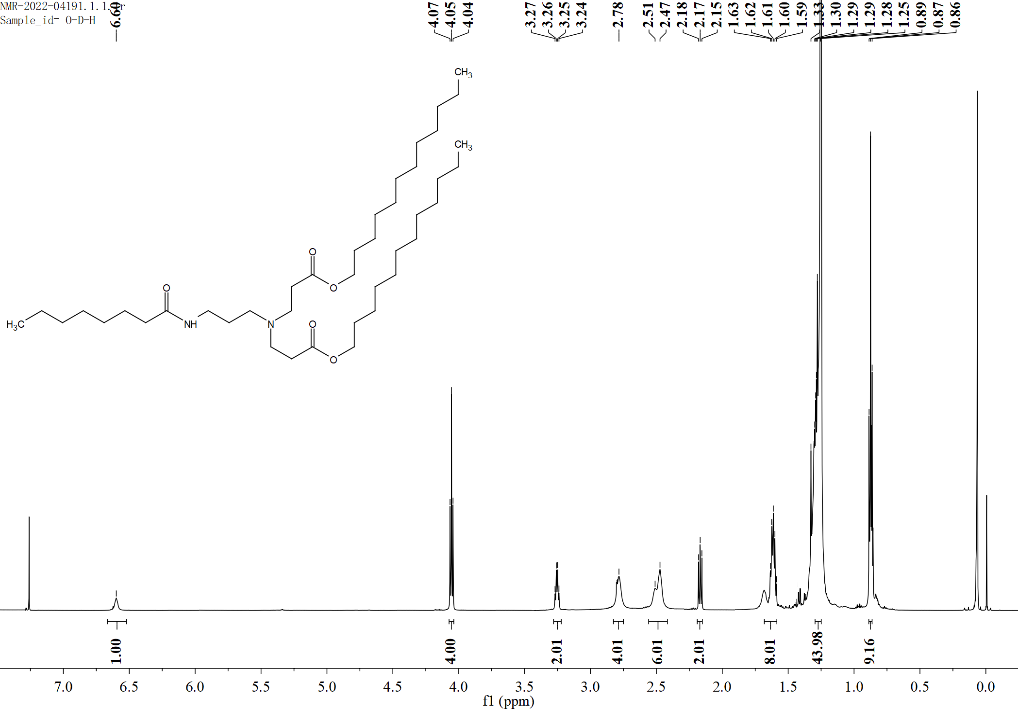

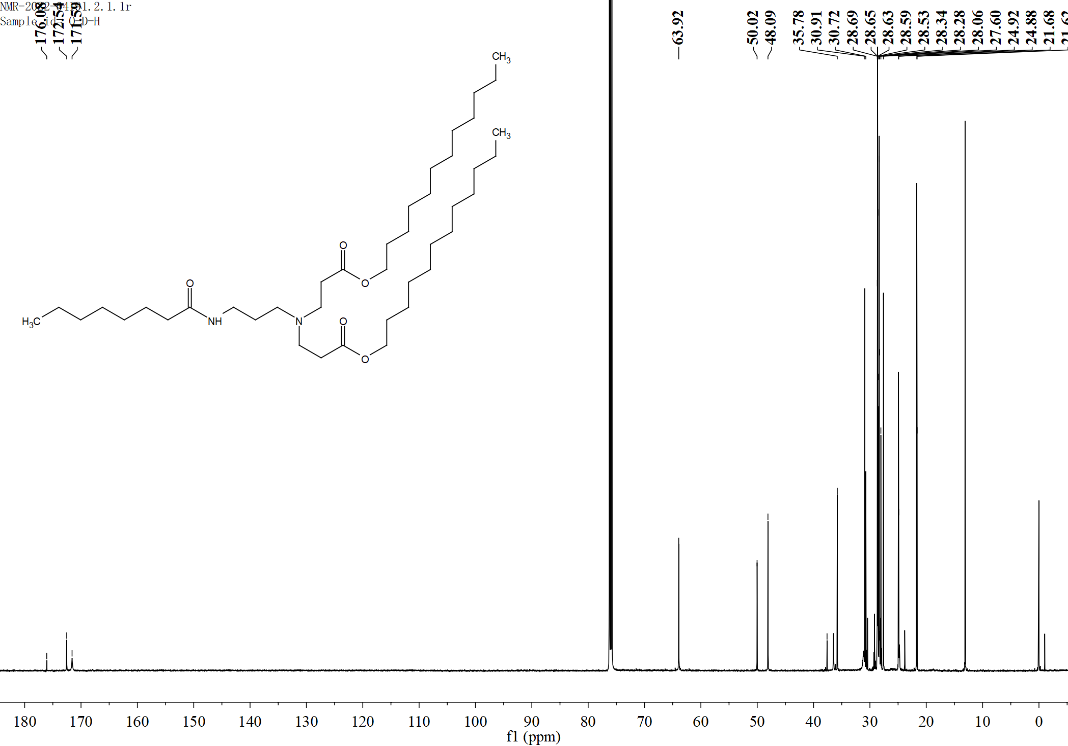

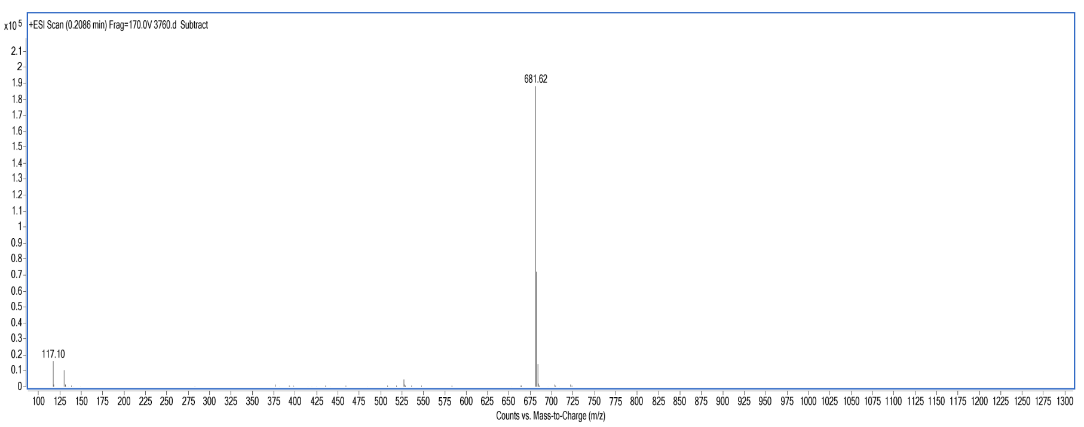

O18-D3-H7

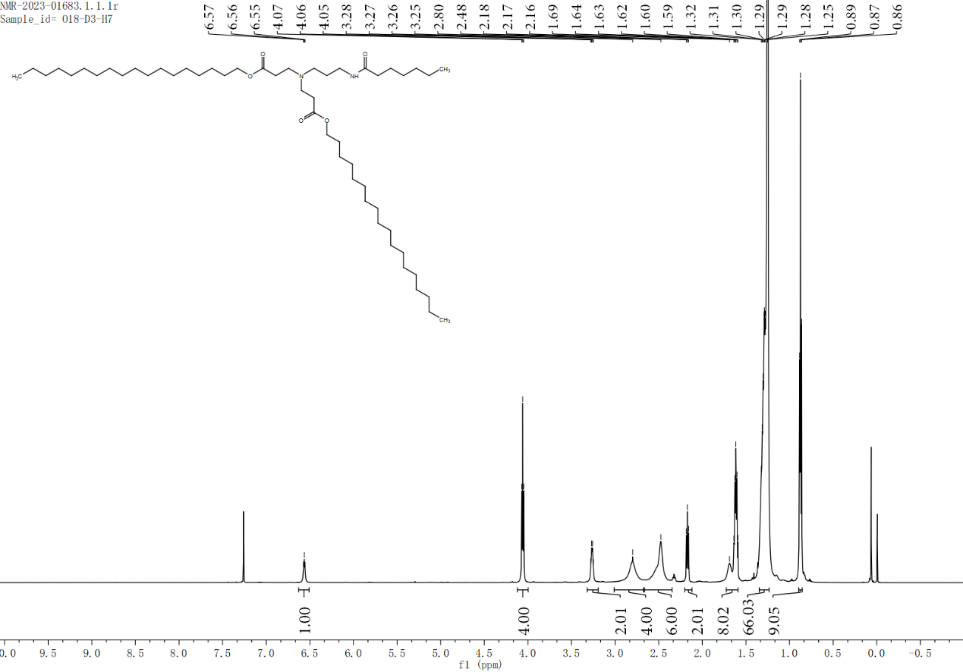

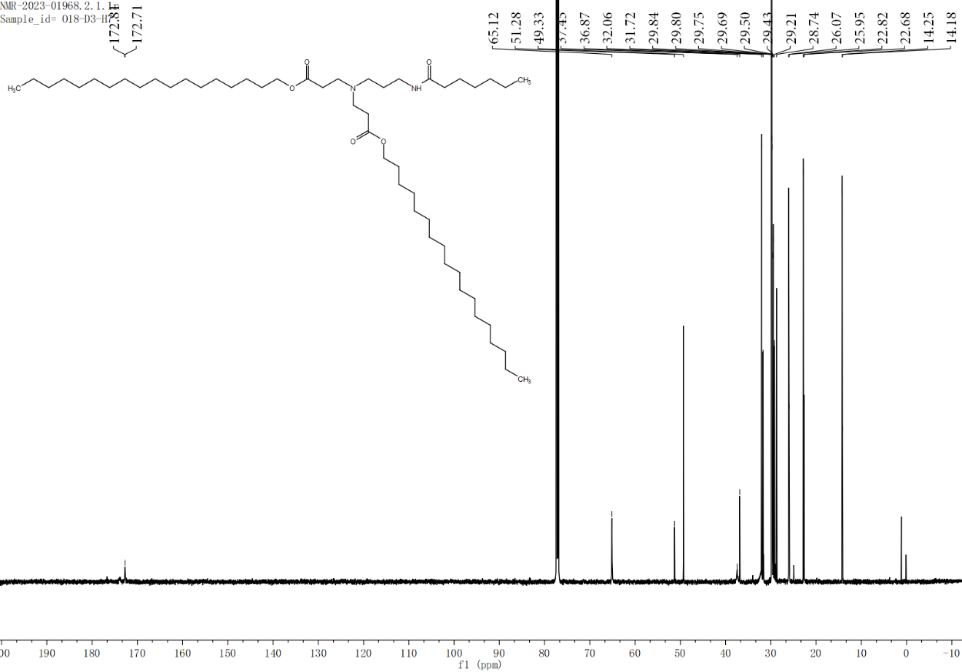

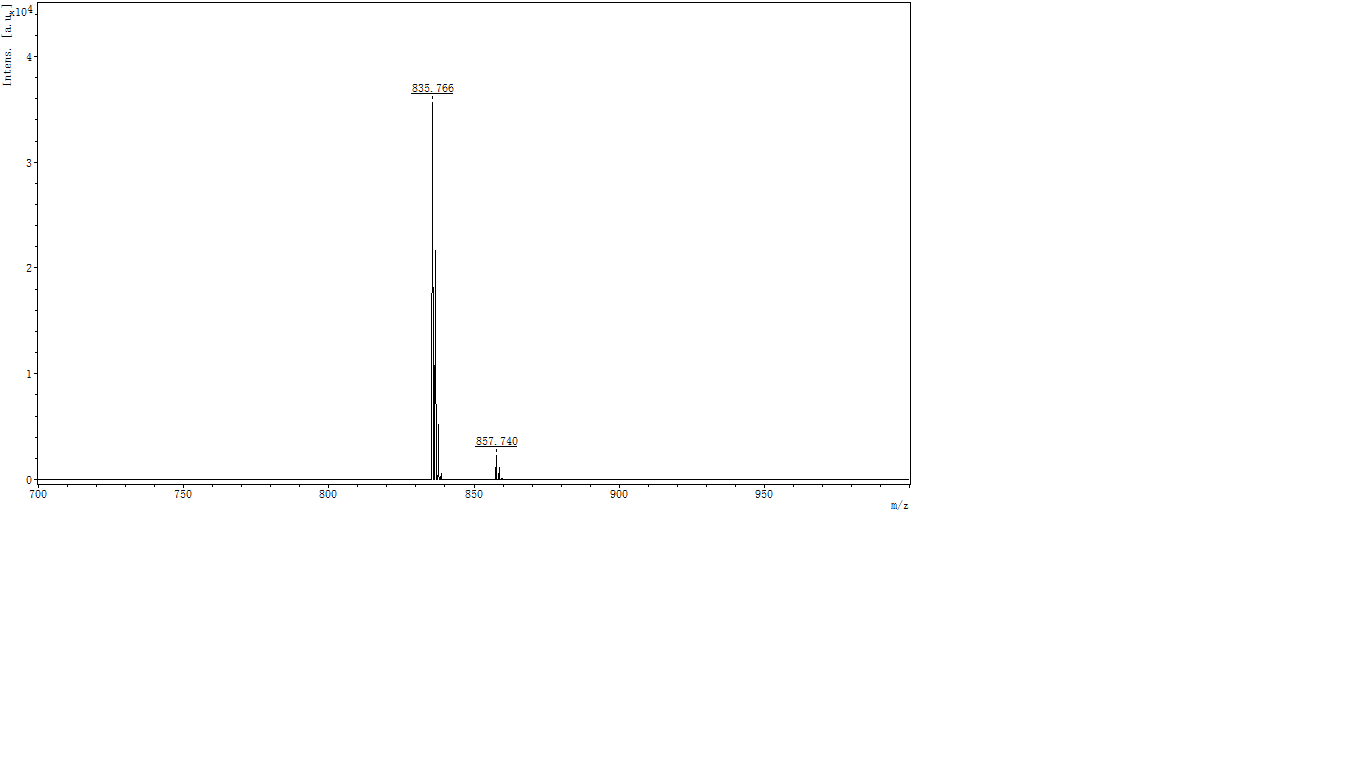

O18-D3-H8

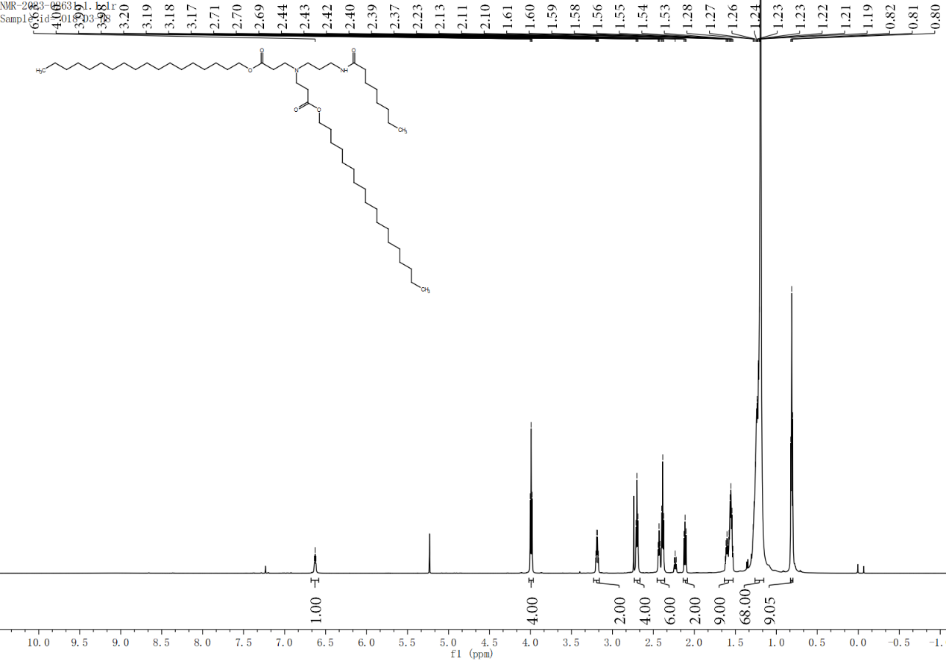

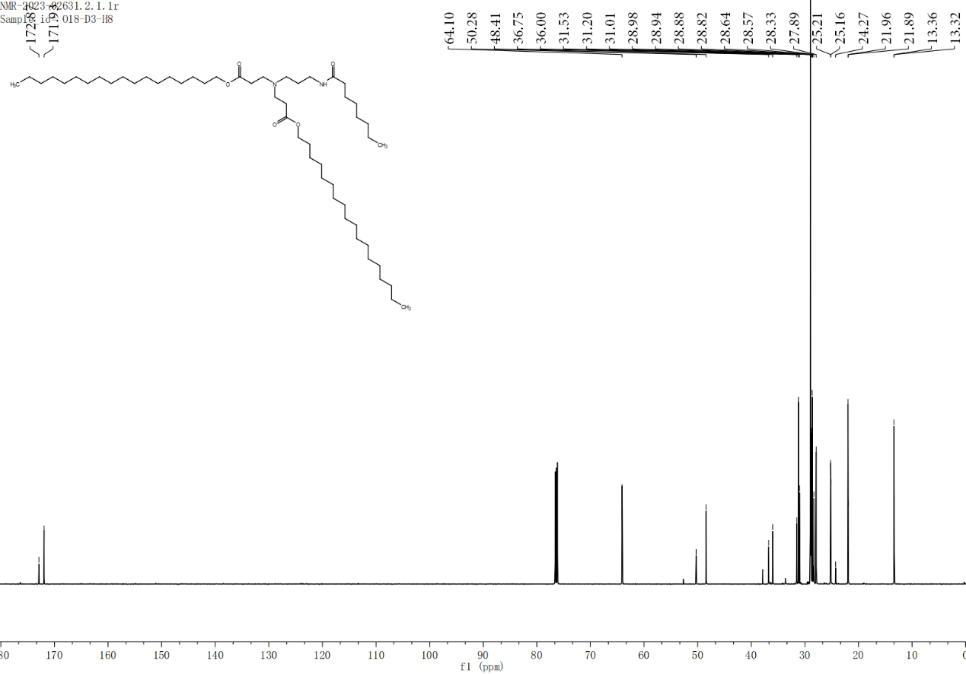

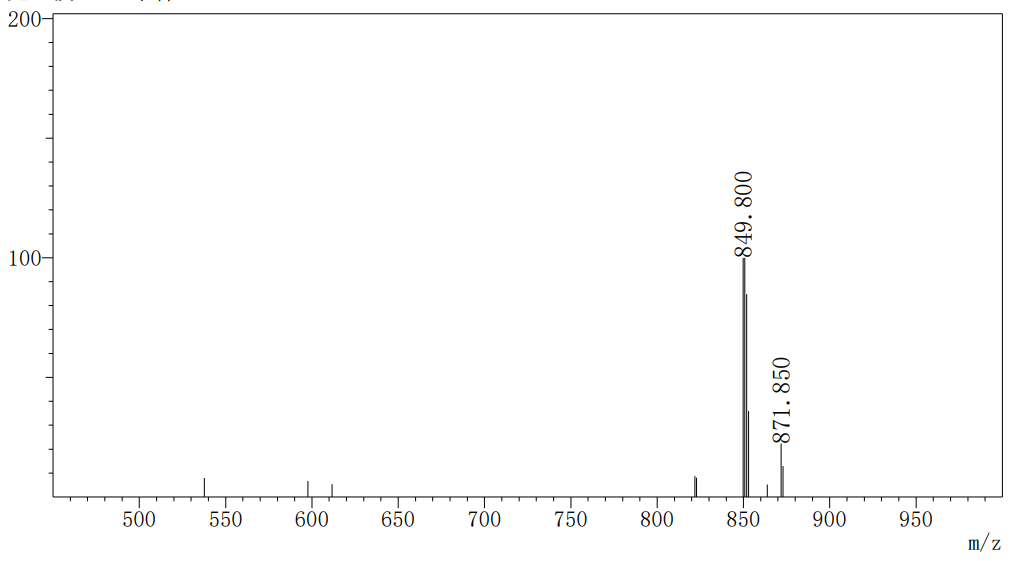

O12-D3-H9

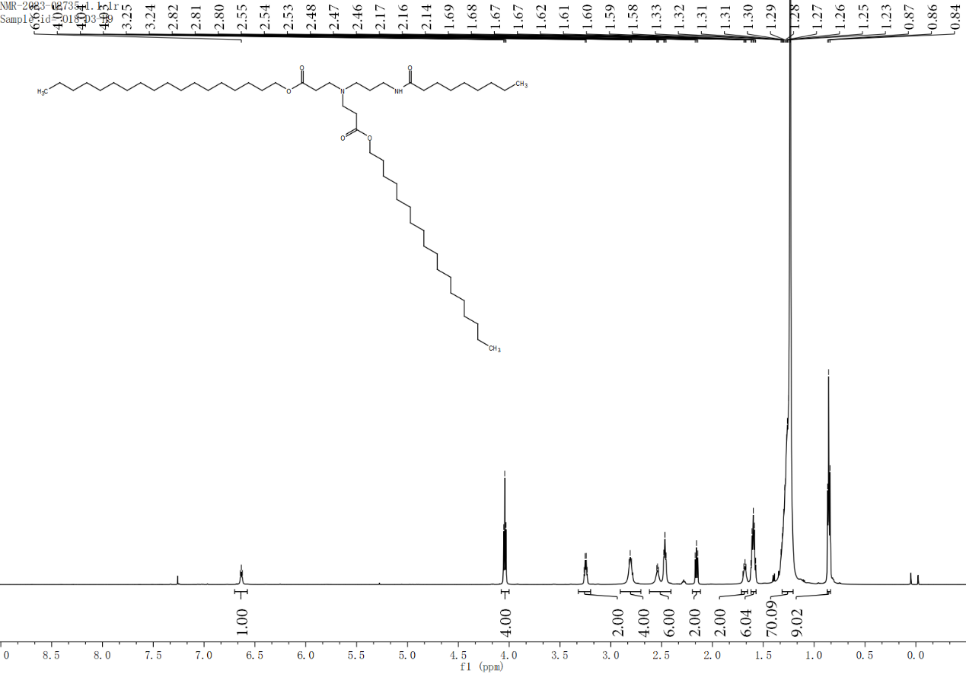

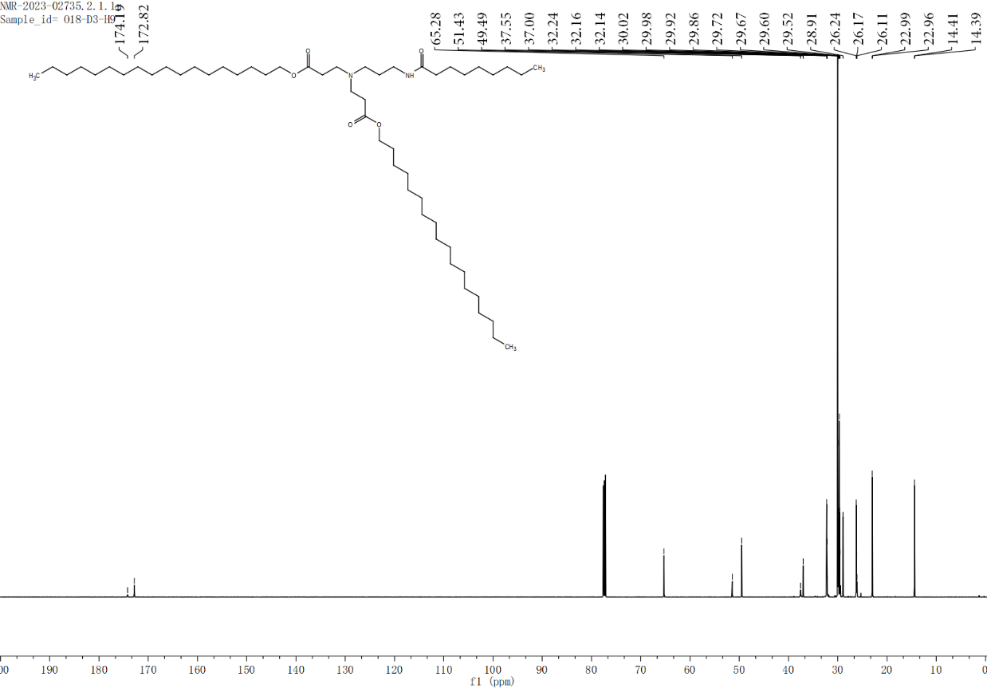

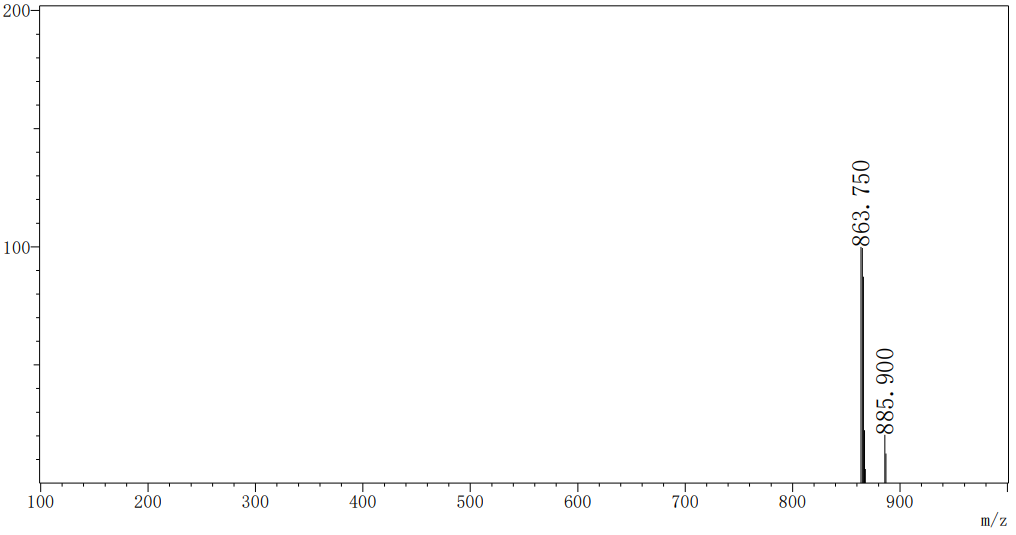

O12-D3-H18a

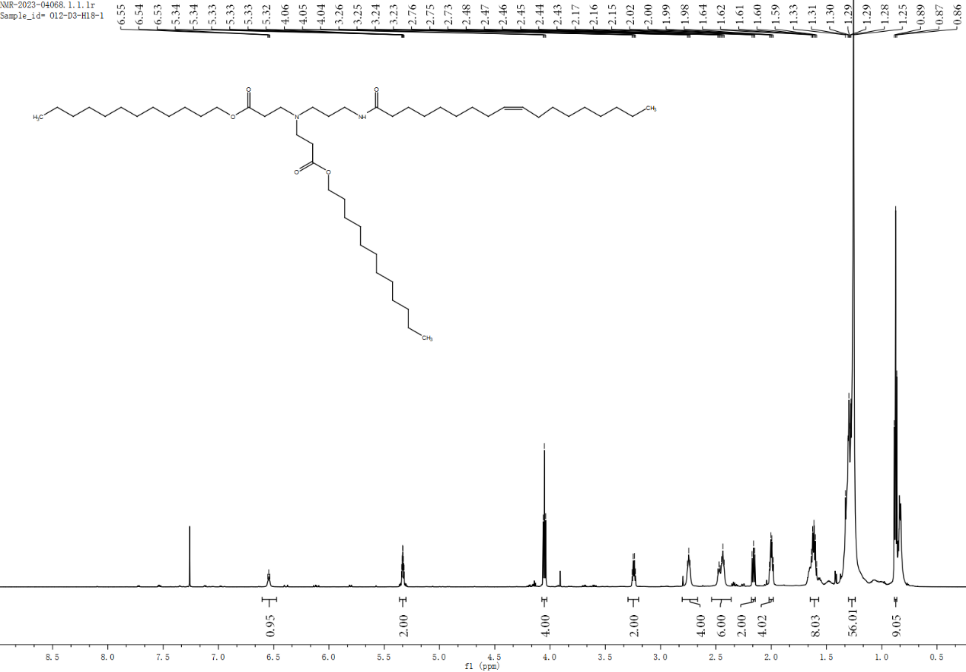

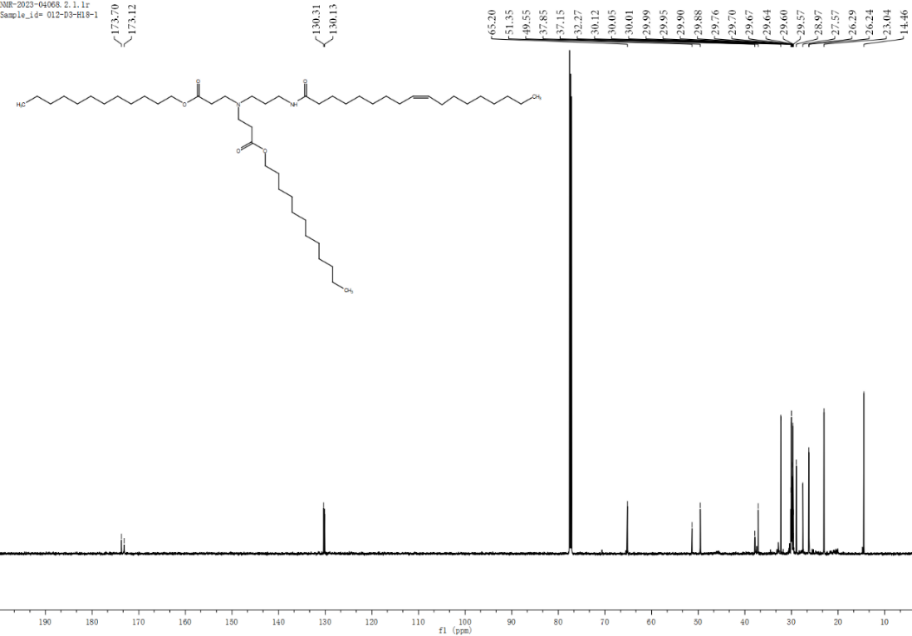

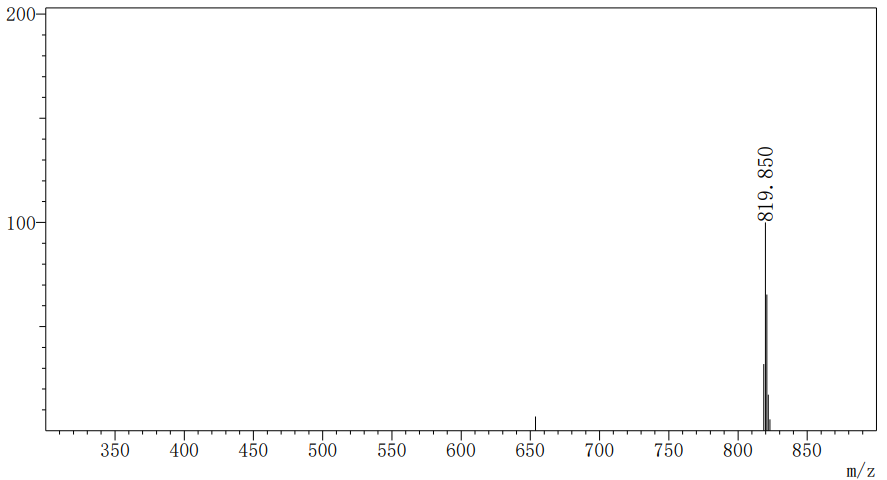

O12-D3-H18b

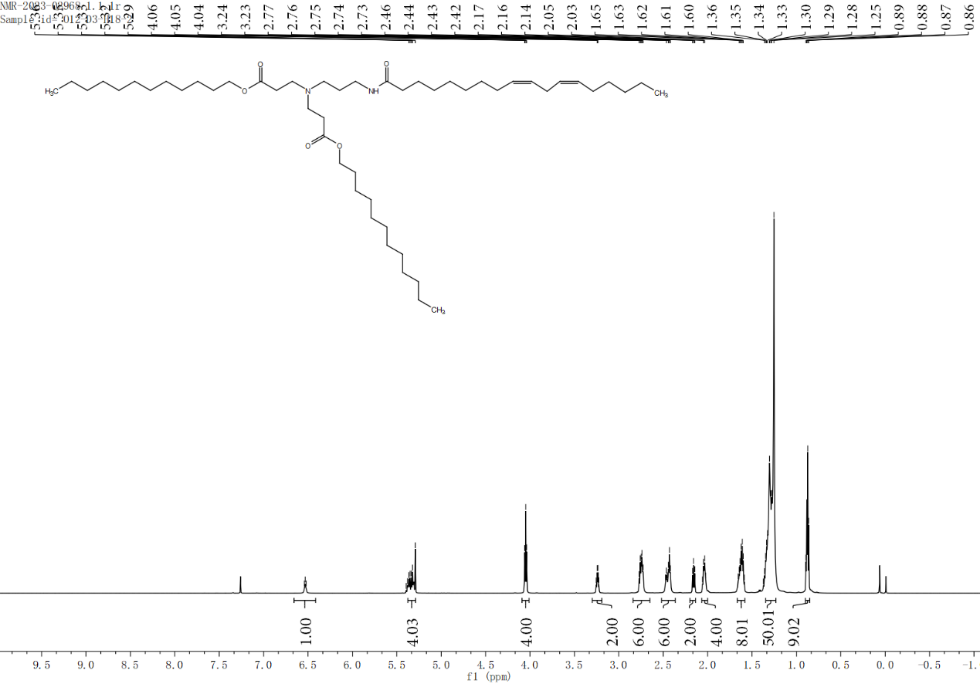

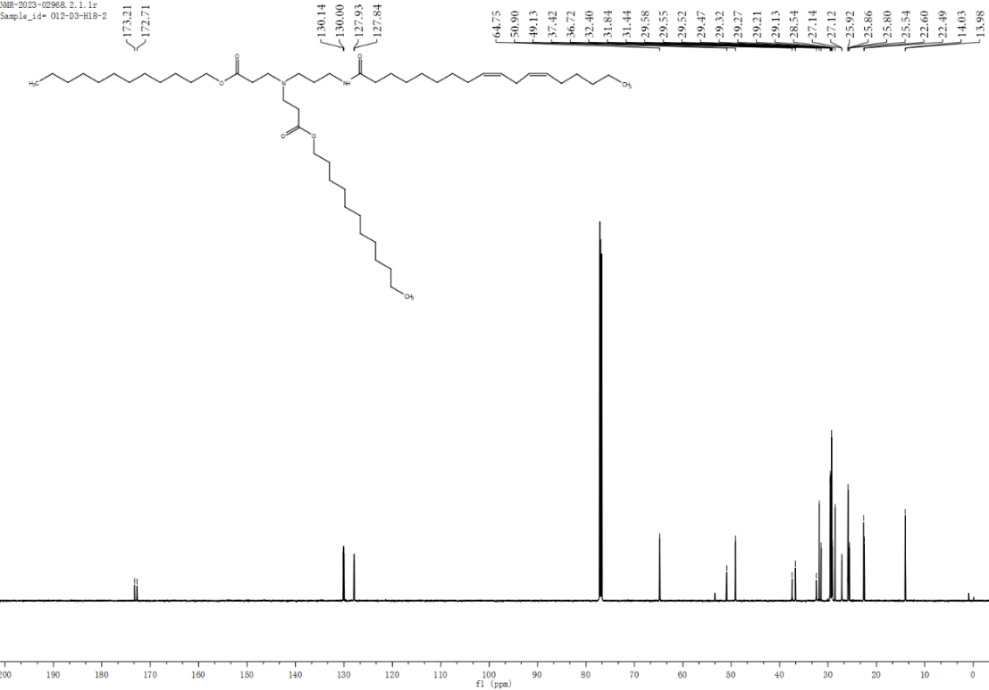

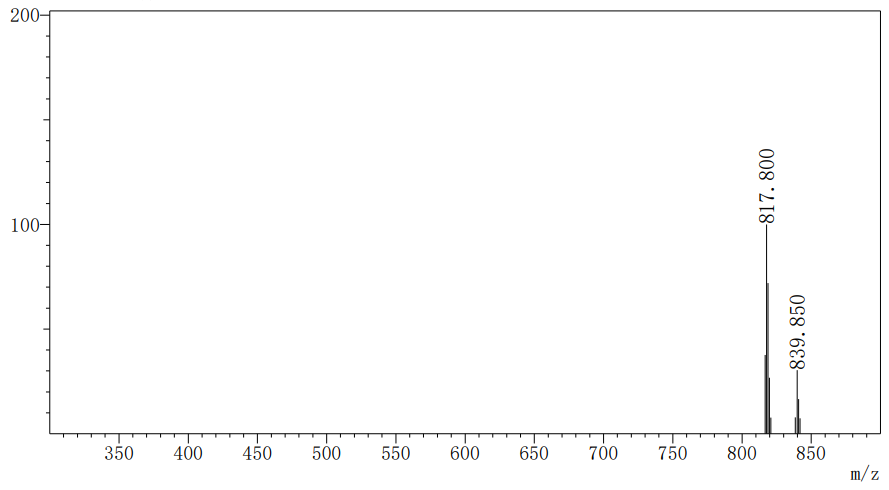

O12-D3-H18c

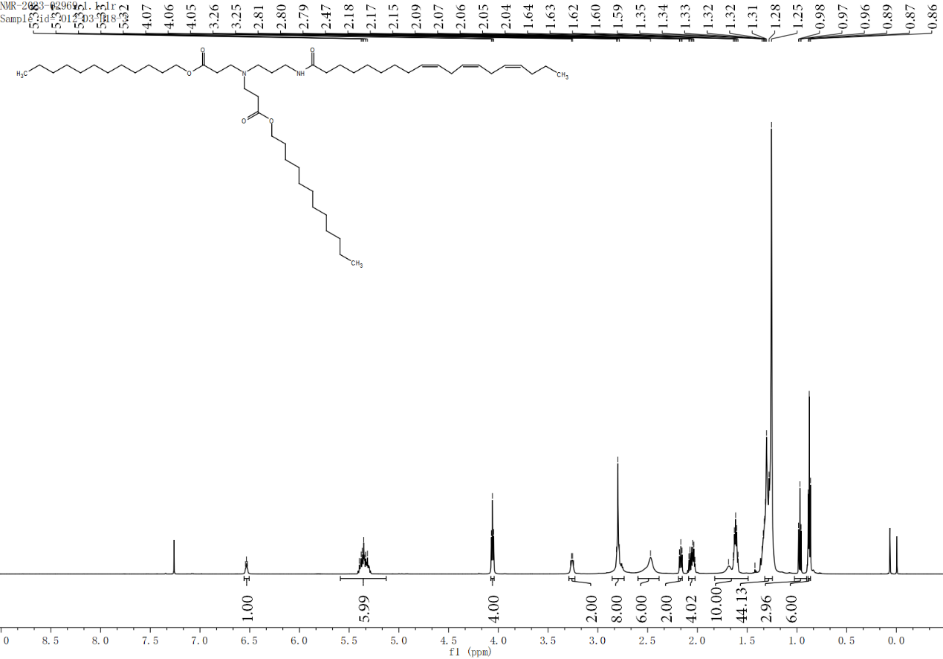

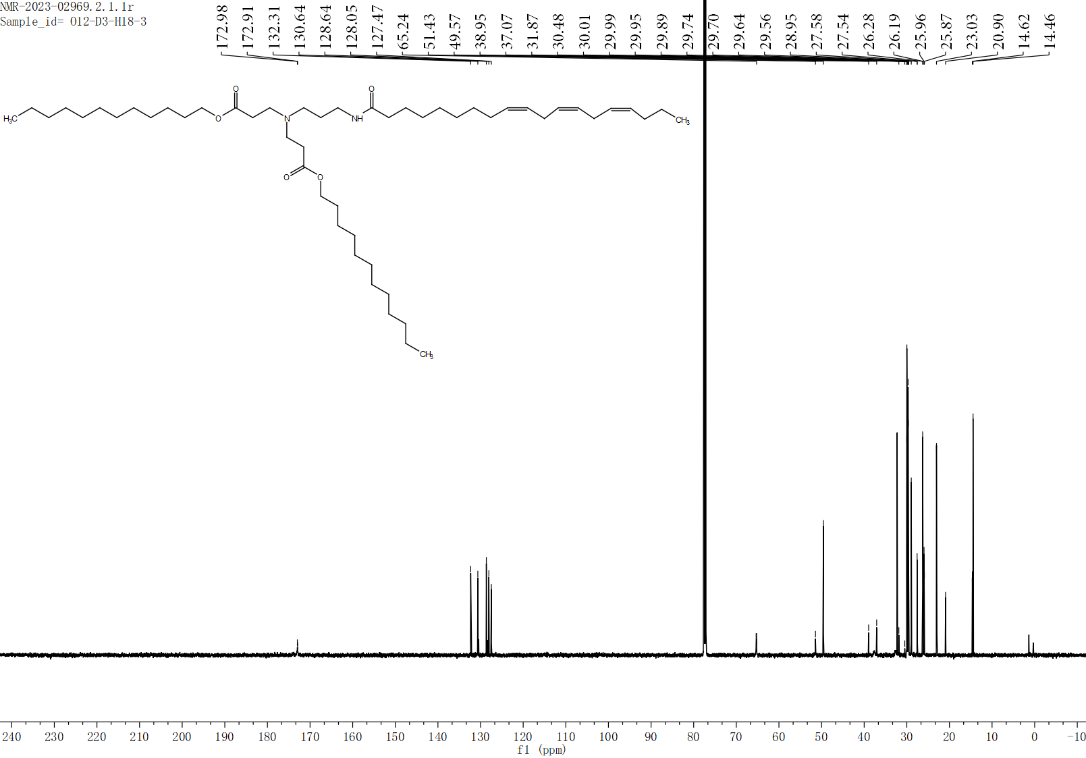

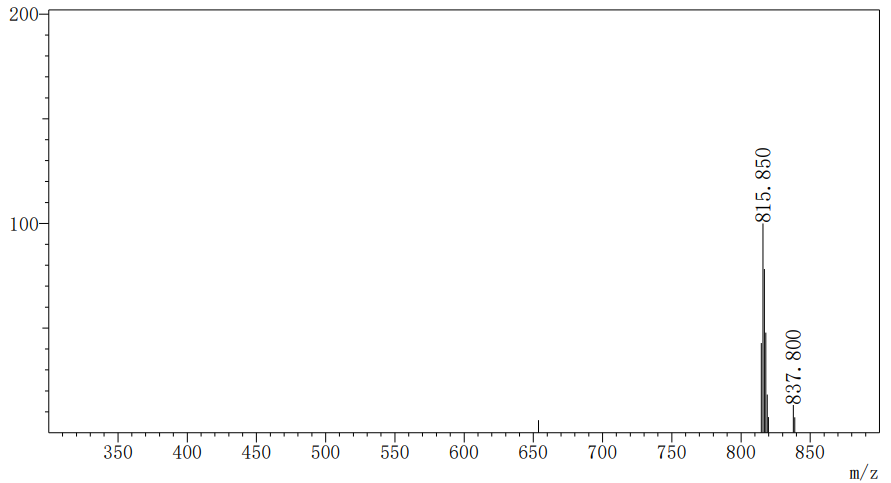

O12-D3-H9

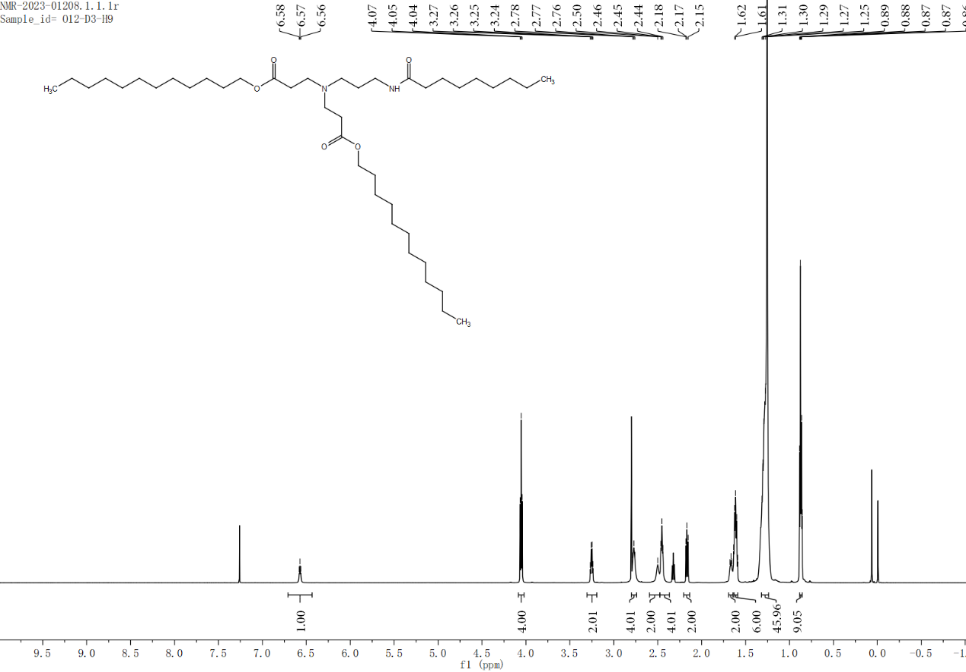

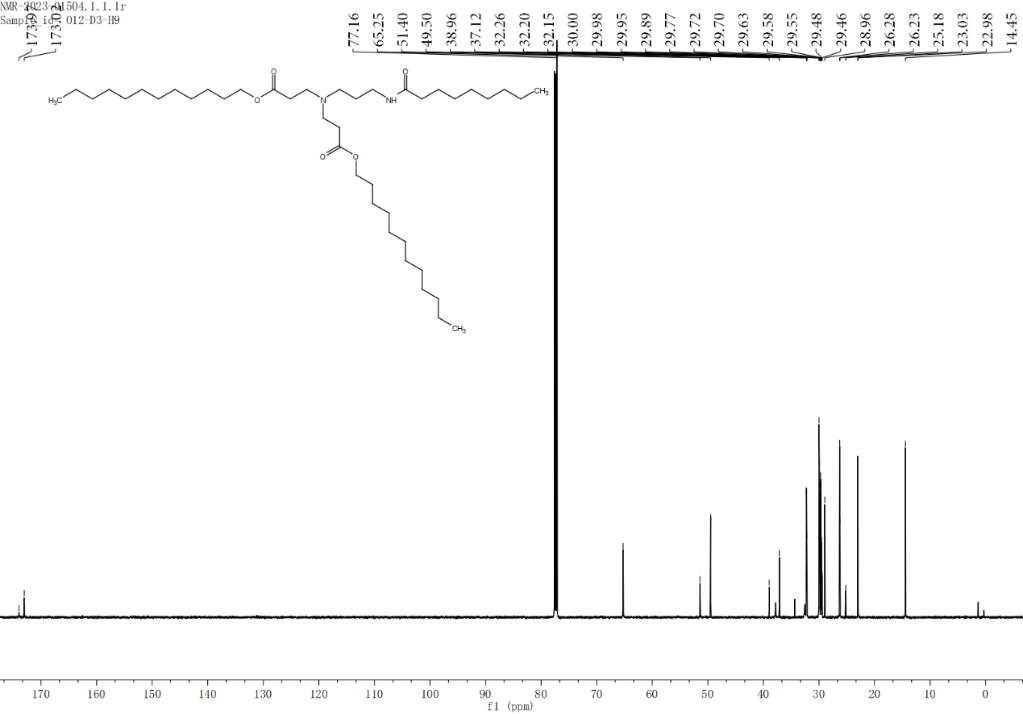

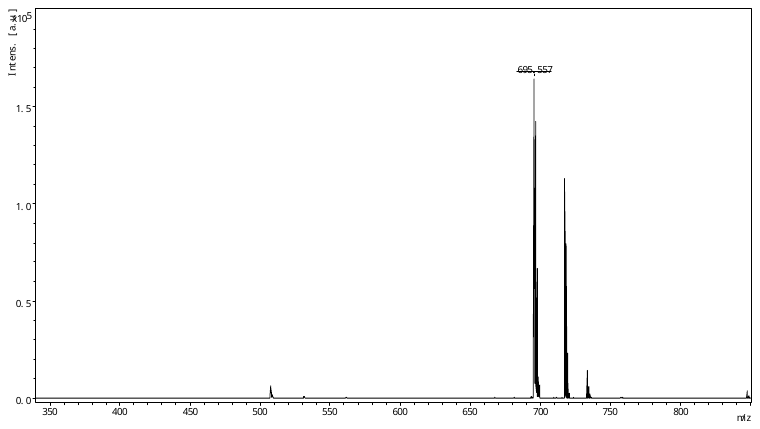

O12-D3-H16

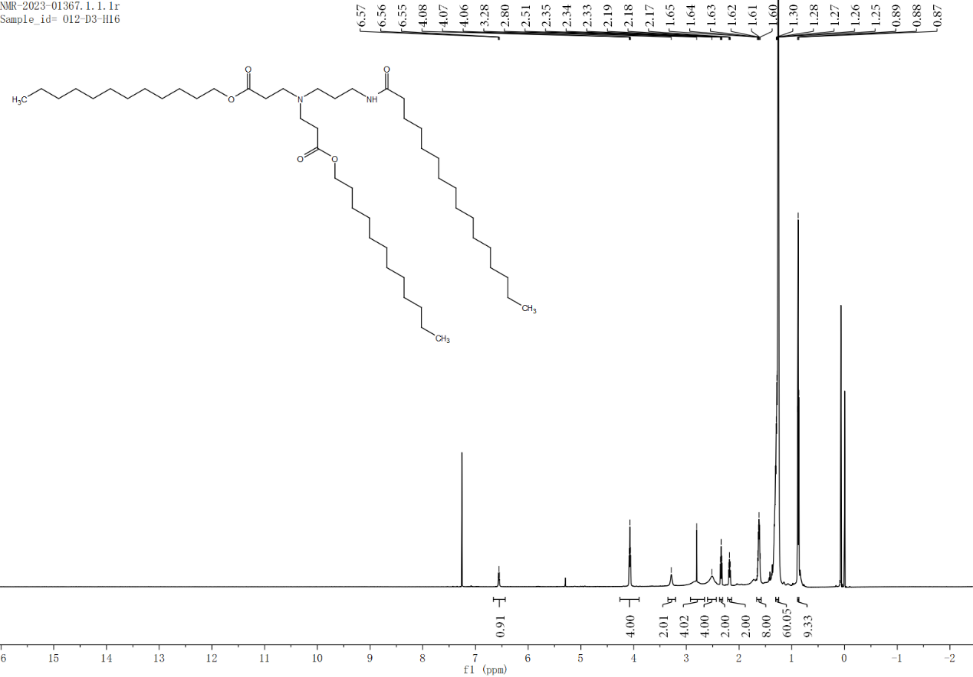

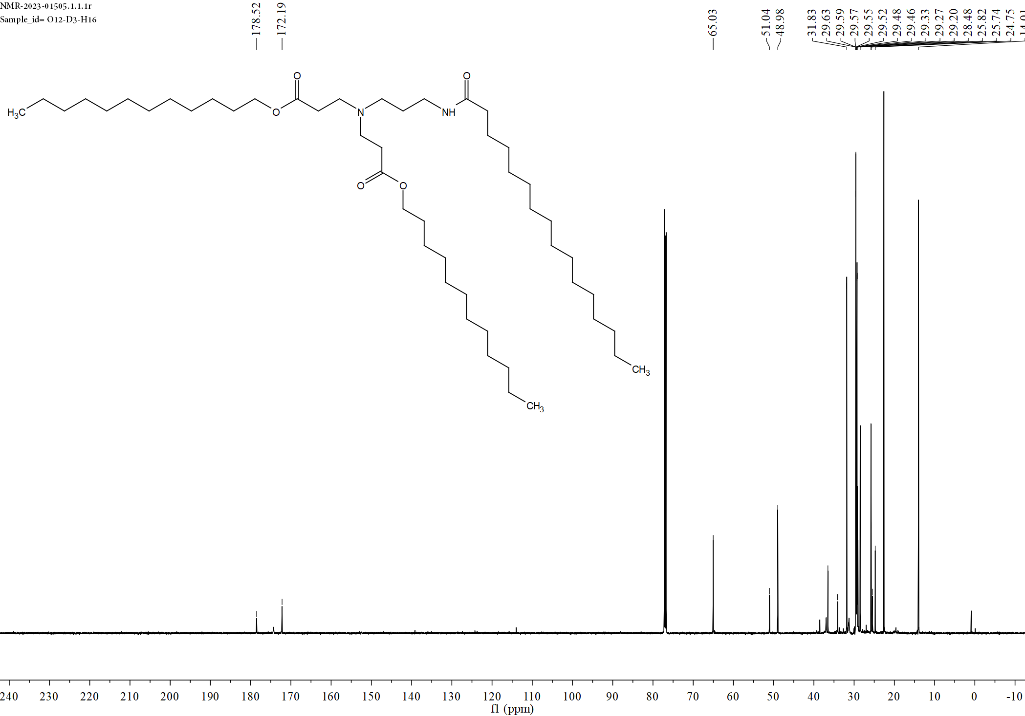

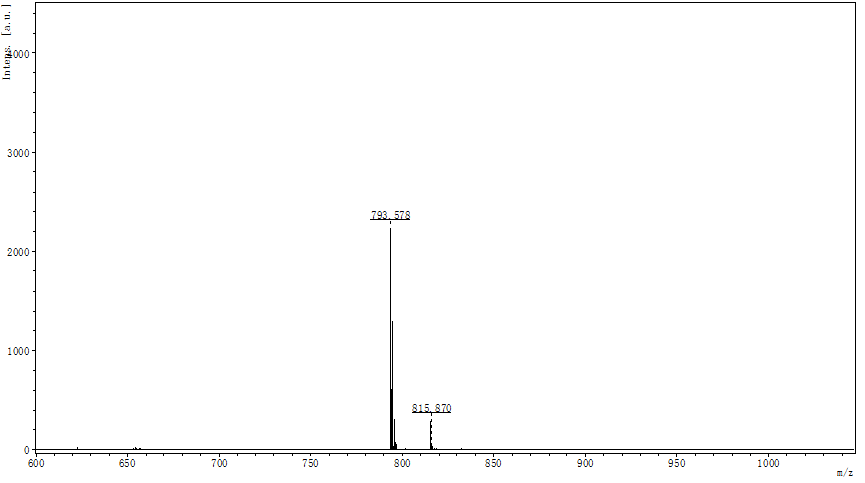

O12-D3-H17

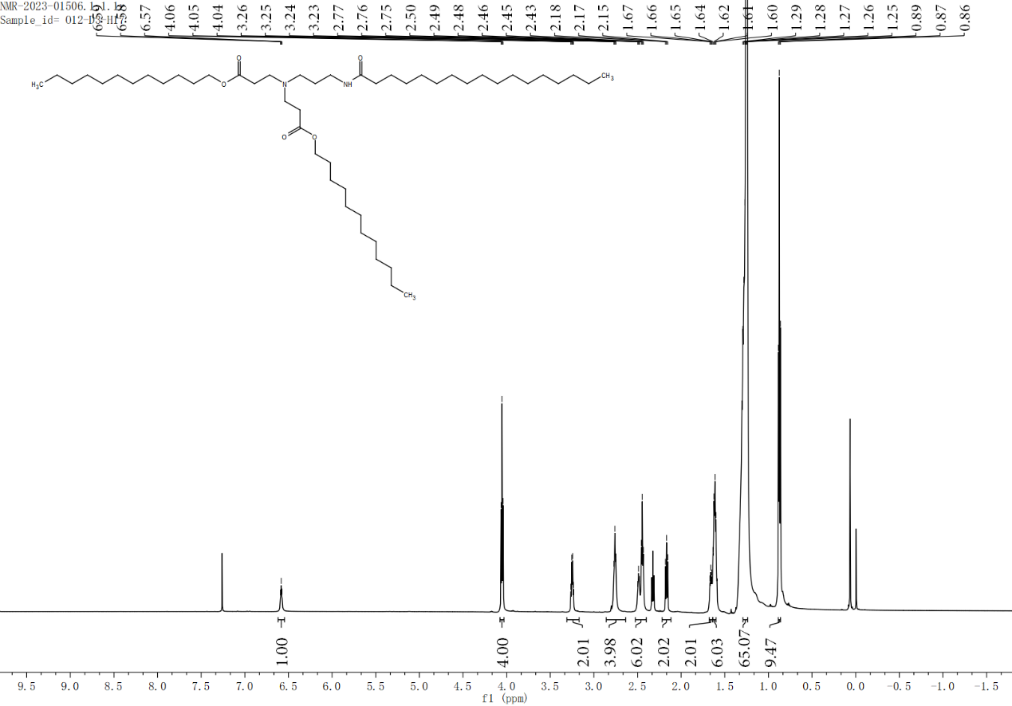

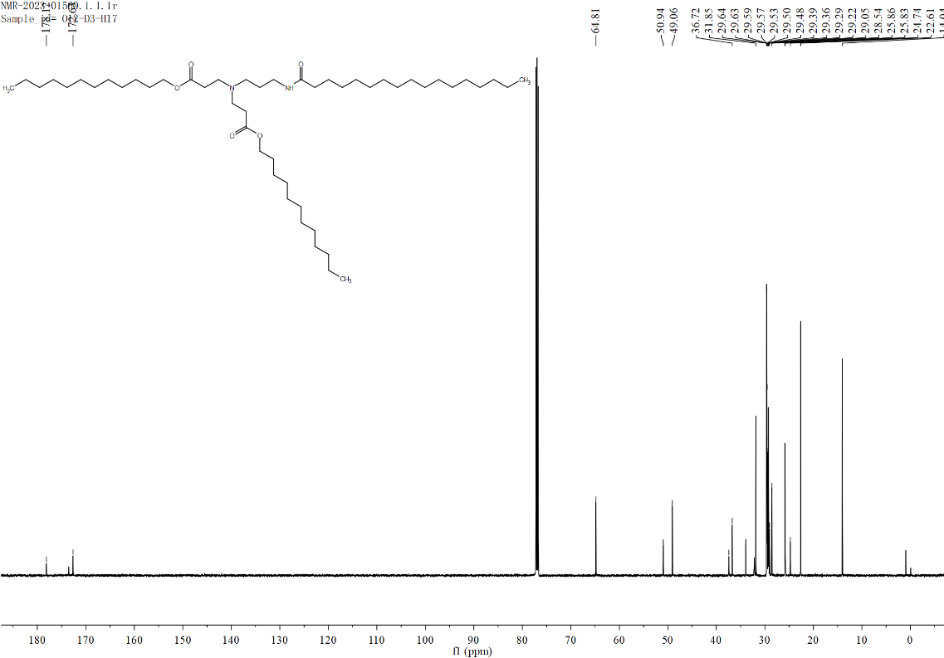

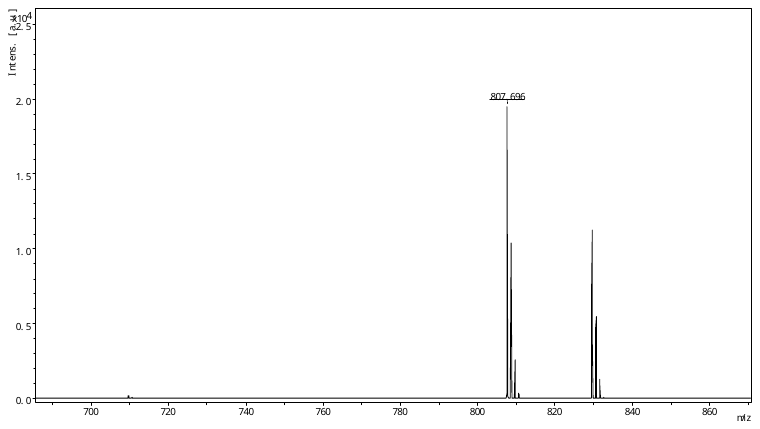

O12-D3-H18

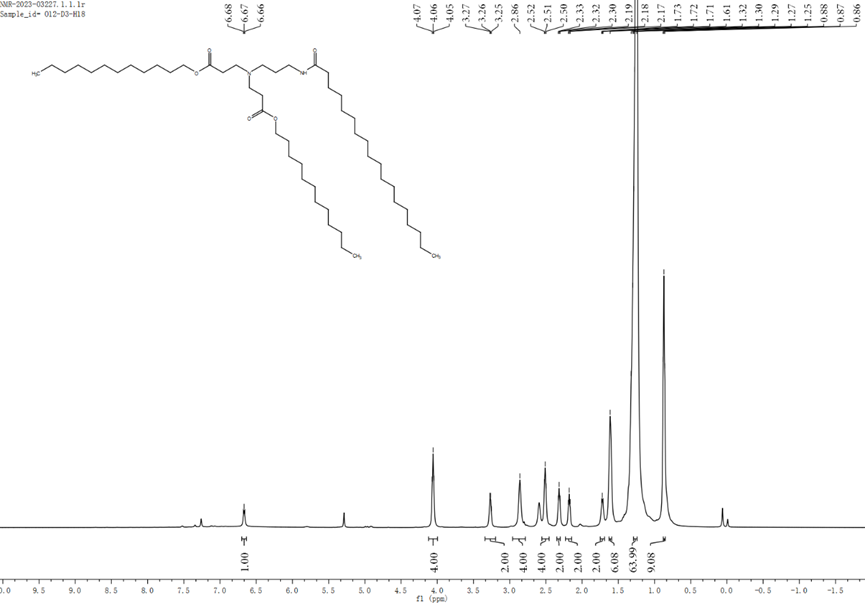

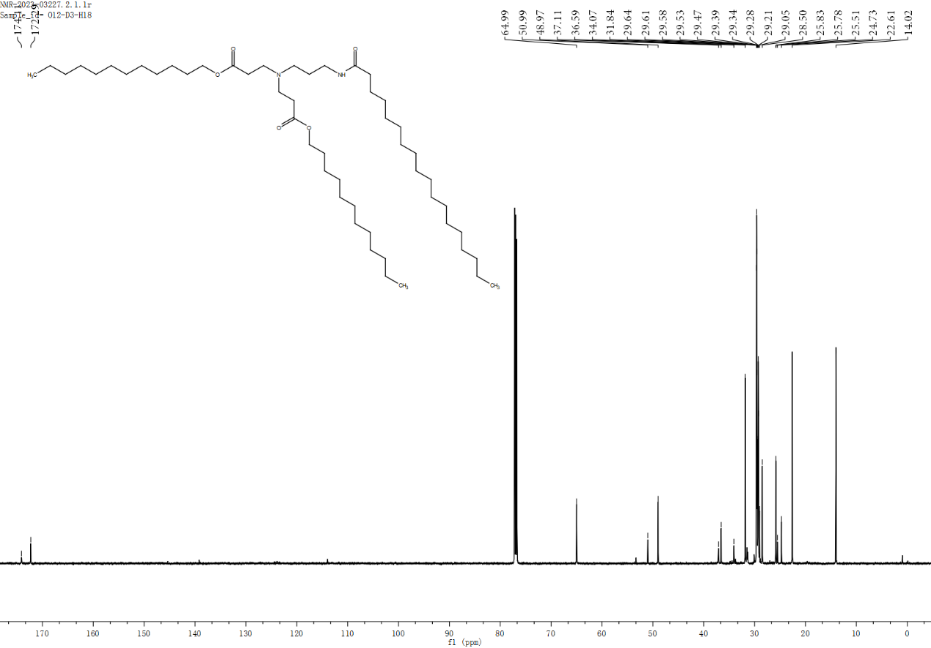

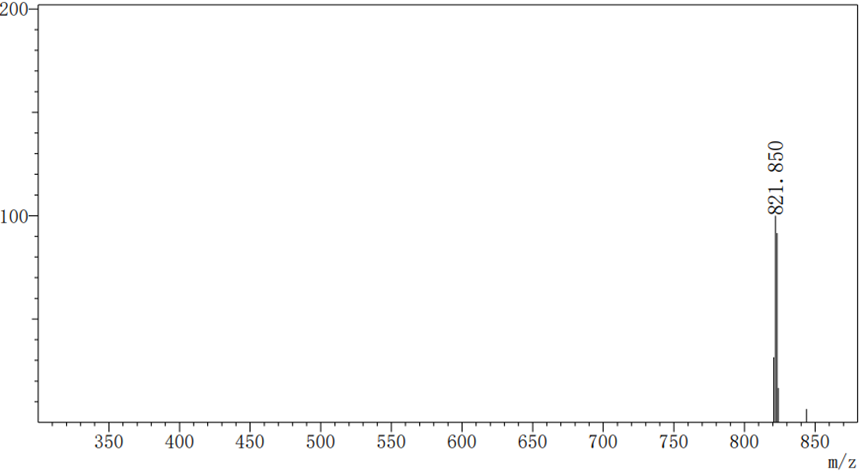

O14-D3-H9

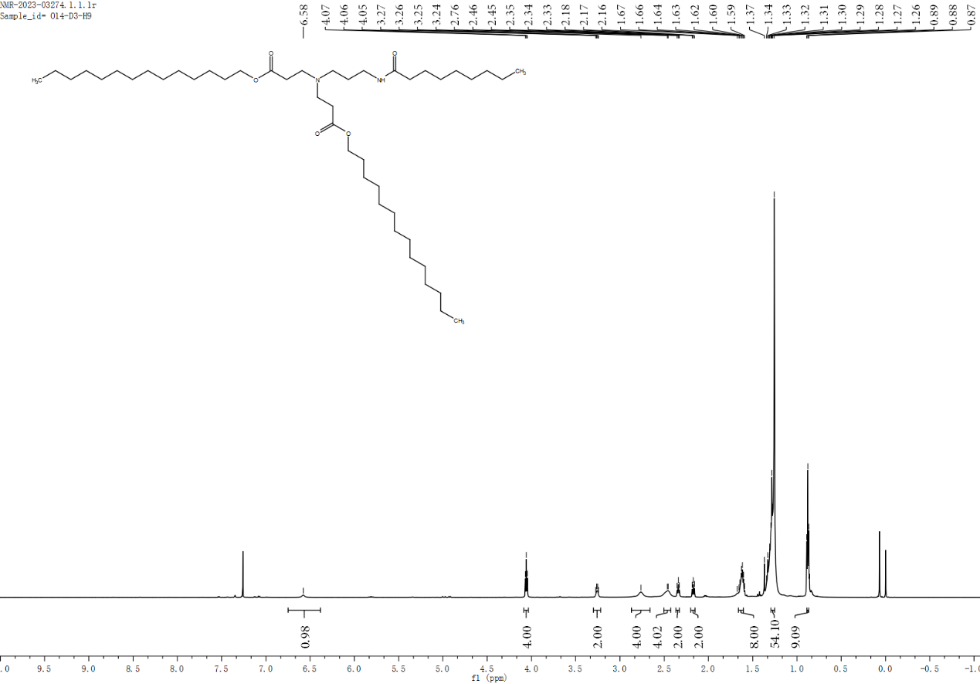

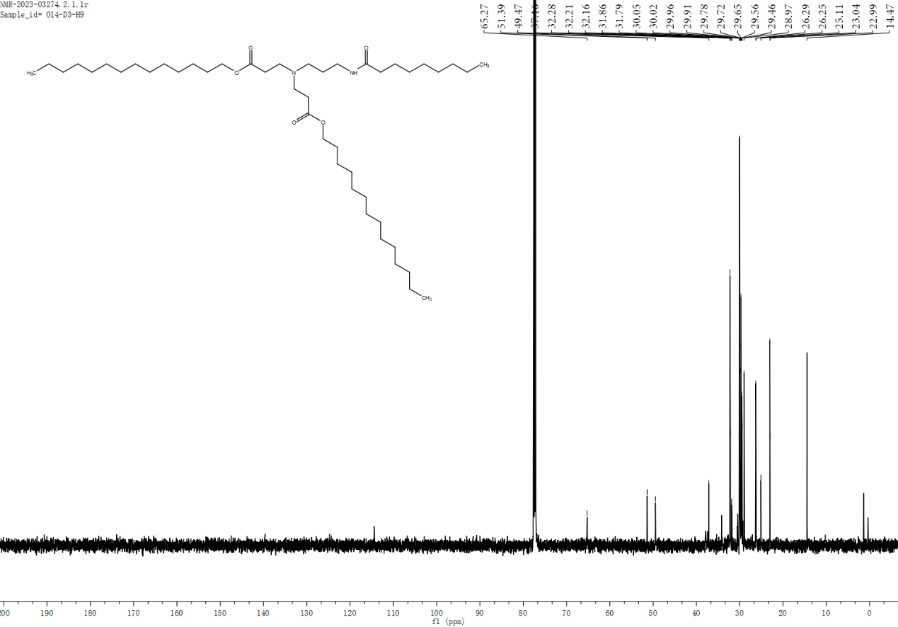

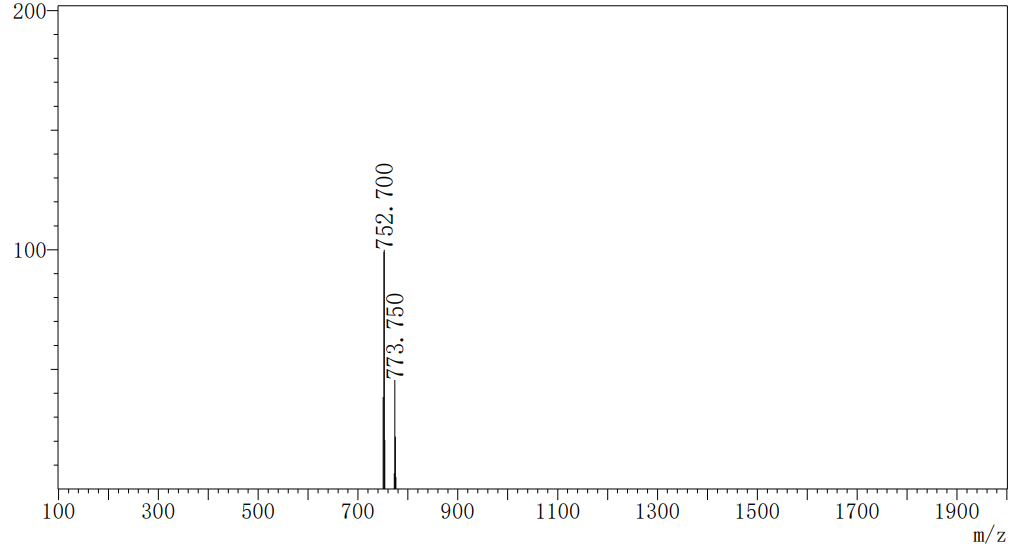

O14-D3-H13

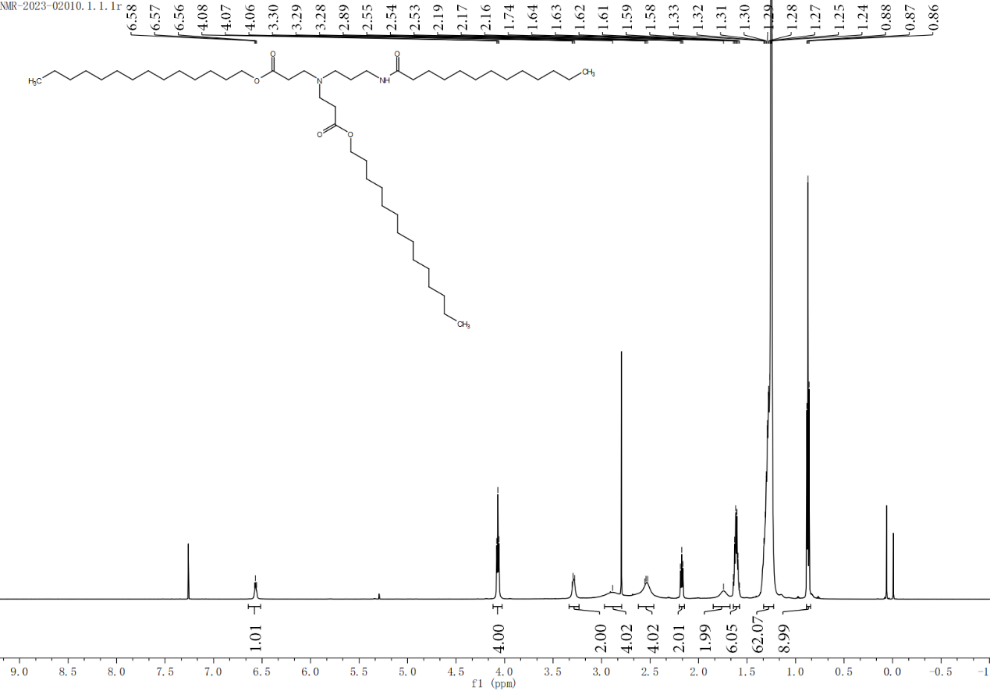

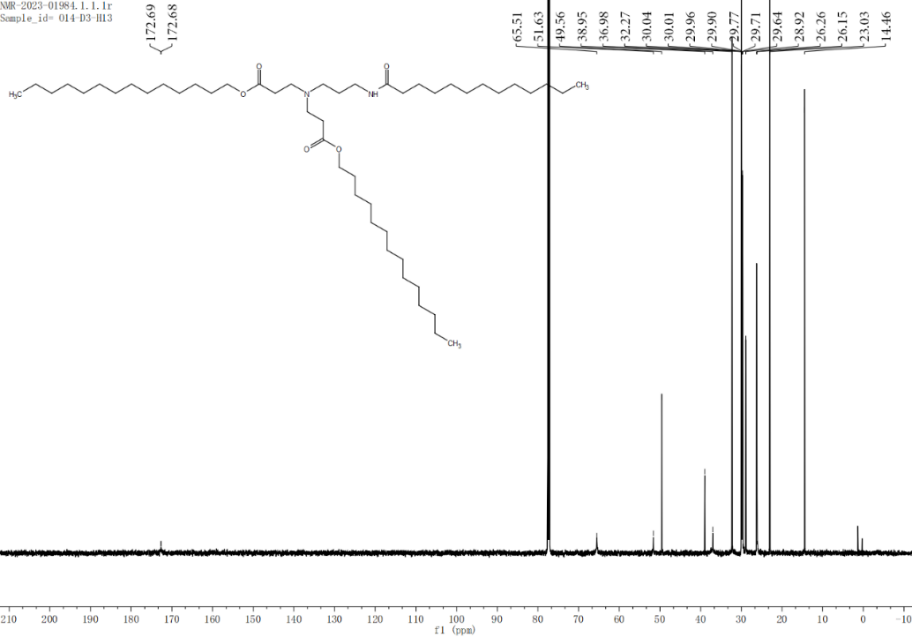

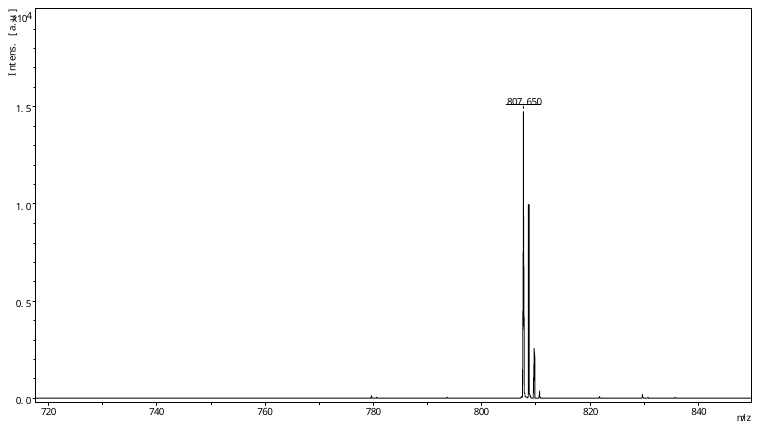

O14-D3-H14

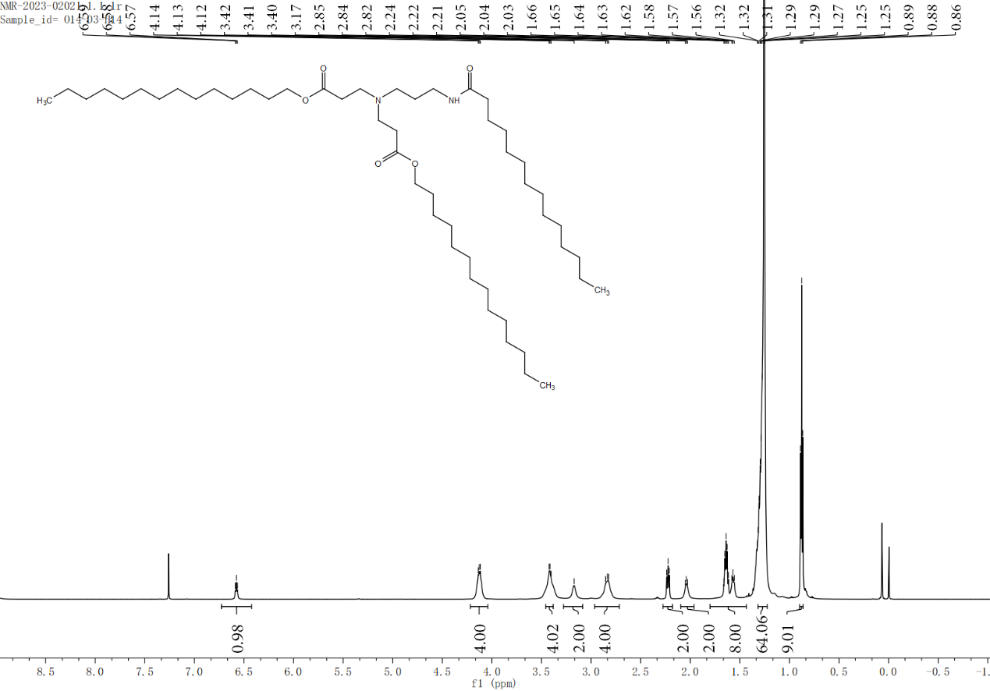

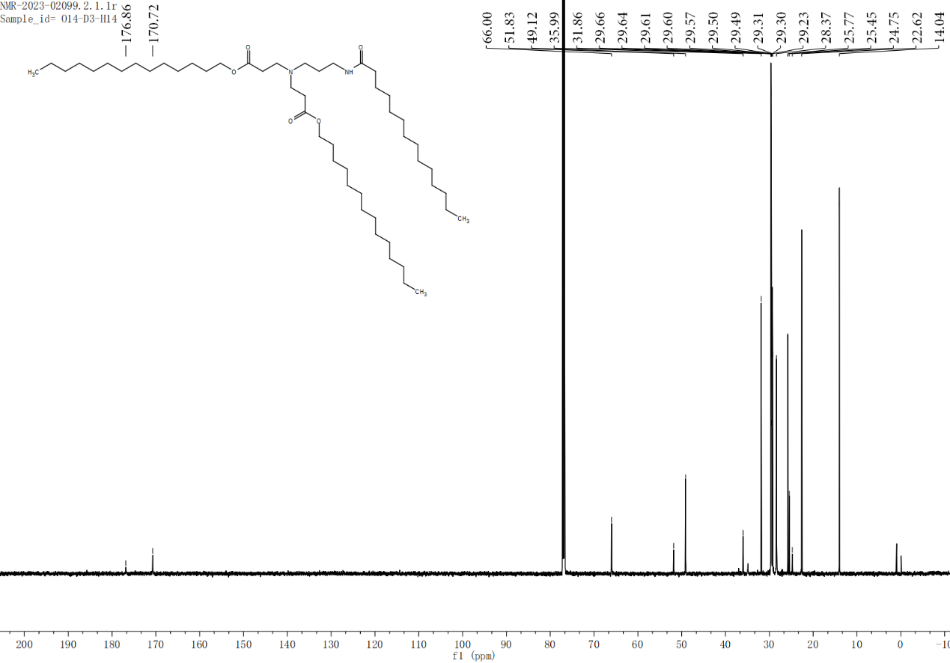

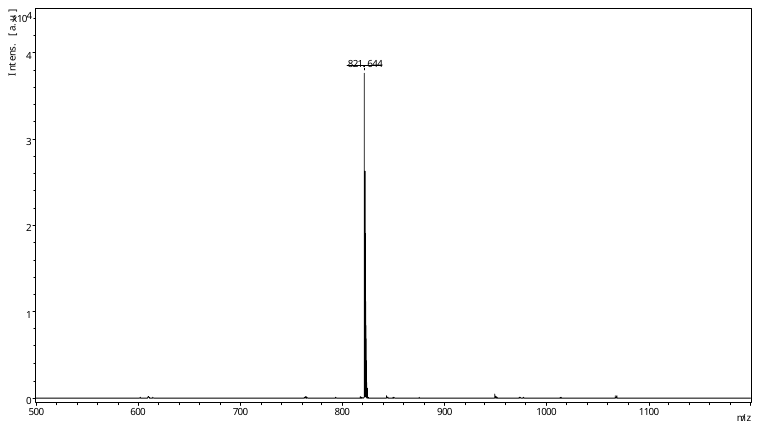

O14-D3-H15

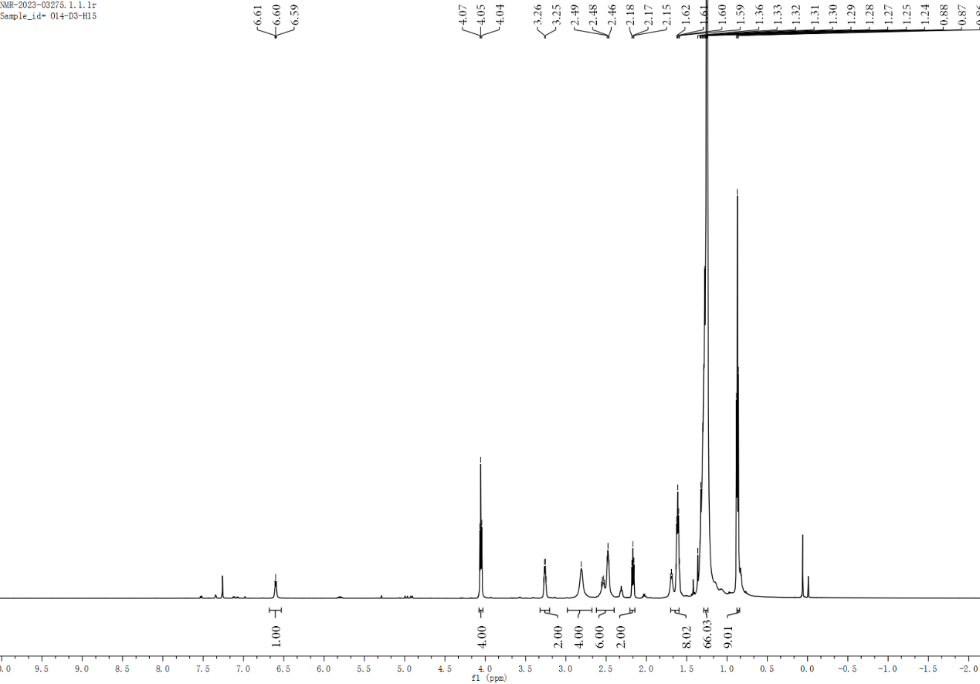

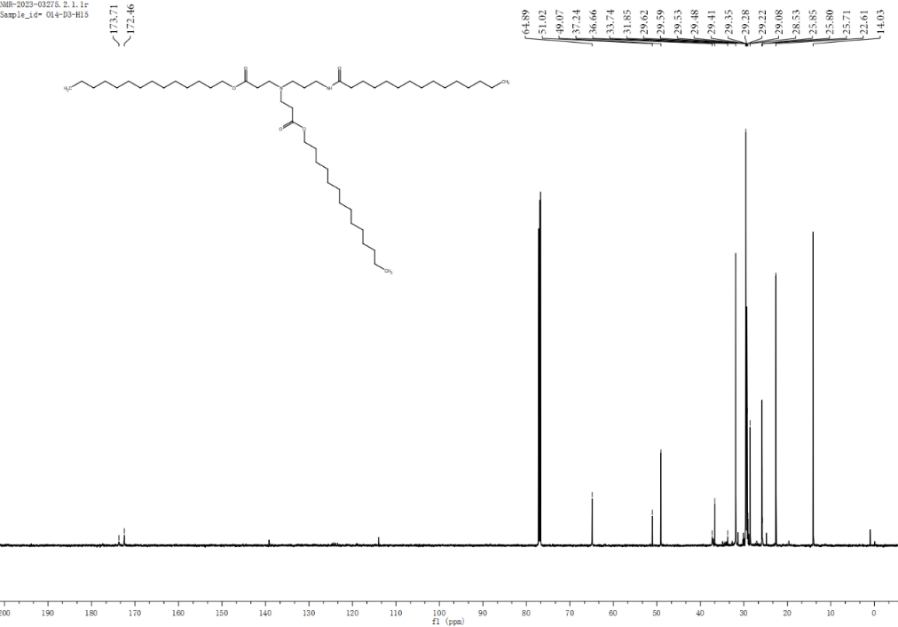

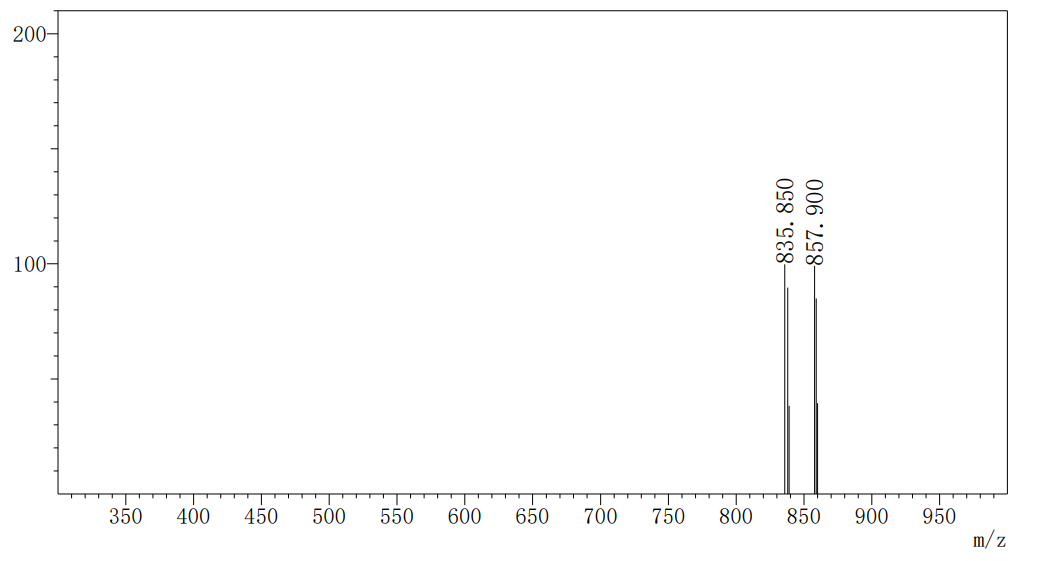

O16-D3-H9

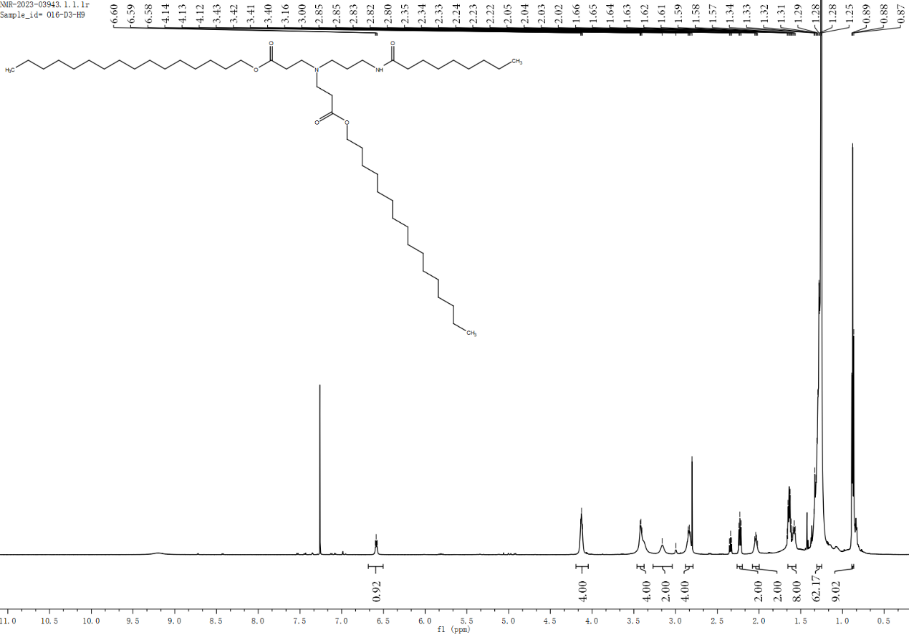

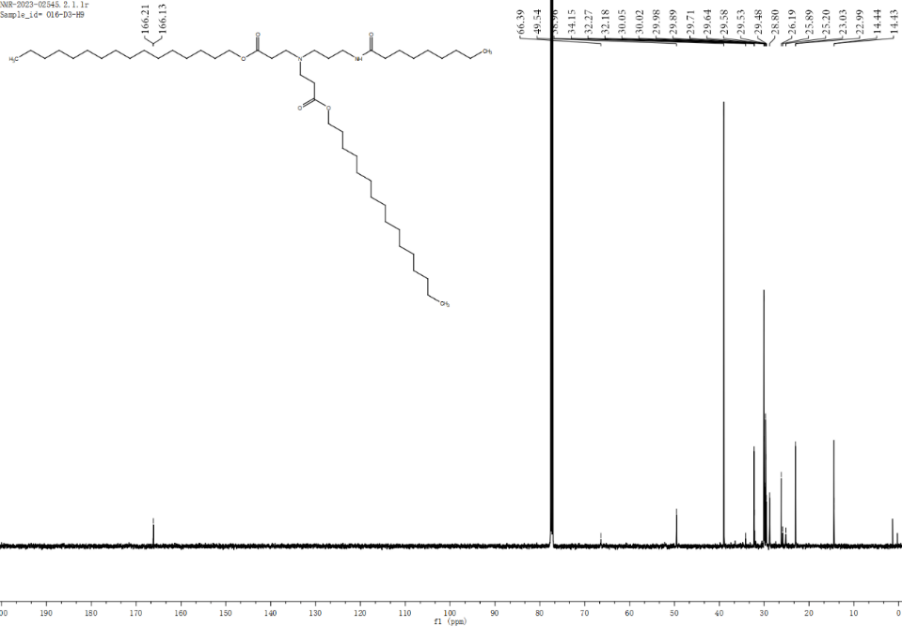

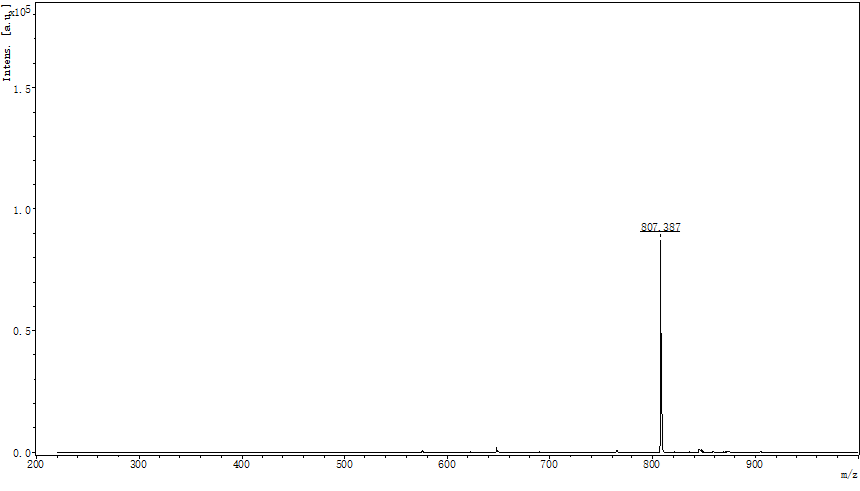

O16-D3-H10

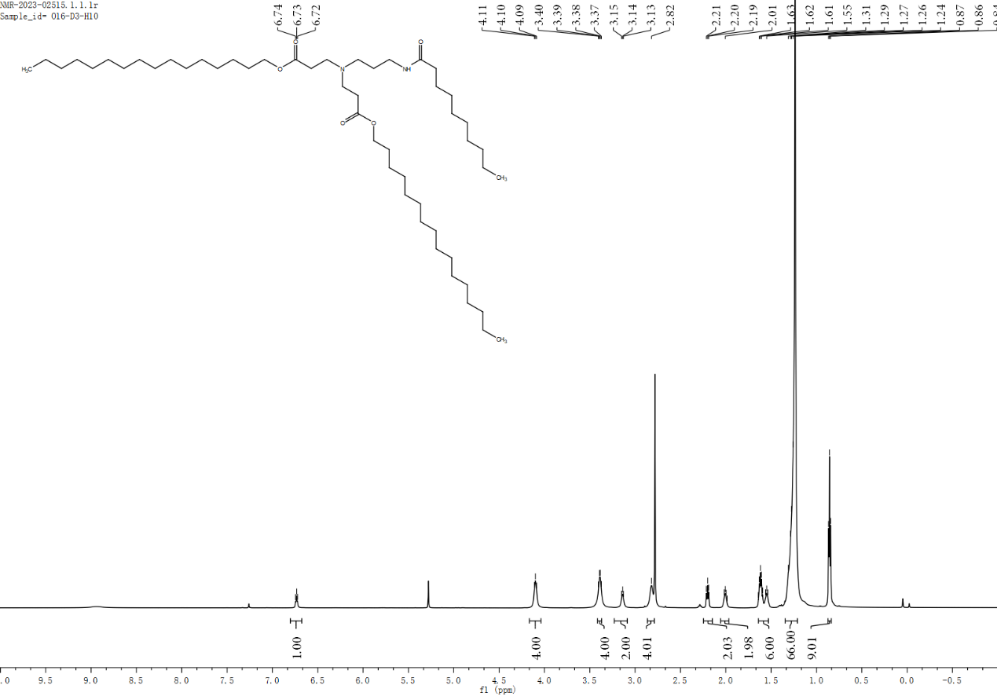

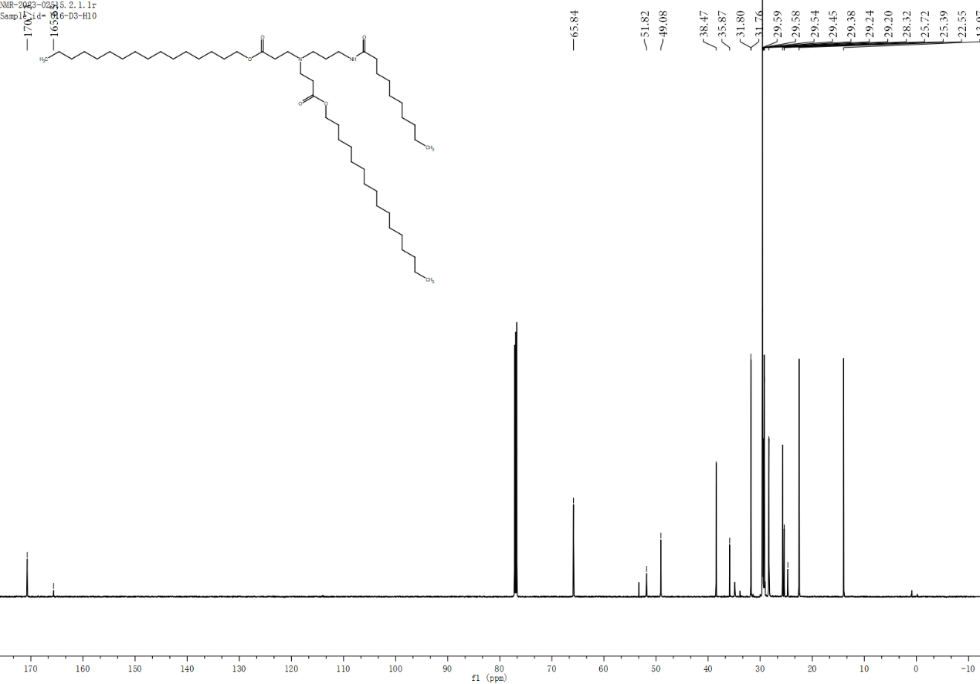

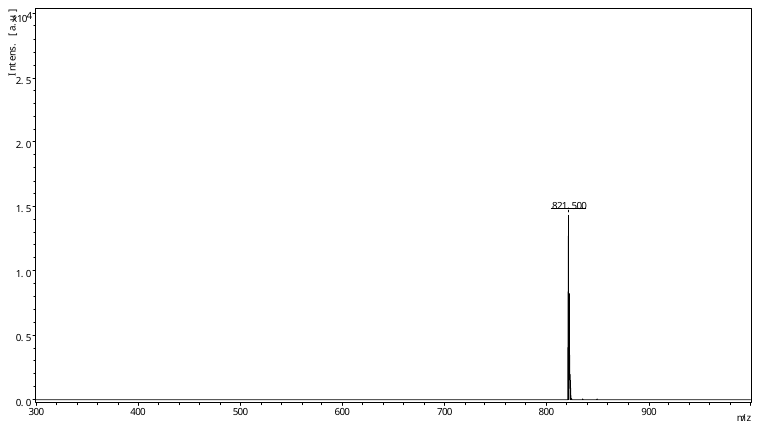

O16-D3-H11

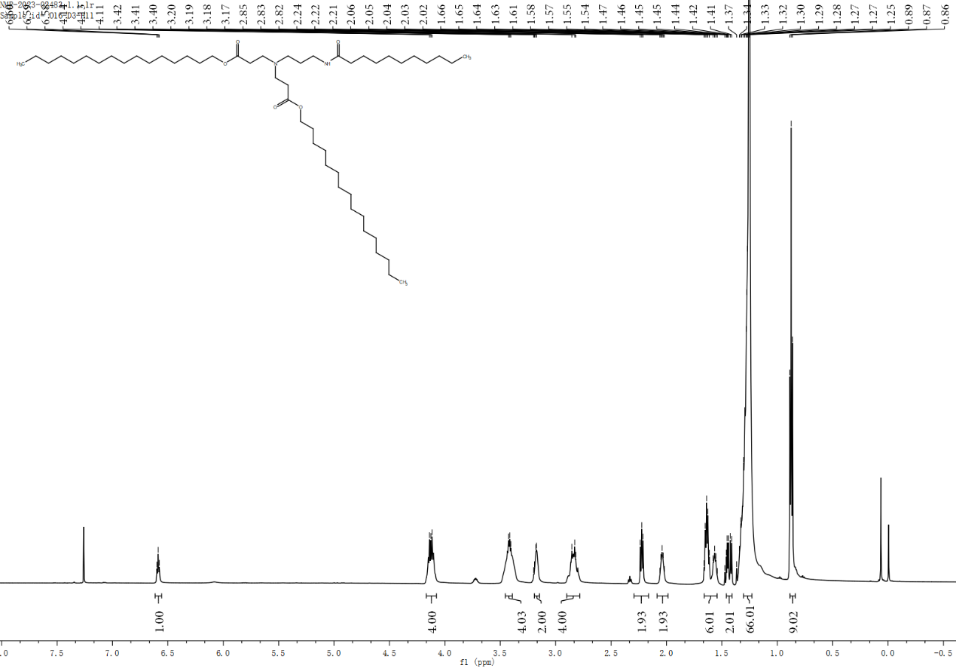

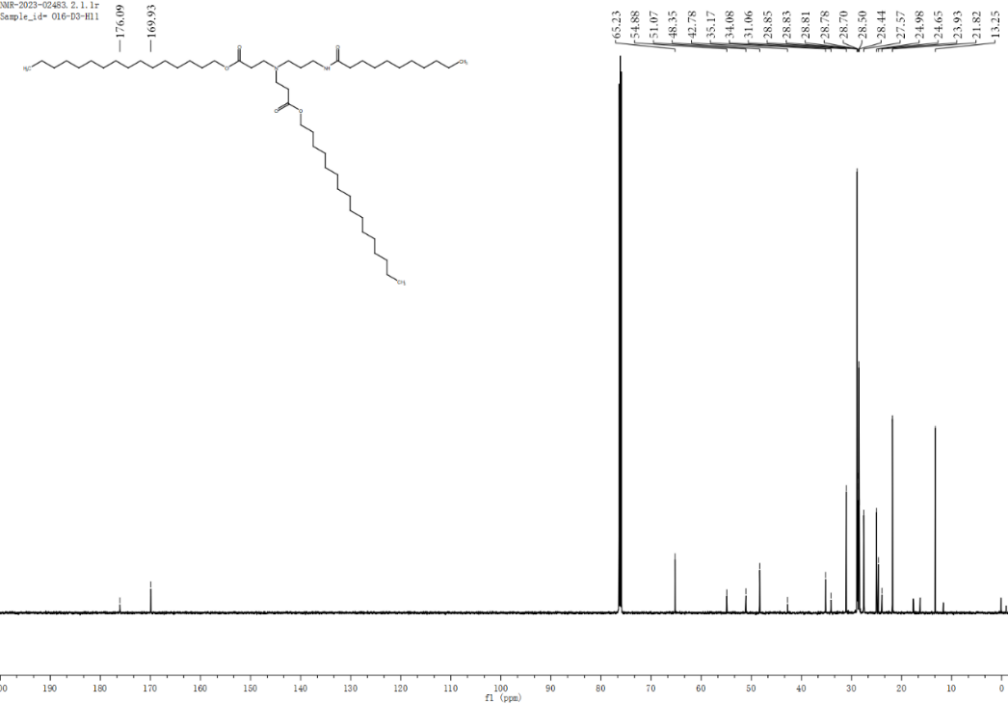

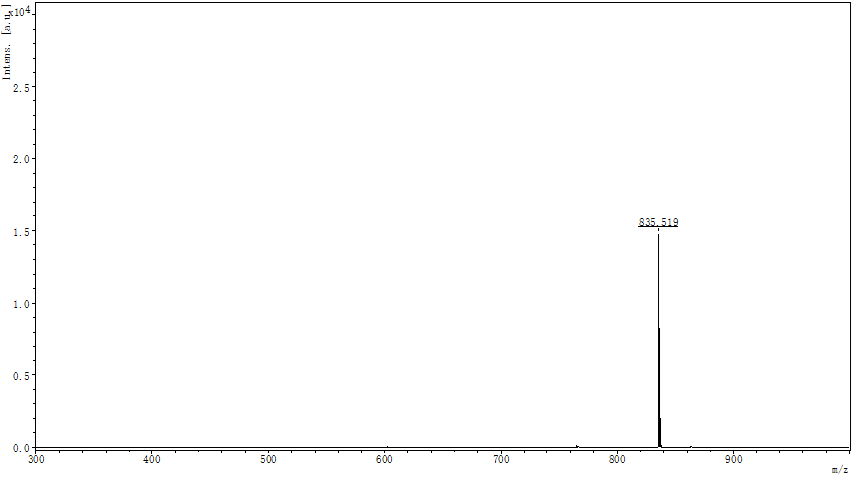

O16-D3-H12

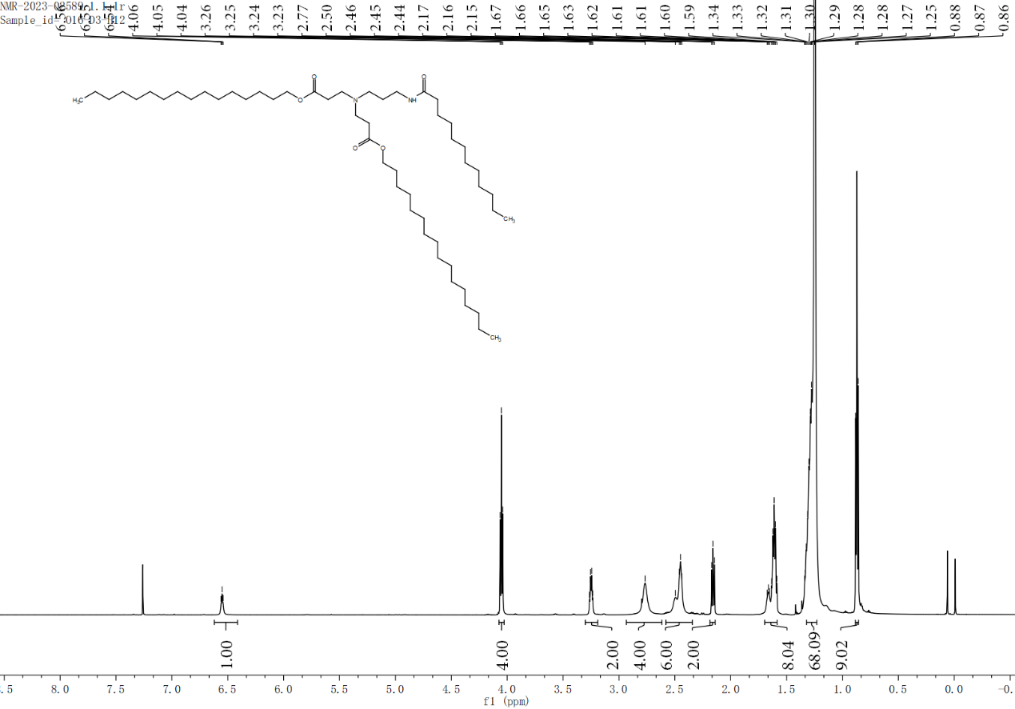

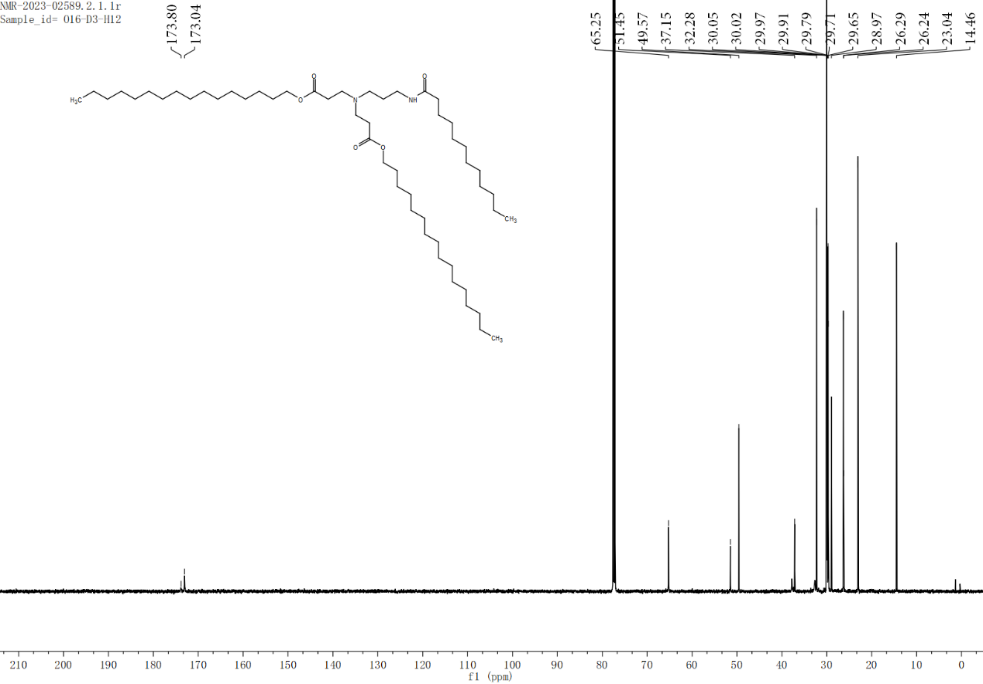

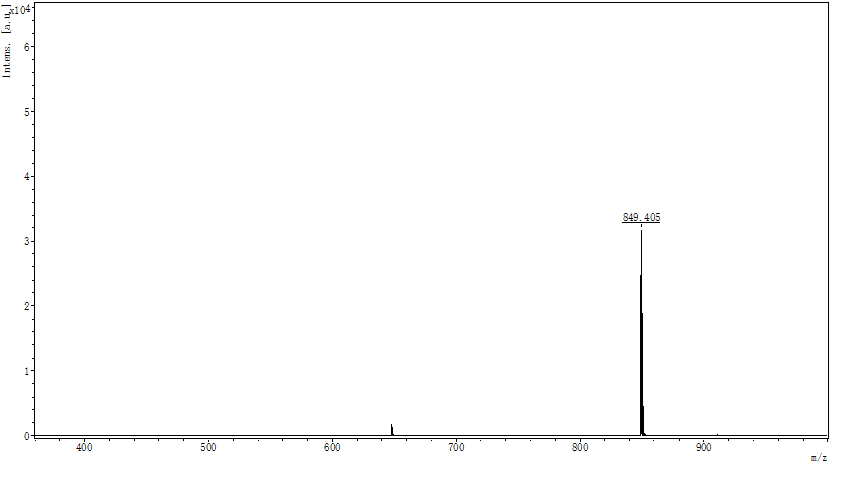

O12-K6-H7

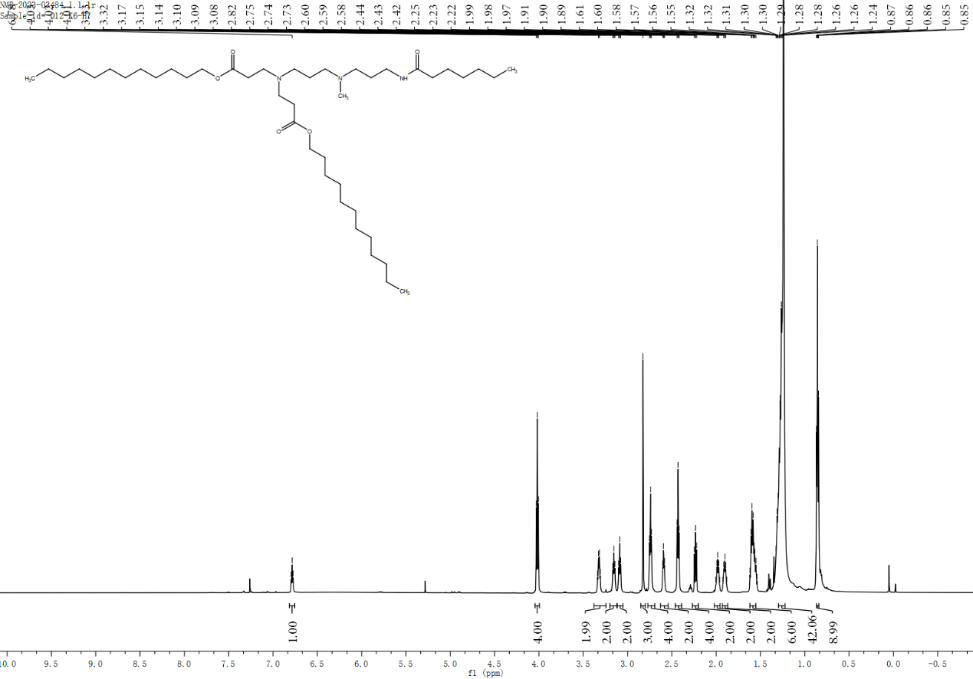

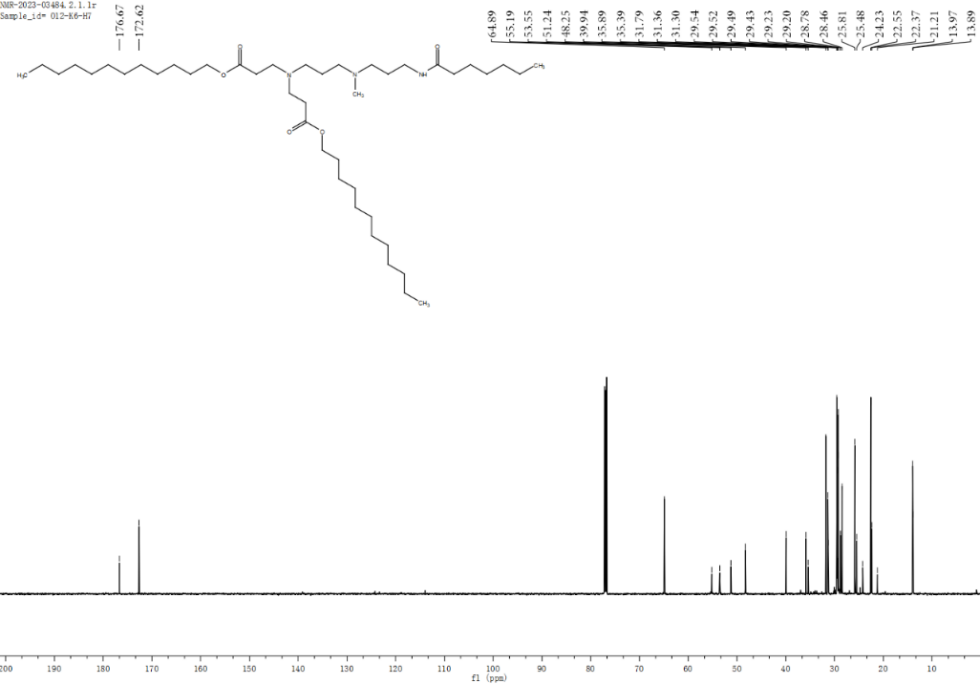

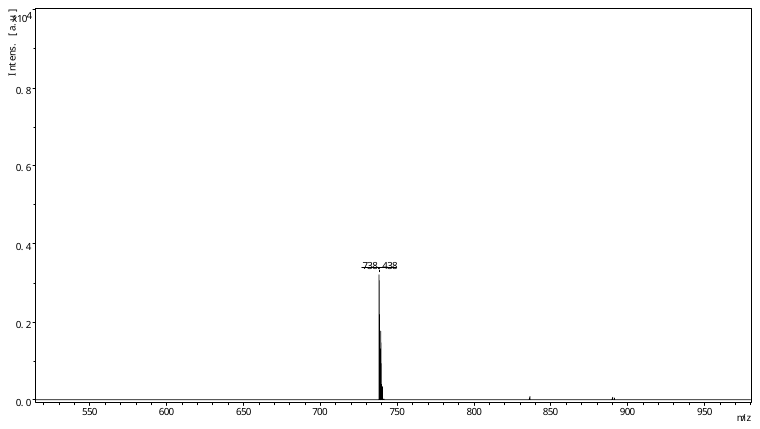

O12-K6-H8

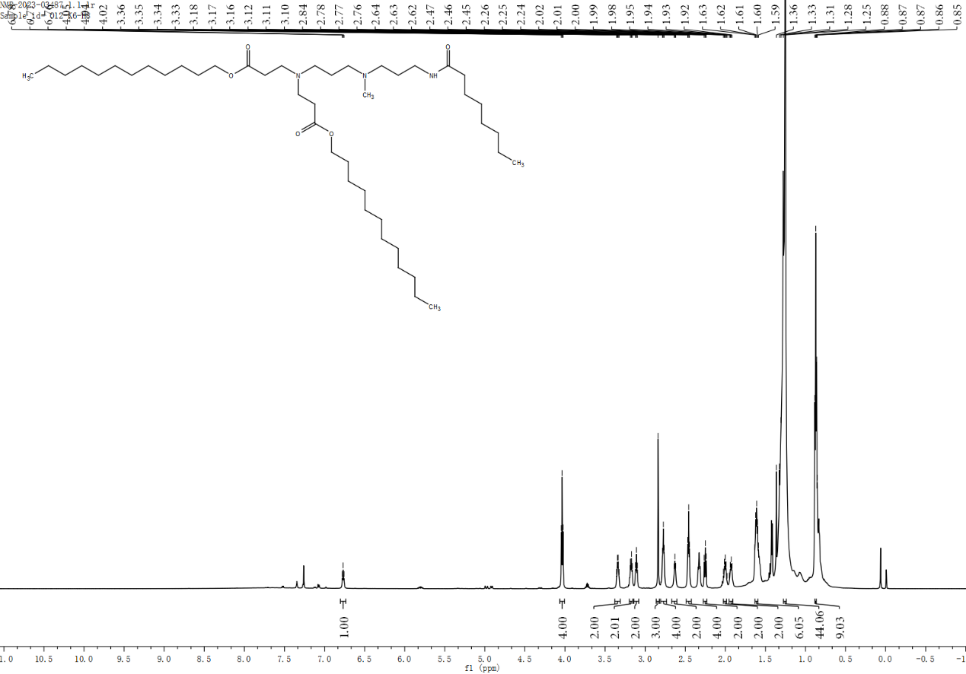

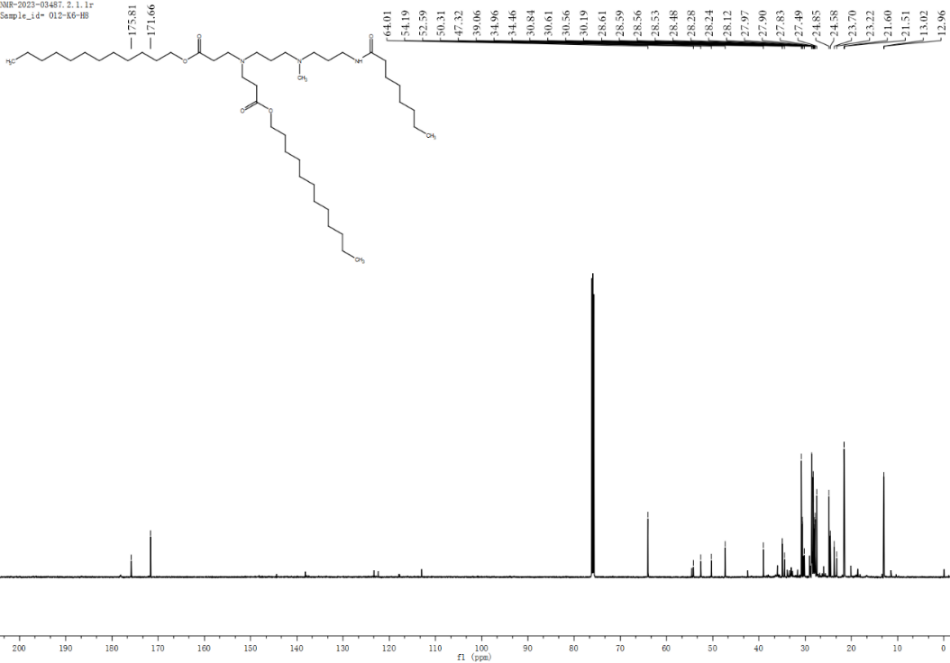

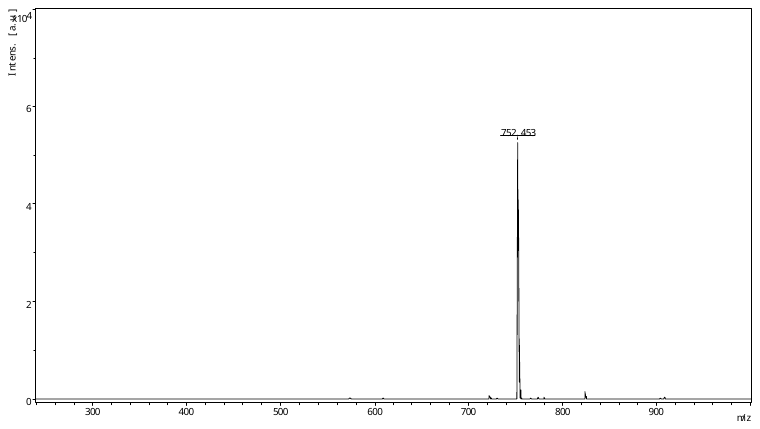

O12-K6-H9

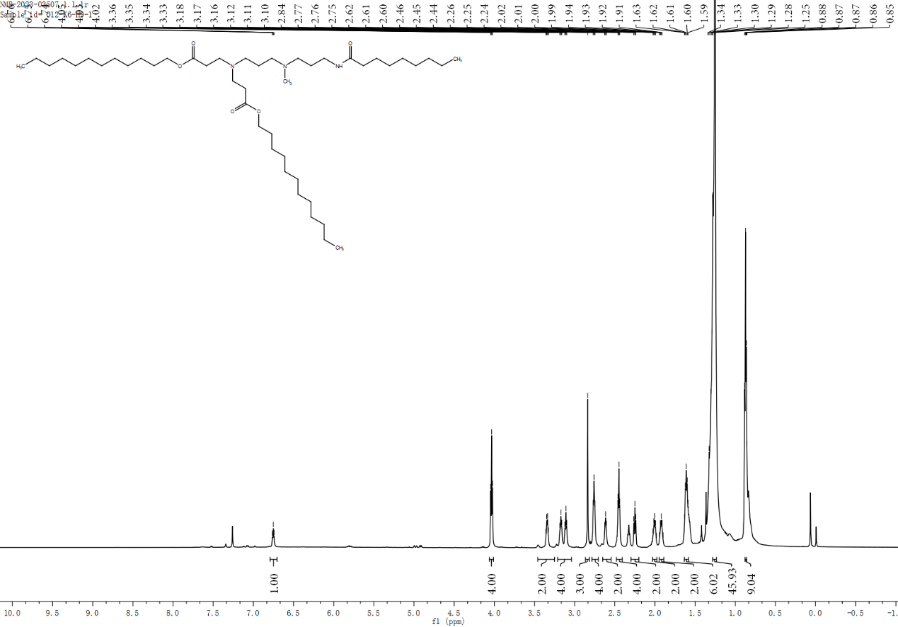

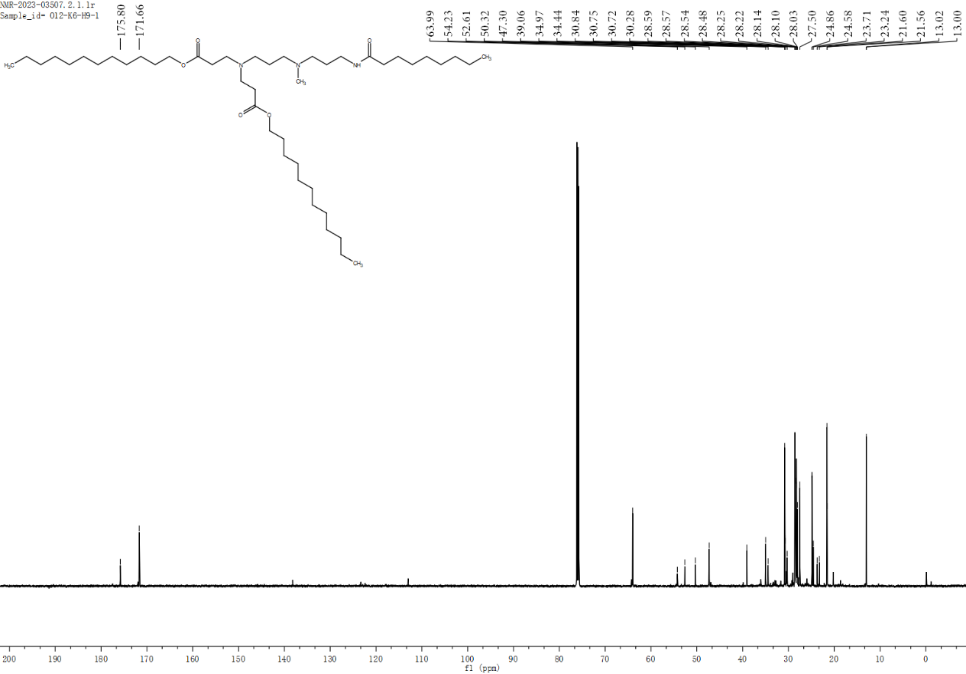

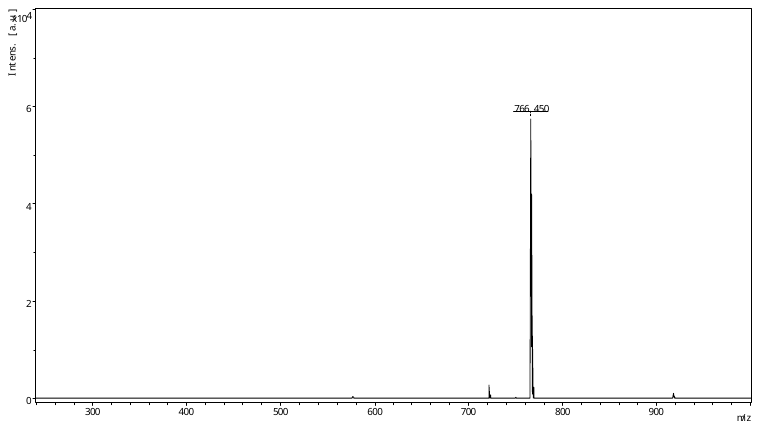

O12-K6-H16

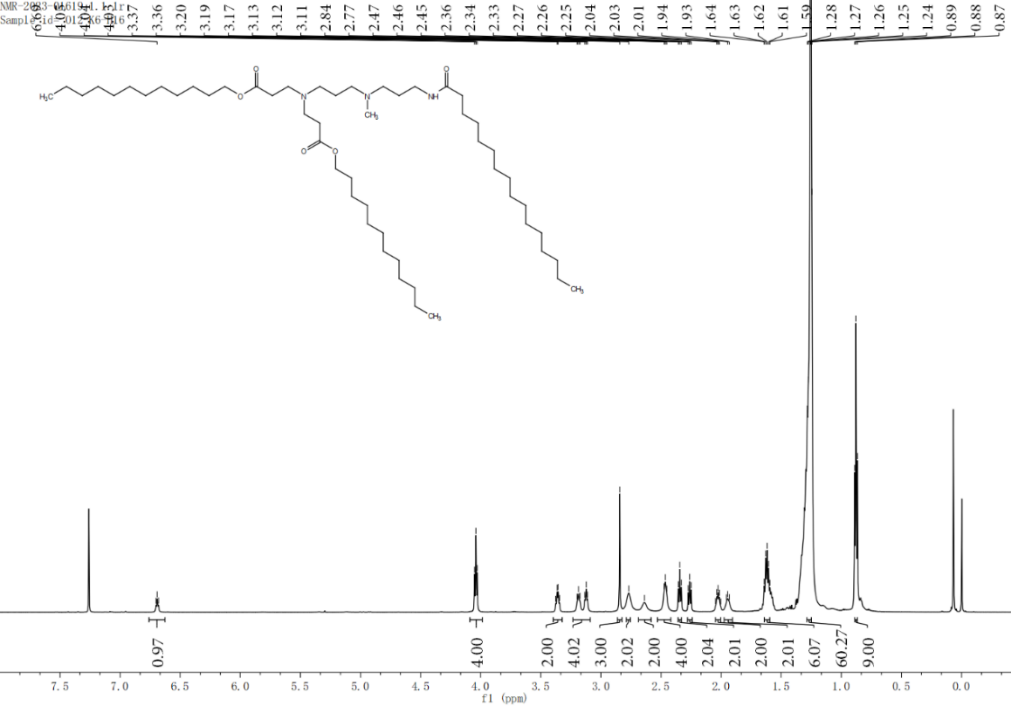

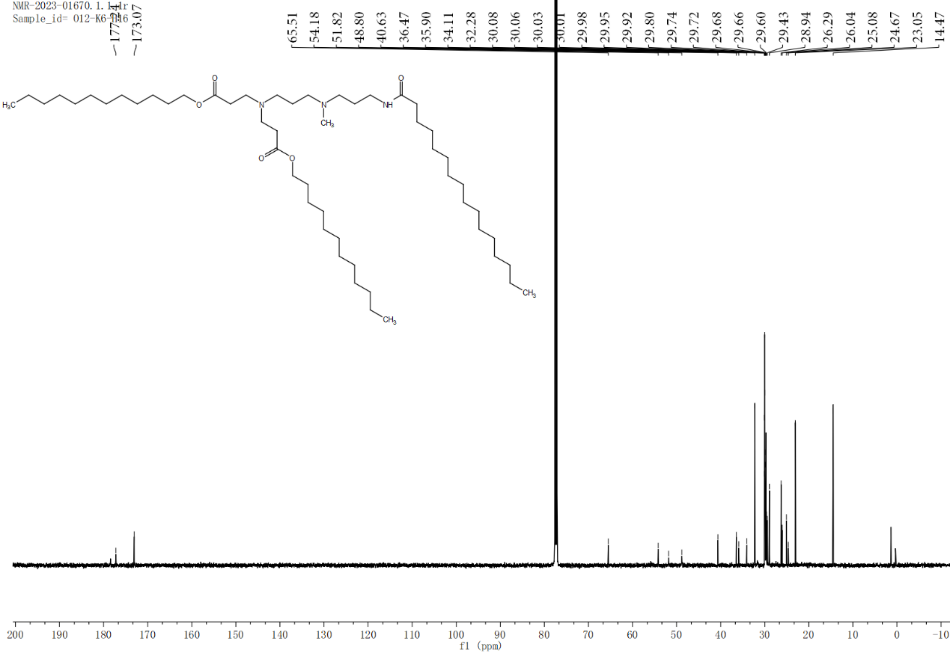

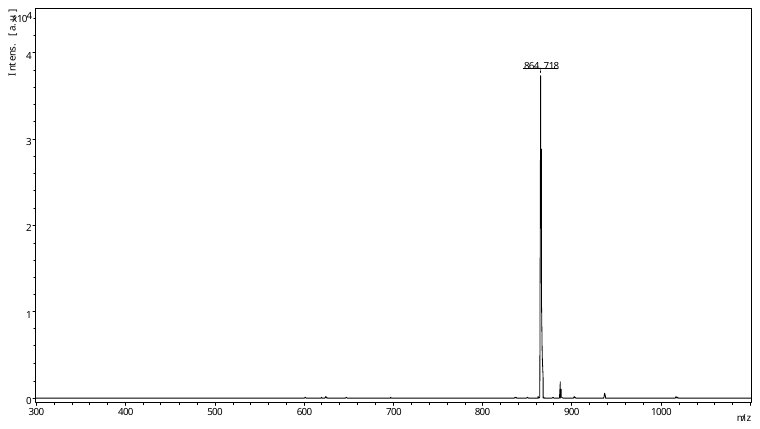

O12-P6-H7

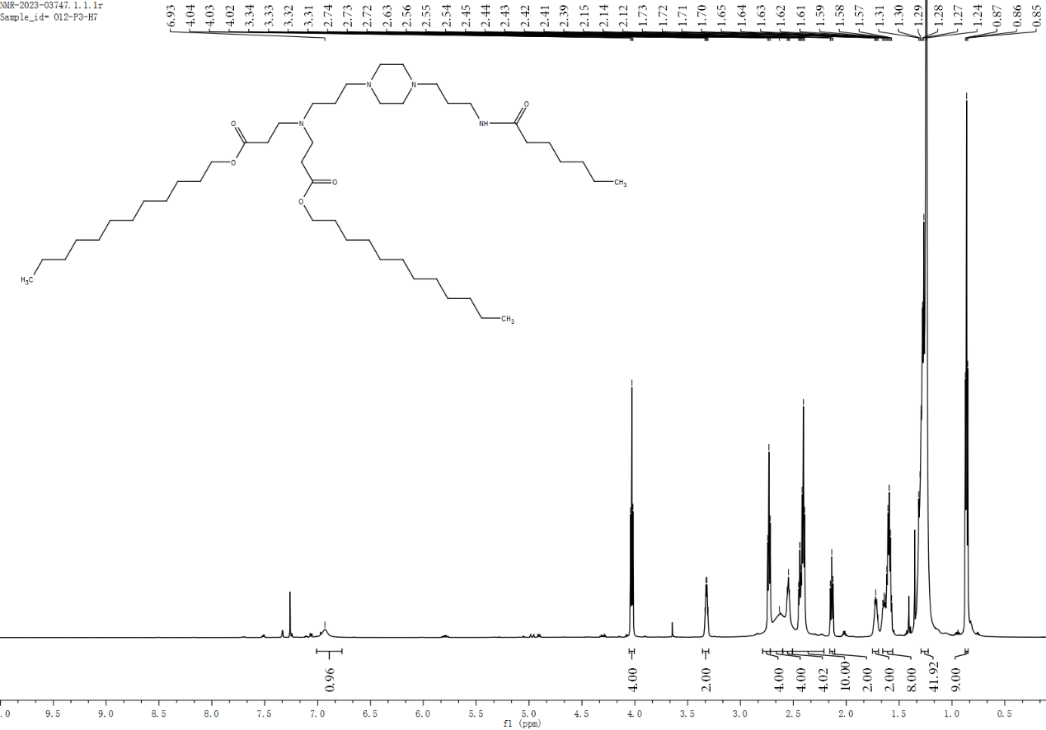

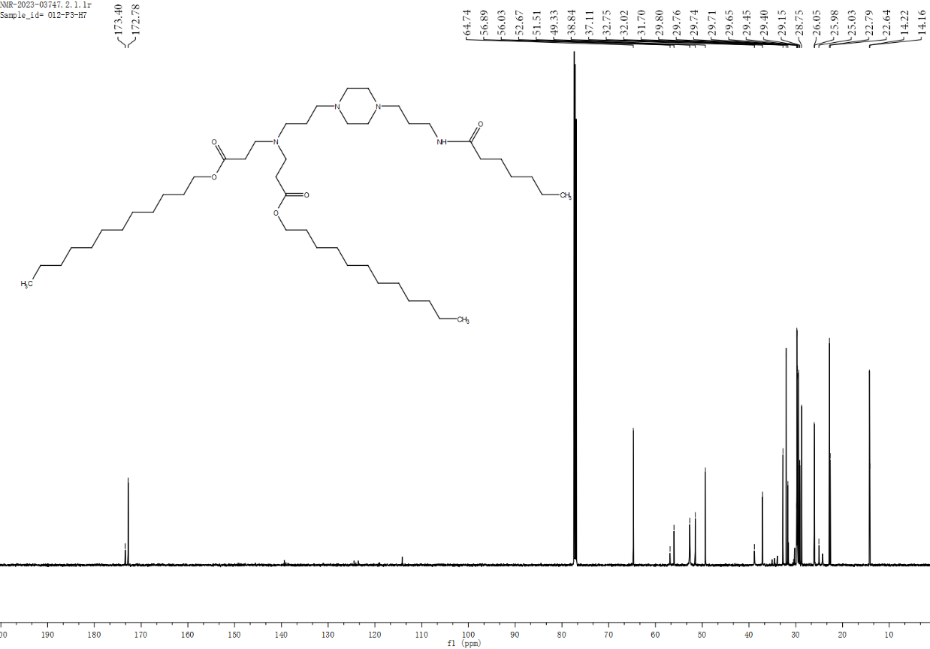

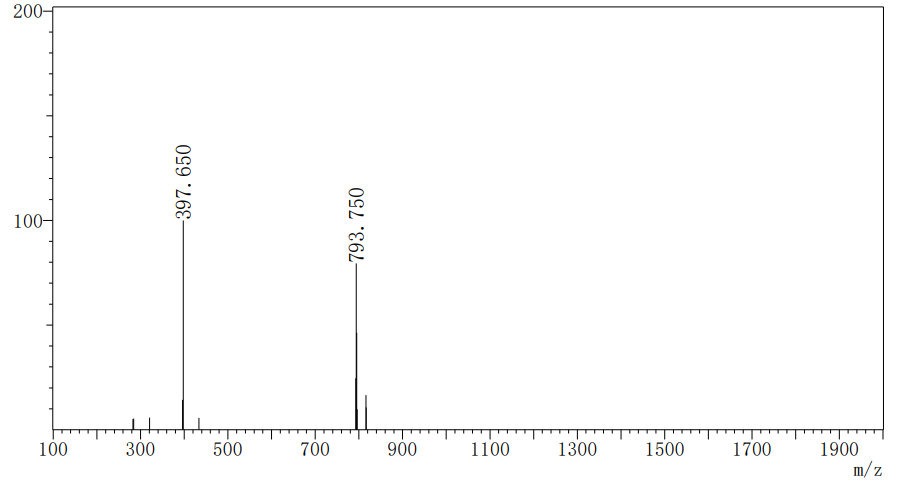

O12-P6-H8

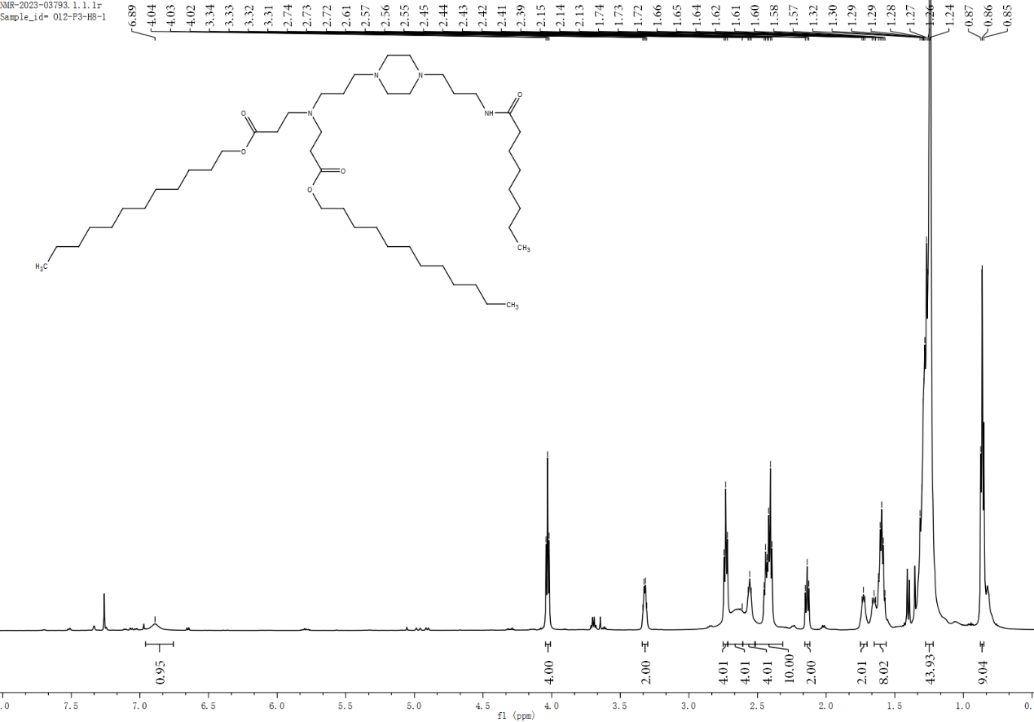

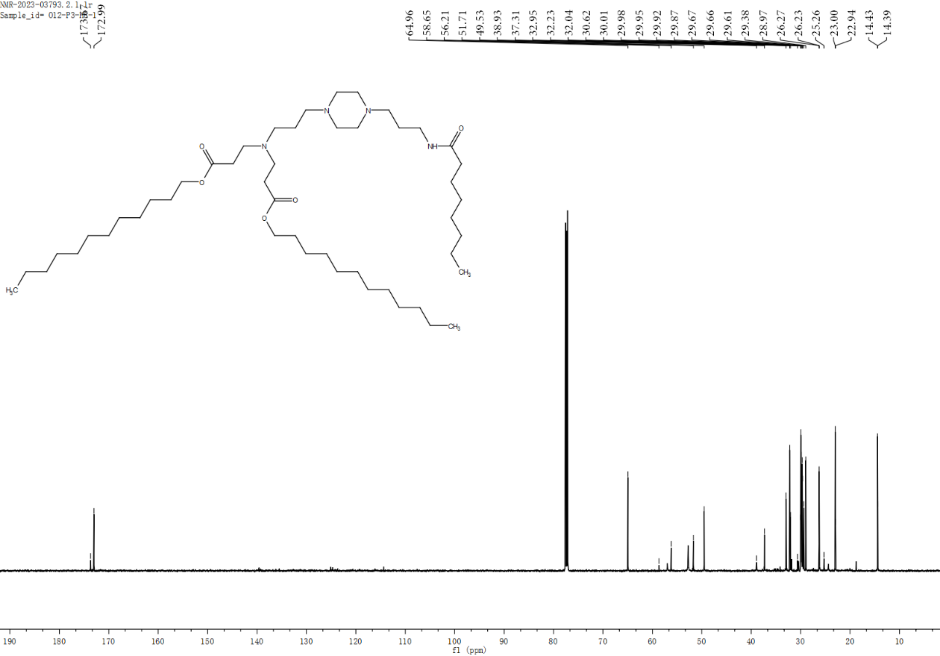

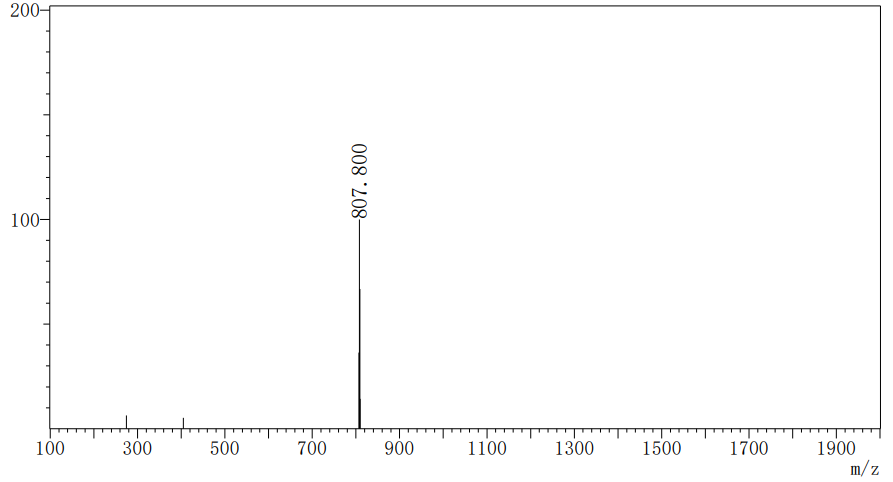

O12-P6-H9

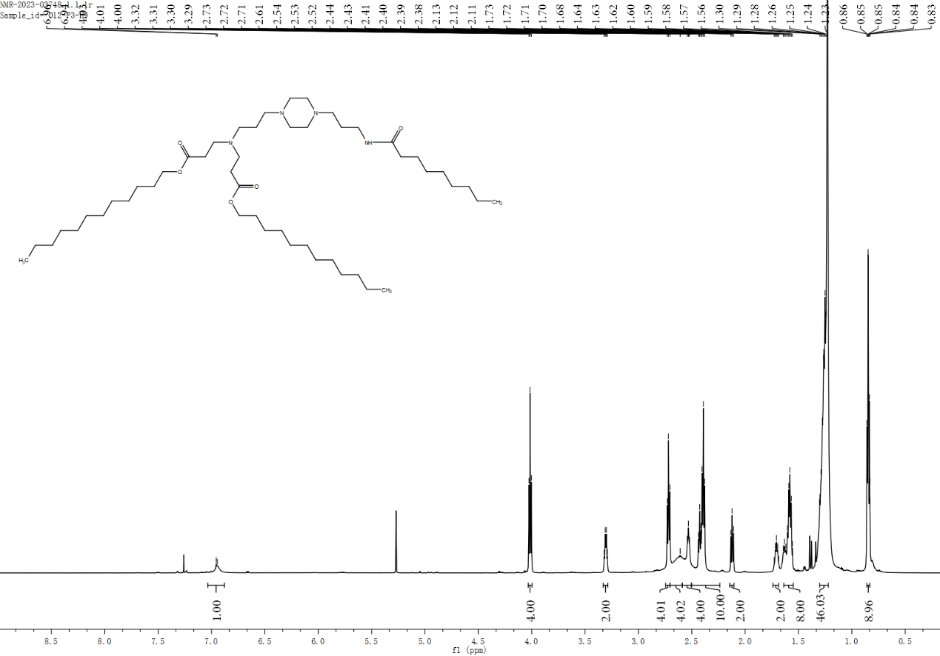

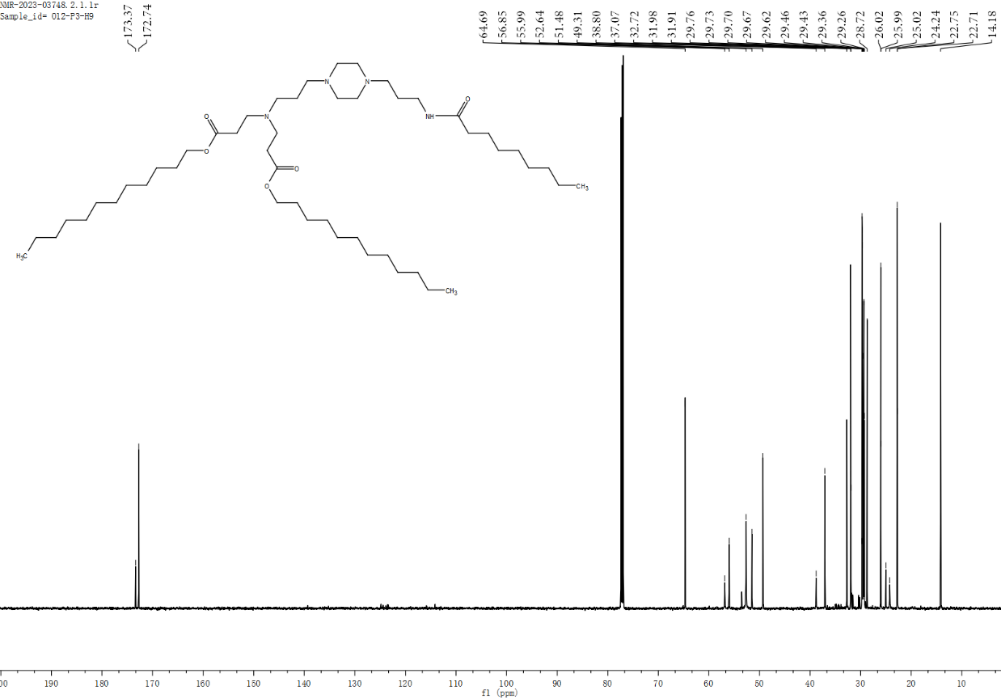

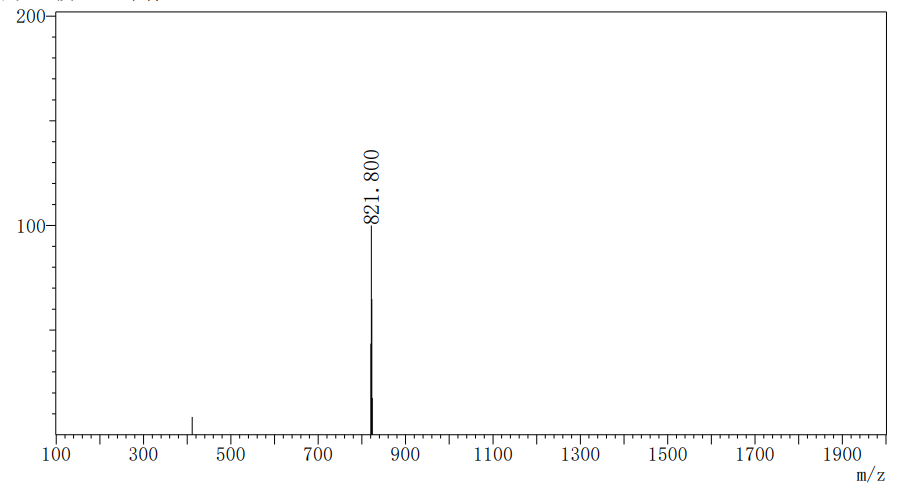

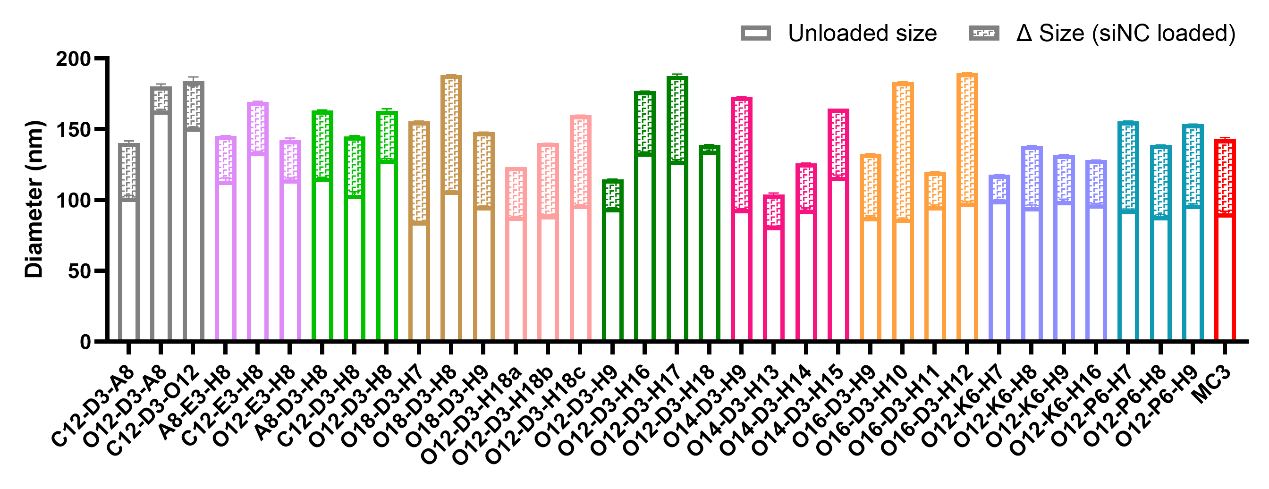

**FIGURE S1.** Particle size changes of LNPs before and after siNC encapsulation. The stacked bars show the unloaded size (solid color) and the size increment after siNC loading (patterned fill) for each LNP formulation. Data are presented as mean ± SD (n=3).

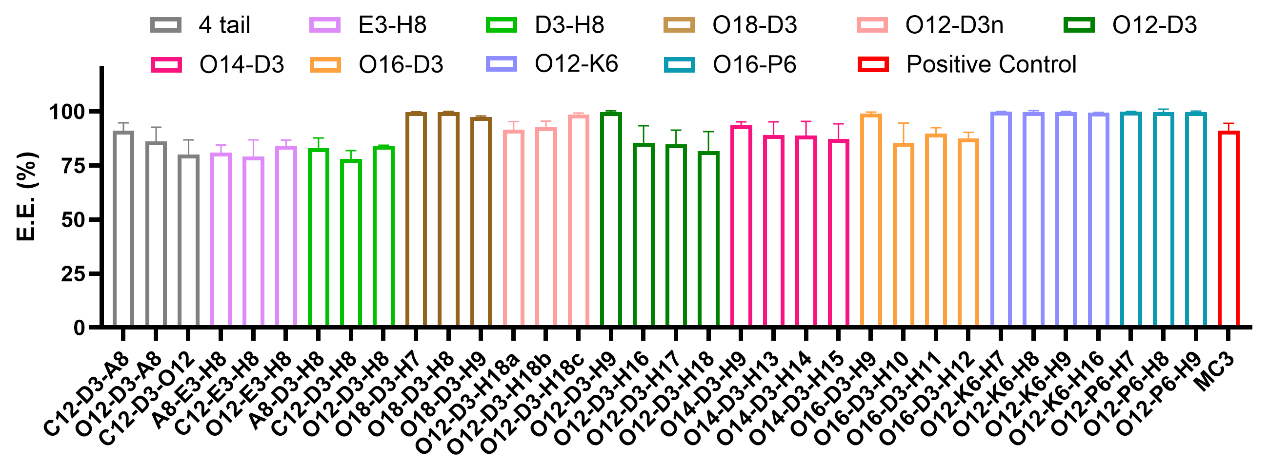

**FIGURE S2.** Encapsulation efficiency (E.E.) of siNC-loaded LNPs. Data are presented as mean ± SD (n=3).

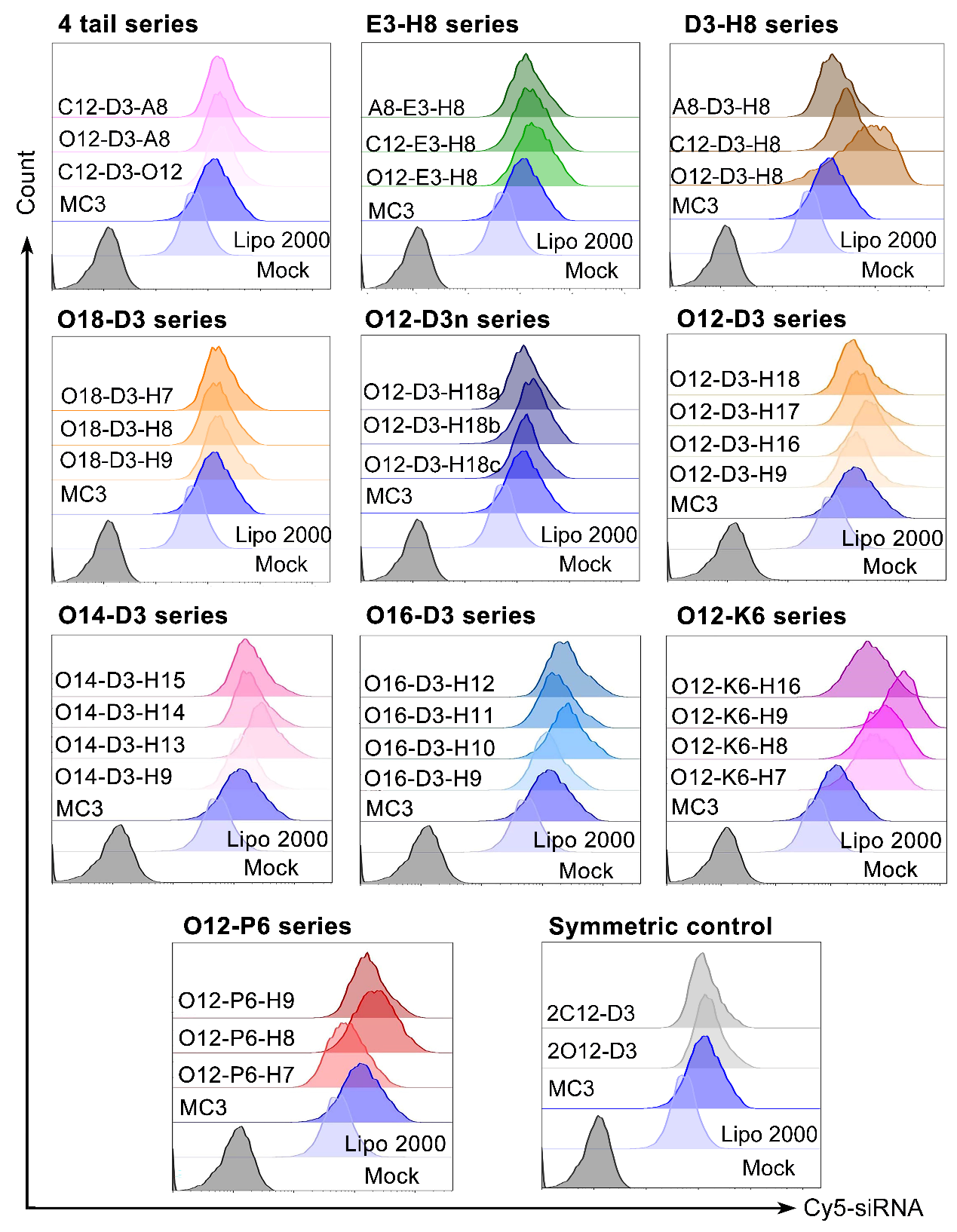

**FIGURE S3.** Overlaid flow cytometry histograms of intracellular Cy5 fluorescence intensity in HepG2 cells treated with different series of dissymmetric hydrophobic-tailed ionizable lipid-based LNPs.

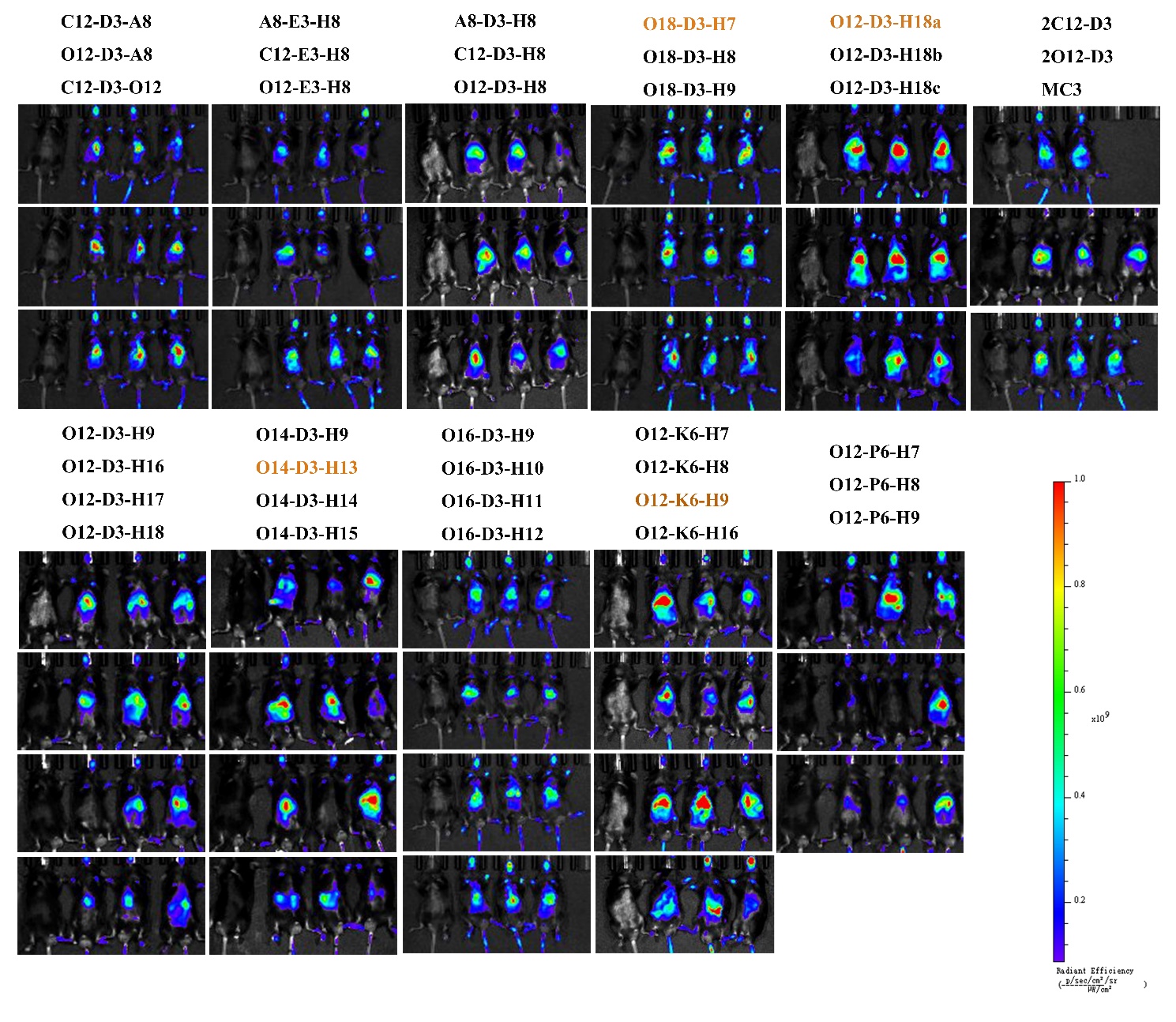

**FIGURE S4.** *In vivo* fluorescence imaging of 34 dissymmetric hydrophobic-tailed ionizable lipid-based LNPs in mice.

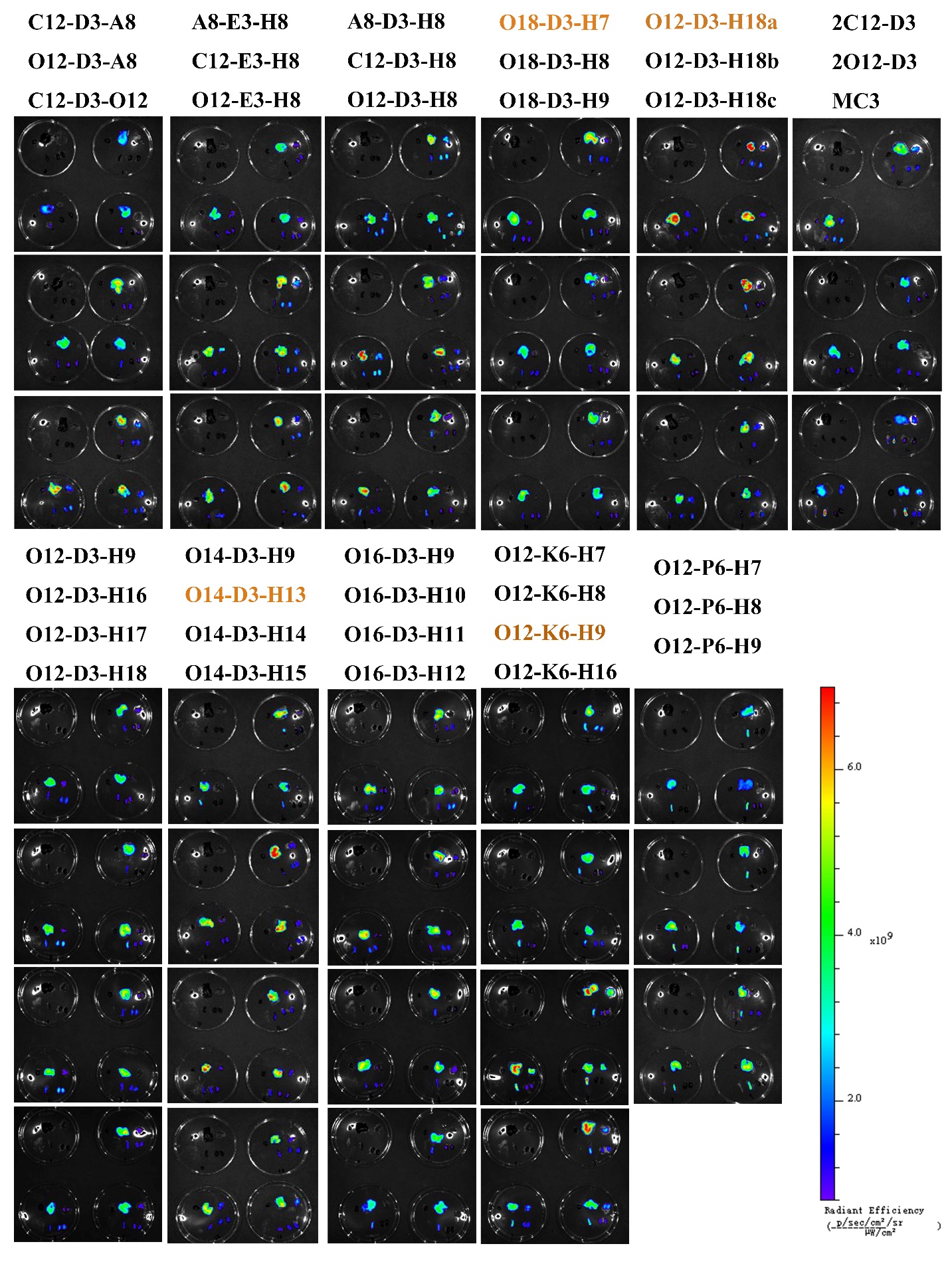

**FIGURE S5.** *Ex vivo* organ fluorescence imaging of 34 dissymmetric hydrophobic-tailed ionizable lipid-based LNPs in mice. Each group of culture dishes displays the organs in the following order: heart, liver, spleen in the first row from left to right, and lung, kidney in the second row from left to right.

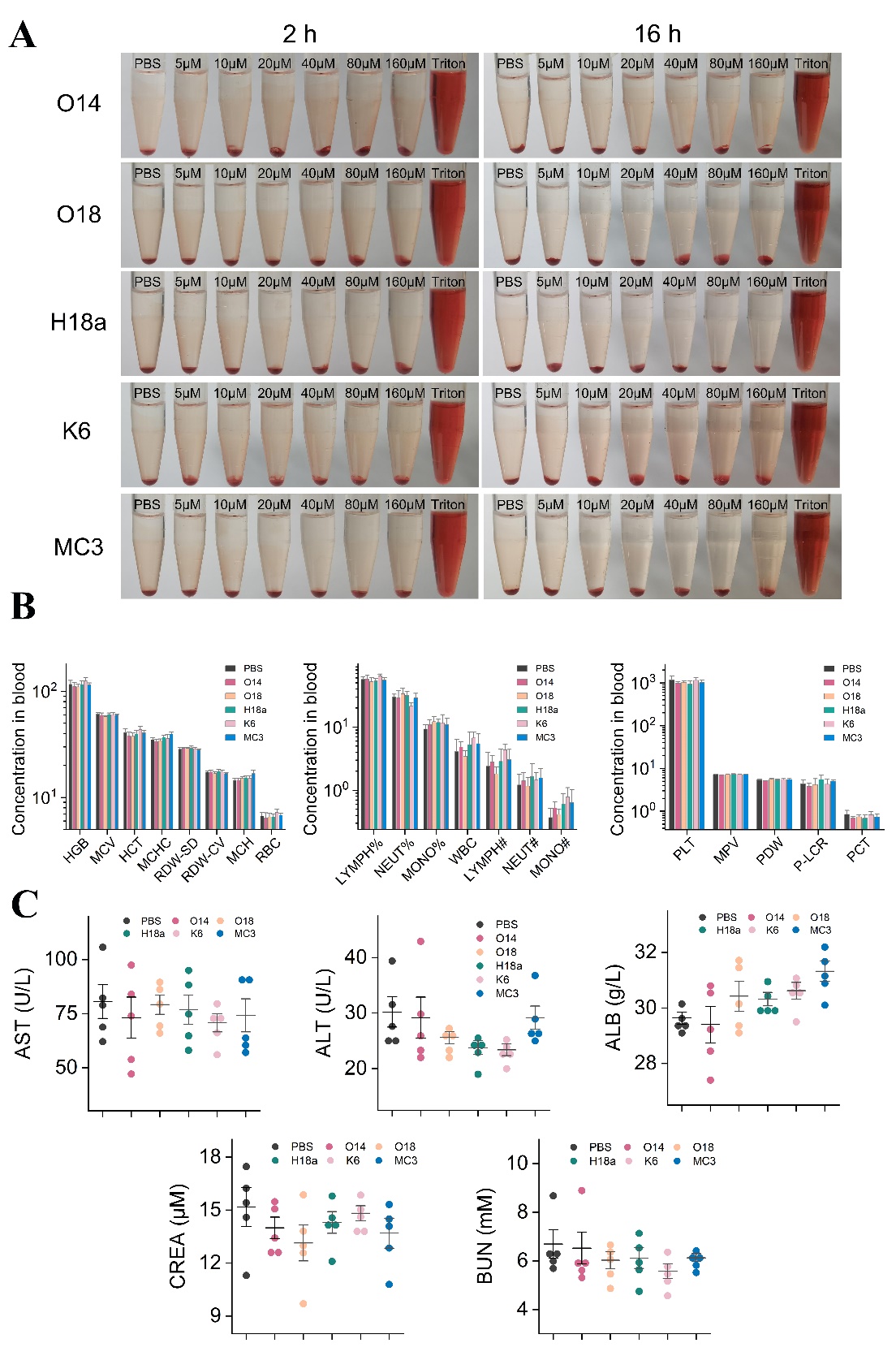

**FIGURE S6.** Supplementary safety evaluation data of candidate LNPs. (A) Representative digital photographs of centrifuged erythrocyte supernatants after incubation with LNPs for 2 h and 16 h, respectively. (n = 3) (B) Quantitative statistics of peripheral blood routine indexes collected from ICR mice. Detected parameters consist of erythrocyte-related indicators (HGB, MCV, HCT, MCHC, RDW-SD, RDW-CV and RBC), leukocyte percentage and absolute count indices (LYMPH%, NEUT%, MONO%, WBC, LYMPH#, NEUT#, MONO#), as well as platelet-associated markers (PLT, MPV, PDW, P-LCR and PCT). (n = 5) (C) Scatter plots of serum hepatic and renal function biomarkers (AST, ALT, ALB, CREA, BUN) from treated mice (n = 5). Data are presented as mean ± SD.

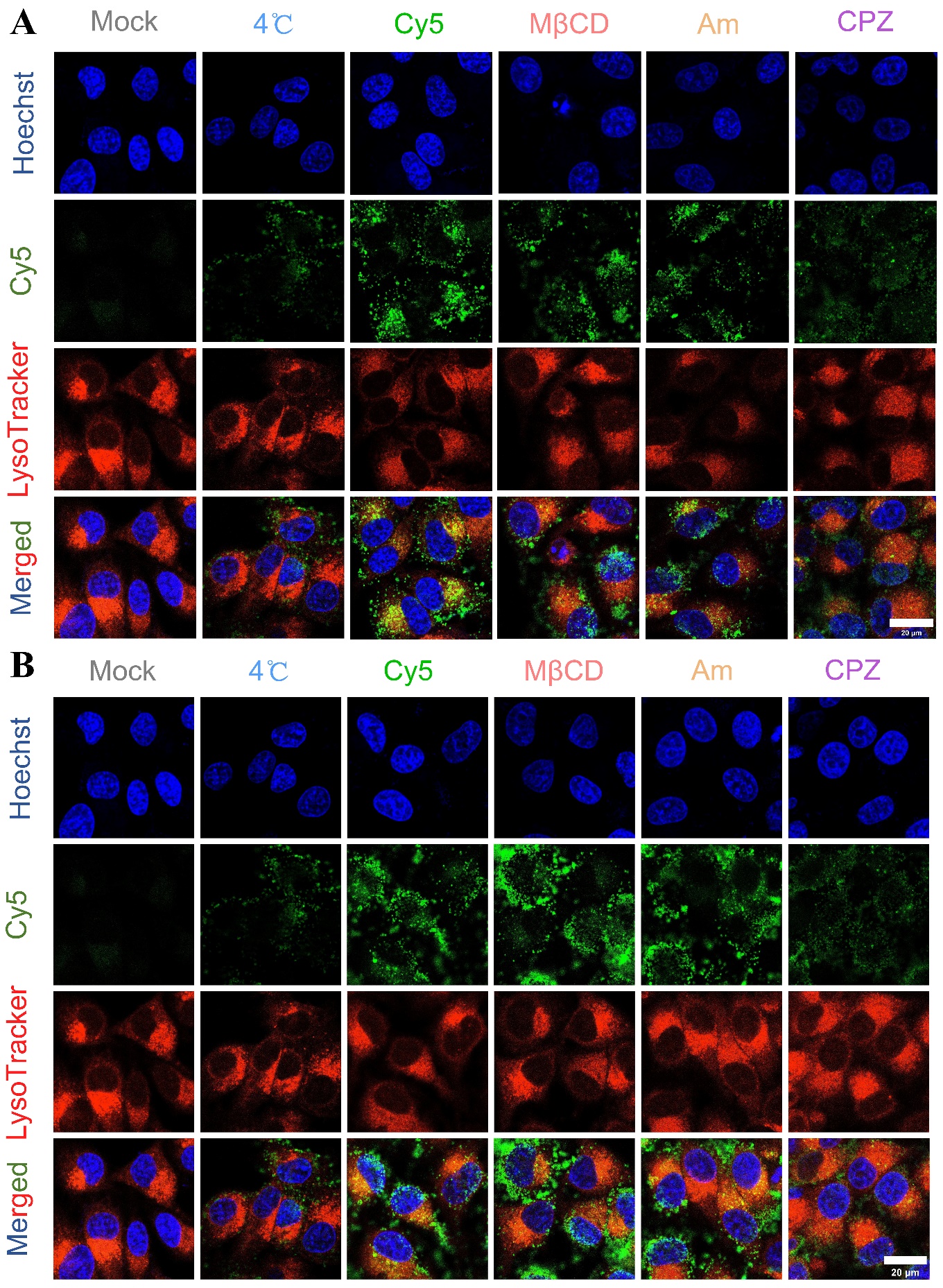

**FIGURE S7.** Representative confocal laser scanning microscopy (CLSM) micrographs of cellular internalization pathways for O14-LNP (A) and H18a-LNP (B). HepG2 cells were treated with Mock, 4 °C, untreated Cy5-LNP control, methyl-β-cyclodextrin (MβCD), amiloride (Am), and chlorpromazine (CPZ). Cy5-siRNA is shown in green, Hoechst-stained nuclei in blue, LysoTracker Red-labeled lysosomes in red. Scale bar = 20 μm.

**FIGURE S8.** Inhibitory effects of siALOX12-loaded LNPs on high glucose-induced lipid peroxidation in RAW 264.7 macrophages. (A) Representative laser confocal fluorescence images. Cell nuclei were counterstained with Hoechst 33342, and intracellular lipid peroxides were detected with the C11-BODIPY (581/591) probe. Scale bar = 10 μm. (B) Quantitative analysis of the mean fluorescence intensity (MFI) of oxidized C11-BODIPY. Data are presented as the mean ± SD from six independent experiments (n = 6), and ^###^*P* < 0.005, GLU group compared to the PBS group; ****P* < 0.005, **P* < 0.05, test groups compared to the GLU group.

**FIGURE S9.** Target validation and in vivo biocompatibility of siALOX12-loaded LNPs. (A) Relative mRNA expression of ALOX12 in liver tissues detected by qPCR. (B–F) Serum levels of hepatic and renal function biomarkers. (G) Routine blood test results, including erythrocyte parameters (left), leukocyte parameters (middle), and platelet parameters (right). (H) Representative H&E staining images of major organs (heart, lung, spleen, kidney) in each group. Scale bar = 200 μm. Data are presented as the mean ± SD from five independent experiments (n = 5), and ****P* < 0.005, test groups compared to the STZ group.

**FIGURE S10.** *In vivo* biocompatibility evaluation of siRNA-loaded H18a-LNP formulations. (A) Routine hematological parameters, including erythrocyte indices (left), leukocyte indices (middle), and platelet indices (right). (B) Serum renal function biomarkers: blood urea nitrogen (BUN, upper panel) and creatinine (CREA, lower panel). (C) Representative H&E-stained sections of major organs (heart, lung, spleen, kidney, liver). Scale bar = 200 μm. Data are presented as the mean ± SD from five independent experiments (n = 5), and ^##^*P* < 0.01, db/db group compared to the m/m group; **P* < 0.05, test groups compared to the db/db group.
